# Can Quantum Chemistry Improve the understanding of Protein-Ligand Interactions? Implications for Structure Based Drug Discovery

**DOI:** 10.1101/2023.06.01.543295

**Authors:** Filipe Menezes, Tony Fröhlich, Grzegorz M. Popowicz

**Affiliations:** Institute of Structural Biology, Helmholtz Zentrum München, Ingolstädter Landstr. 1, 85764 Neuherberg

**Keywords:** Energy Decomposition Analysis, Protein-Ligand Interactions, Interaction Maps, Semi-Empirical, ULYSSES

## Abstract

We introduce an Energy Decomposition Analysis suitable for understanding the nature of non-covalent binding in large chemical systems, like those of drug-protein complexes. The method is atom specific, thus allowing rationalization of the role that each atom or functional group plays for the interaction. Visual representations are constructed in the form of interaction maps, depicting the different contributions for electrostatics, polarization, dispersion (lipophilicity), *etc*. This marks the departure from atomistic models towards electronic interaction ones, that better correlate with experimental data. The maps provide a quick access to the driving forces behind the formation of intermolecular complexes, and the key contributors for each interaction. This allows constructing quantum mechanical models of binding. The presented method is validated against experimental binding data for the difficult to target protein-protein interface for PEX14-PEX5 and its inhibitors.

## INTRODUCTION

Structure based drug discovery aimed at placing protein-ligand interactions at the core of drug discovery.^1,2^ Despite the promise for increased rationale, the success rate for clinical drug development is still frustratingly low.^3,4^

Understanding which forces drive the formation of non-bonded complexes is essential for many fields of applied sciences, ranging from drug discovery to self-assembly and materials sciences. In drug discovery, alanine scan, structure-activity-relationship (SAR) data and scaffold hopping are methods probing experimentally the nature of the interactions between small ligands and their biological targets.^5^ The modifications explored with these techniques may however lead to unpredictable results. Availability of experimental structures seldom allows for rational optimization. Change of binding pose or of pocket renders quantification of the impact onto binding impossible. Crystal artefacts may also suggest wrong interpretations. All this leads to waste of resources and time. Solely relying on chemical intuition to understand the nature of intermolecular forces and the effects of key modifications is thus insufficient. More sophisticated techniques and methods are sought, which improve on the rationalization of interactions in (large) chemical systems.

Energy Decomposition Analysis (EDA) partitions interaction energies into several contributions, pinpointing the driving forces for forming non-covalent complexes. The first EDA was proposed by London,^6-8^ who categorized intermolecular interactions using quantum mechanical perturbation theory. London’s method was extended by many, but perhaps the most widely used method is Symmetry-Adapted Perturbation Theory (SAPT).^9-11^ Despite the very clear definition of what are electrostatics, dispersion, *etc*., energy partitioning is not unique, and other hallmarks were described.^12,13^ Common to these approaches is that they severely limit the system size, as they are based on wavefunction or density functional theories. Tackling molecules with thousands of atoms is impossible without drastic truncation, rendering the models unrealistic.^14–18^

Based on semi-empirical quantum chemistry, van der Vaart and Merz proposed a simple EDA technique that addresses large chemical systems of arbitrary size.^5,19-21^ However, their method only splits interaction energies into electrostatics, polarization, and charge transfer. Electronic density repulsion is not quantified, though it contaminates electrostatics and polarization. Dispersion forces are also absent from the treatment.

Another limitation of most EDA algorithms is the strict focus on total energies. A single value describes electrostatics, another expresses polarization, *etc*. An atom-specific partitioning would prove crucial, particularly in inhibitor design. Not only this specifies the molecular fragments dominating certain interactions, but it also facilitates the rationalization of trends. Again, Merz and coworkers were pioneer when they proposed an atom-specific partitioning.^5^ Their focus was however on the ligand-residue interactivity. The atom-atom partitioning of SAPT by Parrish and Sherrill is also noteworthy.^11^

Here, we report a fully atom-specific EDA method that quantifies all relevant contributions to the interaction energy: electrostatics, polarization, charge transfer, electronic density repulsion, lipophilicity (dispersion) and implicit solvation. The atom-specific partitioning is fully flexible, permitting focus on the whole complex, on residue-ligand interactions, or even between functional groups. This is based on interaction matrices, which can be easily explored by machine learning techniques to describe (bio)molecular binding. Interaction maps (*imaps*), intuitively interpreted by non-experts, are assembled from those matrices, easing rational ligand optimization. These offer orthogonal information, not directly retrievable by direct structural analysis. Our EDA method allowed us to explain the SAR data of antiparasitic PEX14-PEX5 inhibitors as a proof of concept, by generating a complex quantal binding selectivity model.^22,23^

## RESULTS

The binding constant characterizes the strength of interactions between two molecules, *e*.*g*., a protein and an inhibitor. Via their connection with affinities, binding constants relate to molecular models. When fine tuning an inhibitor, relative affinities typically suffice. For interactions of similar nature,^24^ binding entropies become quasi-constant and may be excluded altogether: drug discovery may focus exclusively on enthalpies of binding. Here is where EDA’s role is critical, by allowing one to understand the nature of the interactions in biomolecular complexes. Ligand optimization decisions become physically motivated, instead of intuition based.

Our EDA program determines 7/8 components that characterize interaction energies. Electrostatics (ES) measure the mean-field Coulomb interaction between molecules.^25^ This stabilization/repulsion that results from representing atoms as point charges is obtained from the electronic densities of the isolated entities.^20^ Polarization (POL)^20^ characterizes the intramolecular flow of electronic density due to an interaction partner. It is calculated using constrained Self-Consistent-Field^26,27^ to optimize an electronic state in which interaction partners do not formally exchange electrons. Charge transfer (CT) also describes a flow of electrons,^20^ however, this flow takes now place between different molecules. Electronic density repulsion (REP) evaluates the increase in energy due to overlapping electronic densities, a consequence of the repulsion electrons experience from each other. If fragments come too close to one another, overlap-repulsion grows exponentially.^17^ We also calculate the contribution of dispersion (DISP),^6-8^ which reflects the correlation in electronic motion. Electronic distributions around nuclei are non-static and instantaneous charge fluctuations are constantly generated. Though present in all chemical systems, this interaction dominates in apolar molecules. It therefore correlates with lipophilicty.^28^ The last term that may be obtained in our EDA is solvation (SOLV), if an implicit solvation model is requested. This measures the relative stability of all species involved with respect to each other and to a hypothetical gas phase (S17). When summed, all contributions yield the binding or interaction energy (INT).

Chemical biology considers π-stacking as an important interaction. Though it is still under debate,^29^ our EDA sees π-stacking as the DISP forces from interacting π-clouds. Hydrogen-bonds are decomposed as a combination of ES and CT contributions.^30^ The energy partitioning is designed to be fully compatible with semi-empirical quantum chemical methods,^31-34^ which renders calculations on protein-ligand complexes affordable on desktop computers.

We decided to use the PEX14-PEX5 interface as a model system, a novel antiparasitic target.^22,23^ However, the method is applicable to nearly any non-bonded complex, not necessarily of biological interest. PEX14 binds PEX5 in a shallow lipophilic interface exacerbating the influence of solvent mediated interactions (Figure 1). This presents a challenging medicinal chemistry target. We chose complexes of PEX14 with inhibitors – 5L87, 5L8A, 5N8V, 5OML, 6RT2, 6SPT – and with peptides – 2W84, 2W85, and 4BXU.^23,35-46^ A modified variant of 2W84 was prepared, where a key aspartate is converted into a glycine (2W84’).

**Figure 1.**
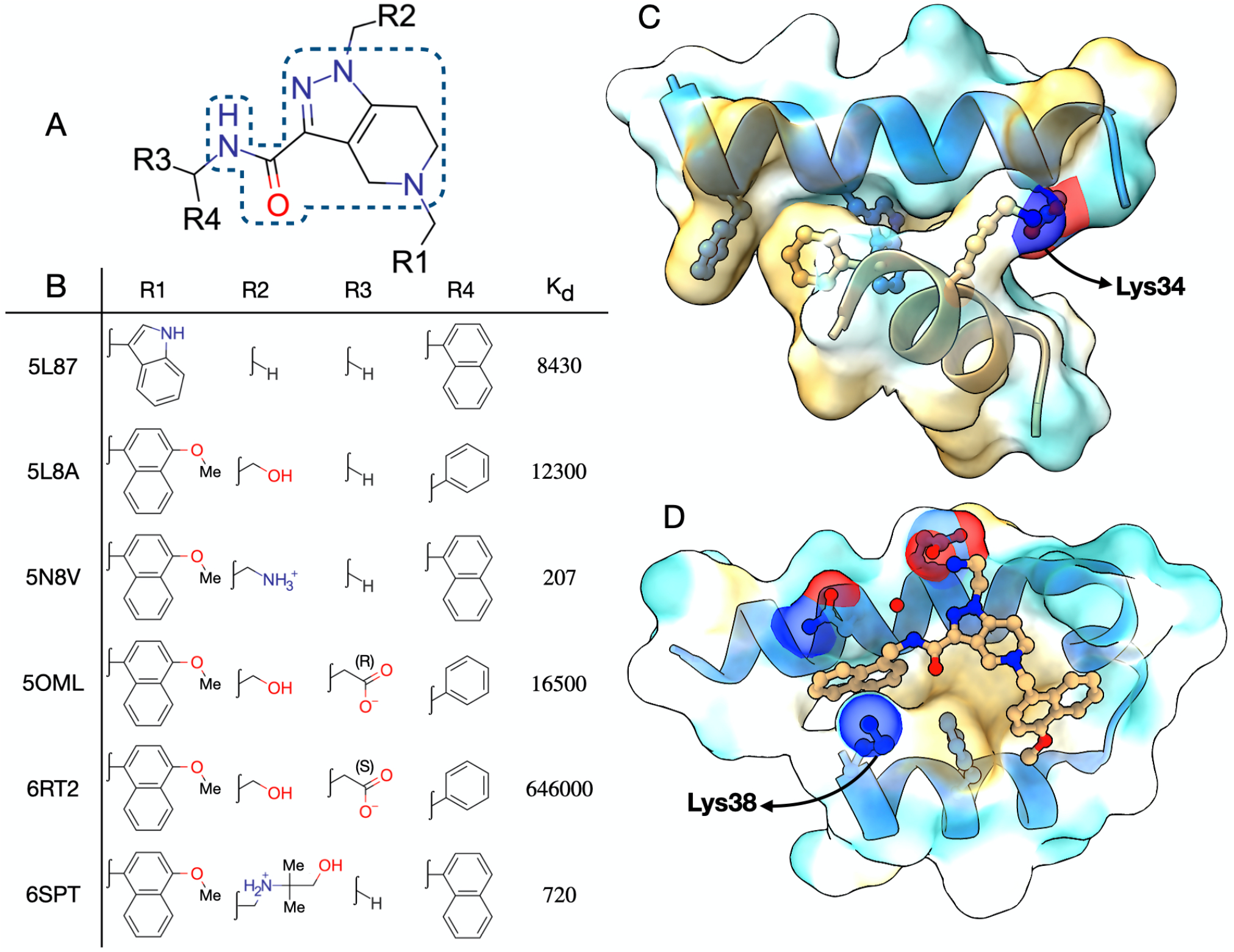
A) Structures of the ligands used to validate the EDA method. The tetrahydro-pyrazolo-pyridine-carboxamide scaffold demarked within the dashed area. B) Summary of SAR data on several PEX14 inhibitors collected from the literature, the K_d_s are in nM.^23,35-41^ C) Part of PEX14 structure with a small peptide, 2W84,^42,43^ showing the target protein-protein interface. The blue ribbon and atoms are the small peptide (up), while the brown ribbon and atoms represent PEX14 (down). D) PEX14 with a strong, small molecule binder, 58NV.^23,37^ C and D are both clipped to the essential interaction surfaces.

By construction, *imaps* are normalized to the strongest contribution of each force, and different *imaps* should not be directly compared. Reference points may however be easily replaced.

When comparing SAR with simulation data, protonation states cannot be circumvented. Since binding affinity assays employed conditions identical to those for crystallization, a total charge of +5 was used for PEX14. This was determined from the structures in PDB.

### A PHYSICAL BASIS TO SAR

In all cases analyzed, ES maps show the conservation of the role of the tetrahydro-pyrazolo-pyridine-carboxamide core (Figures 2 and 3; S12). Indole’s nitrogen in 5L87 is seen mimicking the electrostatic behavior of the methoxy group in all other ligands. This indicates that EDA may be used to define non-classical bioisosterism. Differences arise however in the groups in positions R2 and R3: neutral polar groups, like alcohols, show overall attractive interactions; a methyl substituent results in weak repulsion; an ammonium, leads to strong repulsion. Note that calculations weight local and global protein-ligand interactions.

**Figure 2.**
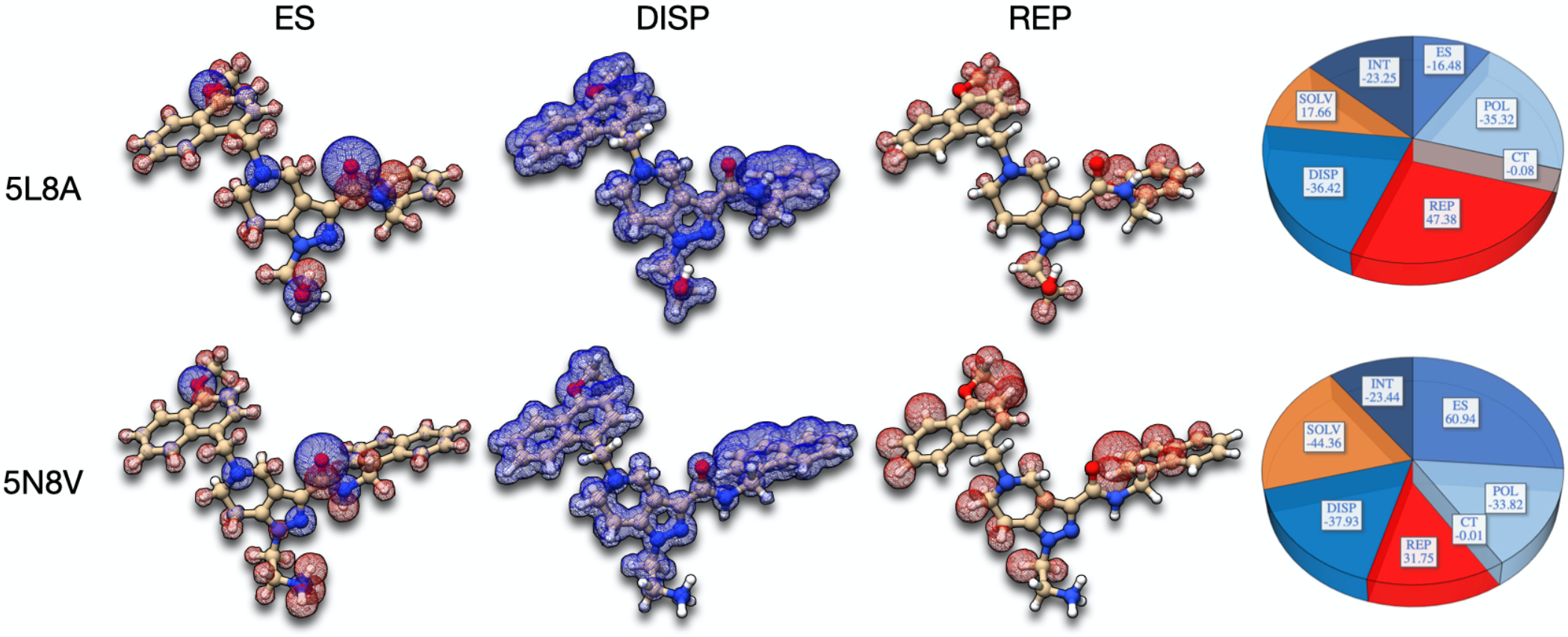
EDA representations of binding energy contributions for chemically similar molecules. A selection of interaction maps for 5L8A,^23,36^ a weak binder, and for 5N8V,^23,37^ a strong binder. Blue represents attractive interactions, whereas red stands for repulsion. Pie plots provide information on the EDA for these two complexes. Energies therein reported are in units of kcal/mol.

**Figure 3.**
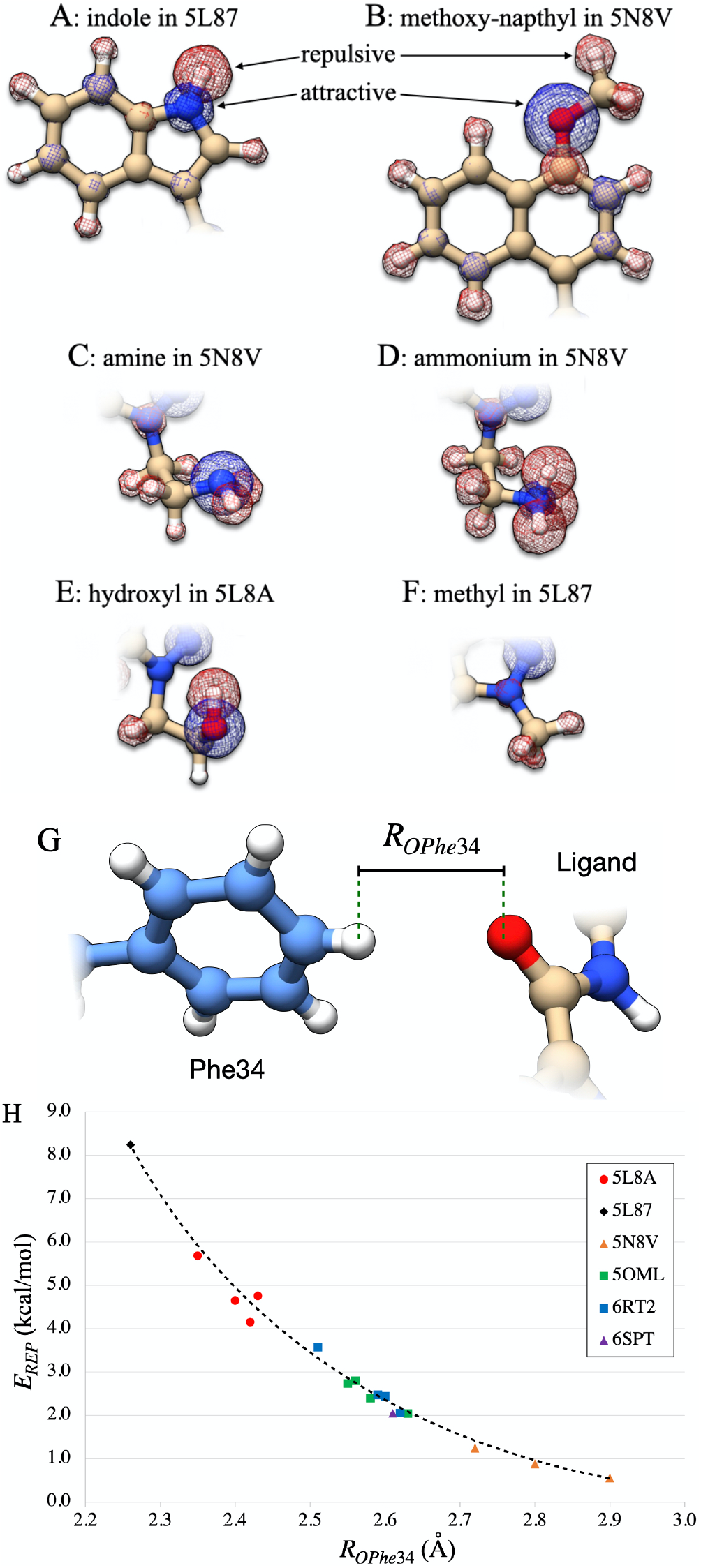
A) - D) ES contributions for different functional groups in the ligands. G) – H) Electronic Density Repulsion (kcal/mol) against the shortest distance between the ligand’s carboxamide’s oxygen and PEX14 (R_OPhe34_, in Å). Different ligands are marked with different colors and shapes, so that each point is easily associated with one protein-ligand complex (compare Figure 1).

Water’s solvation effect is predominantly electrostatic. Solvation is penalizing whenever the electrostatic surfaces, available to interact with the solvent, decrease. On the other hand, when the total charge increases upon complexation, solvation becomes favorable. Most cases show repulsive solvation terms, attesting the electrostatic nature of protein-ligand interactions. CT is in all cases negligible, contrasting with previous findings.^21,47^ This negligible stabilization seems to be particular of some PEX14 crystal structures. Later we demonstrate that dynamical relaxation of structures can strengthen CT.

Dispersion maps show minor to absent contribution from polar groups, bespeaking the parallelism with lipophilicity. It is readily observed that most atoms contribute to lipophilic interactions. The few exceptions are immediately identified with the *imaps*. There is also higher lipophilic density on atoms closer to the protein’s pocket. Alone, they suffice to identify groups in the ligand going into the protein’s pocket (Figure S12.7).

Since REP terms decay exponentially with the interatomic distance,^17^ these maps are sensible to fine structural details in binding. This pointed us to subtle, but systematic, differences in the REP contribution of the ligand’s carboxamide, which was furthermore correlated with the electrostatic prowess of groups in R2 and R3 (Figure 3H). When hydrogen-bonds and ionic-bridges are possible, the carboxamide, more than 7 Å apart from those groups, is shifted. Interestingly, the isobutanol-extended ammonium in 6SPT’s R2 substituent places this ligand between 5L8A (hydroxyl only) and 5N8V (ammonium only). We stress that these observations, resulting from extremely fine structural details, are barely noticeable without EDA.

### MIMICKING PROTEIN-PEPTIDE INTERACTIONS

Protein-peptide systems show strong variations in ES and SOLV contributions, not only within themselves, but also with respect to protein-ligand complexes. These are easily understood given the increased net charge of complexes and fragments and the variety of charge distributions. The dispersion stabilization is conserved among all complexes thus far studied (Figure 4; S13). The main outlier is a complex of PEX14-PEX5 on a different pocket. In 2W84, PEX5 points towards PEX14 an aspartate (Asp96) that forms a salt-bridge and a hydrogen-bond with a nearby lysin (Lys34; Lys38 in complexes with inhibitors). *In silico* mutation of Asp96 into a glycine -2W84’ - decreases binding by almost 3 kcal/mol and completely alters the landscape of interactions. Lipophilicity, however, remains unaffected. This further indicates that the PEX14-PEX5 interaction is dominated by local and directed electrostatics.

**Figure 4.**
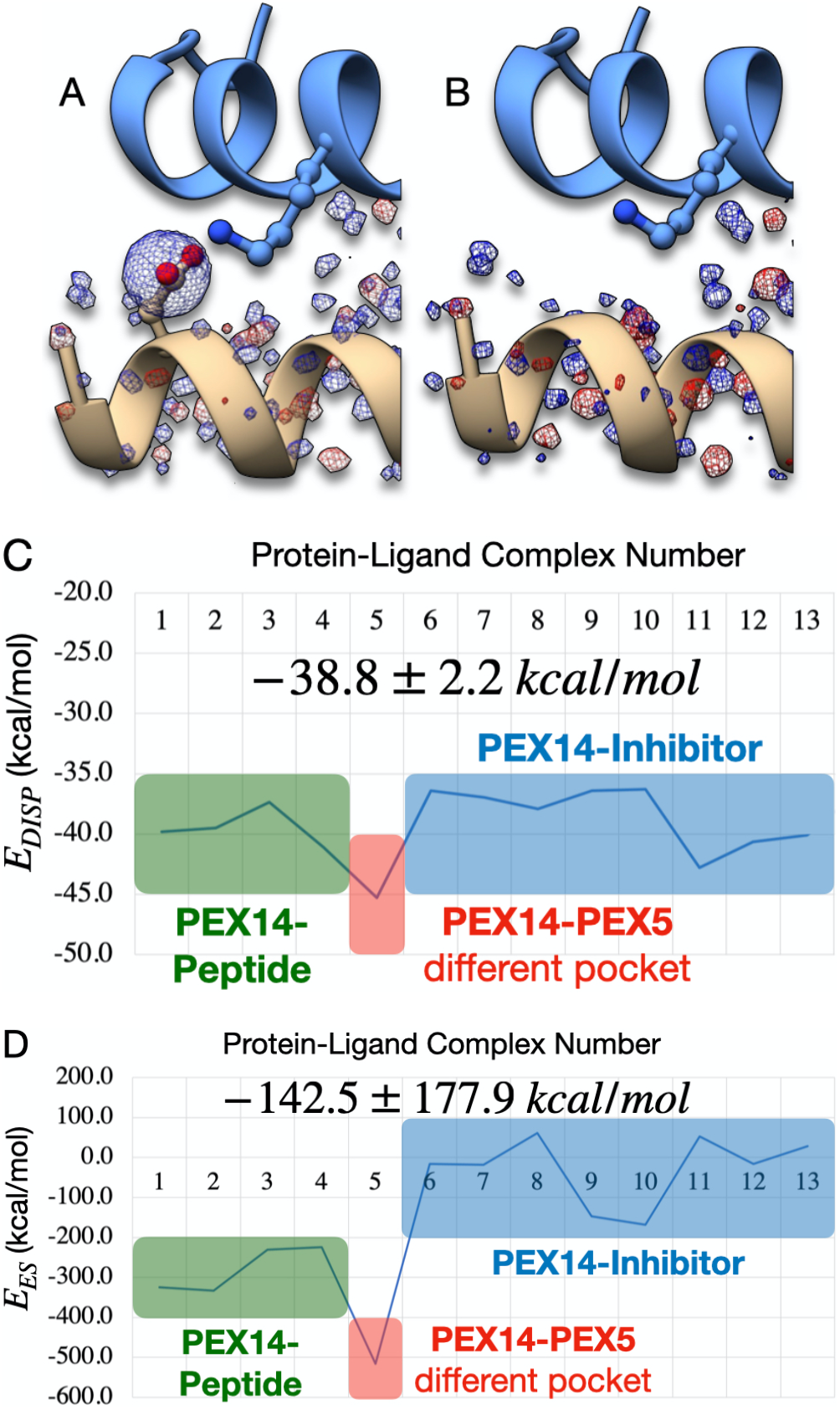
Magnification of the total interaction maps (INT) of A) PEX5 with PEX14 (2W84) and B) modified PEX5 and PEX14 (2W84’) around the residue Asp96. C) The dispersion and D) the electrostatic interaction energy between PEX14 and its ligands, peptides, and inhibitors. Dispersion (lipophilic) interactions in protein-ligand and protein-peptide complexes are remarkably identical and uniform, unlike electrostatics, which are significantly stronger for peptides and highly variable.

Comparing the PEX14-PEX5 and PEX14-PEX19 complexes we see that shape complementarity is better in the former (larger REP). Lipophilicity is slightly larger for PEX19, partly because of the 3 aromatic rings that PEX19 places in PEX14’s pocket. The calculations indicate however the enthalpic selectivity for PEX5 is due to electrostatics. This goes along the picture assembled in the previous paragraphs.

### THE EFFECTS OF WATER AND COUNTER IONS

By treating solvation waters and/or ions as integral part of the protein, their influence on binding is investigated (Figure 5; S16). Explicit waters impact most quantities, except lipophilicity. Interaction energies stabilize by up to 8 kcal/mol. The CT and INT maps gain definition, though the total weight of the former is unaffected. Electrostatics evidence the role of the amide. REP terms accentuate the contribution of solvent for protein-ligand shape complementarity, as water is key for binding onto the PEX14 surface.^22^ The addition of a chloride ion changes pronouncedly solvation and electrostatics. However, the final binding energy is minimally impacted. The maps also show minor sensitivity to the presence of the anion. Nonetheless, including the chloride is essential for more accurate descriptions of the underlying forces that lead to binding.

**Figure 5.**
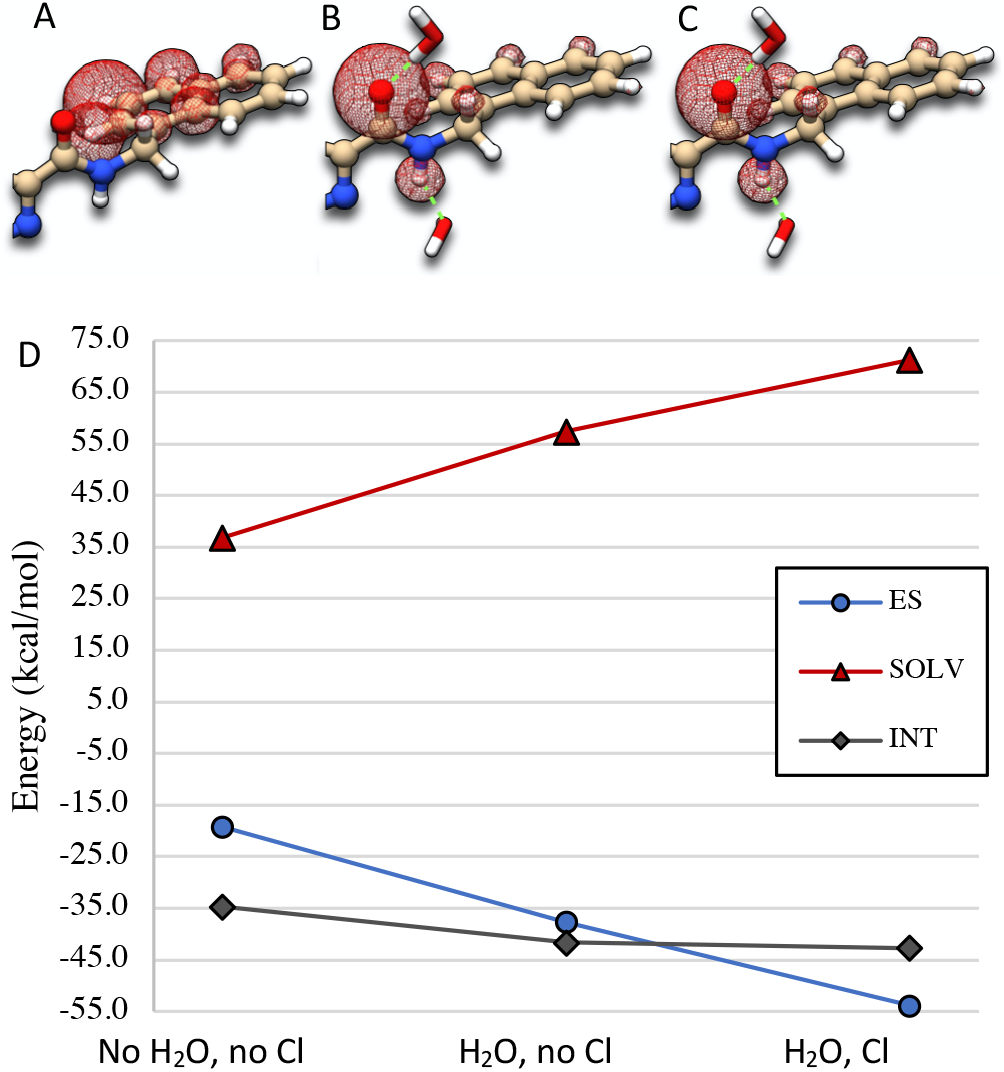
The effect of explicit water and of a chloride anion on the electrostatics, solvation, and total binding of 5N8V. A) the REP maps without explicit waters or counter ions, B) with explicit waters but no counter ion and C), with explicit waters and counter ions. D) Evolution of the electrostatics, solvation, and interaction energies as a function of the conditions.

From the analysis above we conclude that explicit water contributes to the intensification of protein-ligand interactions. However, water does not affect the nature of interactions, not even the salt-bridges or hydrogen-bonds. Water acts instead on assisting already existing protein-ligand interactions, not in the generation of new contacts. We note that, in general, explicit solvation is important, *e*.*g*., for the correct assignment of ligand protonation states (S14). The possibility to include large explicit solvation shells is a significant advantage of our algorithm.

### DYNAMICAL EFFECTS

50 ns long molecular dynamics (MD) were run on the ammonium and amine variants of 5N8V, and EDA was recorded every 10 ns (Figure 6). The first prominent observation is that the binding energy shows minimal oscillations around average values, indicating no special event. But the decomposed energies evidence otherwise: the electrostatics for the ammonium-ligand show a maximum at 10 ns, corresponding to a reorganization of the hydrogen-bond network as supported by CT. Then the system is mainly driven by electrostatics. Solvation also reflects the formation of electrostatic contacts between protein and ligand, and REP indicates that these contribute to maximization of shape complementarity.

**Figure 6.**
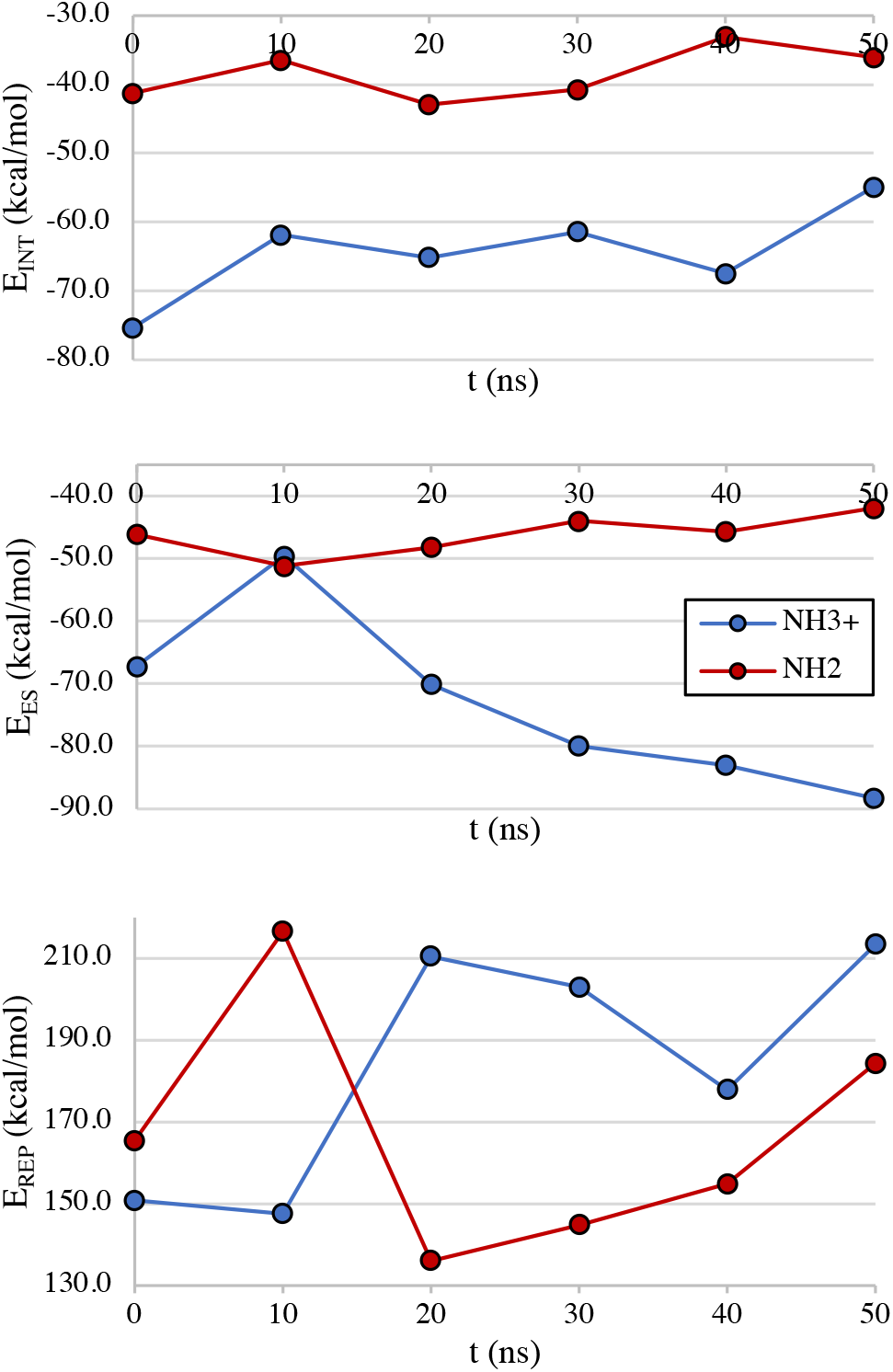
The evolution of some EDA contributions as a function of time.

It is noteworthy that electrostatics are clearly attractive in both protein-ligand complexes and even more favorable for the ammonium-ligand. This results from the structural relaxation by MD and the optimized hydrogen-bond network of 4 Å of solvation sphere (270 waters). The improved shape complementarity in the ammonium-ligand is interpretable based on strengthened electrostatics. The SI contains further MD traces (S16, S19). Increased CT contributions are observed. Interestingly, this is not the case for the acid-form of 5N8V, as CT reduces the electrostatic prowess.

### CRYSTAL ARTEFACTS

Binding energies for protein-ligand dimers in the asymmetric unit are quite heterogeneous, indicating crystal artefacts in several subunits (S18). This is the case of 5OML and 5L8A, where ligand driven π-stacking is particularly strong. Unsurprisingly, dimerization is lipophilicity-driven, as indicated by linear regression of dimerization energies with respect to dispersion. Key hydrogen-bonds are also seen in these structures, which account for a quasi-linear dependence on electrostatic interactions. The best correlation between dimer stability and electrostatics is however seen for 5N8V. In fact, electrostatics-related terms are the ones showing any possible correlation in this series of dimers, which is traced back to specific salt bridges established between units.

## DISCUSSION

In developing potent inhibitors, careful attention should be given to lipophilicity. This is because the inhibitor must find its way across membranes to the PEX14’s surface. Yet, overly lipophilic ligands have suboptimal drug-likeness and low selectivity. EDA calculations indicate that the lipophilic prowess of a series of inhibitors developed in our laboratory mimics the natural biological target: the dispersion interactions in PEX14-ligand and PEX14-PEX5 complexes all fall in the same range. Reduced polarizations (molecular-polarization/number-atoms) further reinforce this observation, since model peptides and inhibitors all fall in the same range. But if lipophilicity is not properly counterbalanced by electrostatics, an overly lipophilic surface is created for the complex. *In vitro*, this is translated into lipophilicity-driven aggregation. In biological conditions, several scenarios may be hypothesized. For instance, unspecific binding. EDA results on the PEX14-PEX5 complex reveal however that dissociation does not occur because of local salt-bridges coupled with hydrogen-bonding (electrostatics). Were these to be suppressed, binding would be critically impaired (ΔΔ*E*_*bind*_ ≈ 3*kcal*/*mol*). Studies performed in implicit wet octanol, a less electrostatic medium, further emphasize the role of electrostatics. EDA calculations on MD traces reinforce this picture even further, as the dynamics of protein-ligand complexes are electrostatics-driven. Here, EDA clearly correlates with classical principles of drug discovery where electrostatics and lipophilicity must be optimally balanced. Interestingly, in the 5N8V case, protein and ligand are both positively charged. Still, ES interactions are the main driving force for binding. This can only be rationalized given the local nature of the interactions identified. In accord with classical principles, EDA shows that exploiting electrostatics upon achieving sufficient lipophilicity is the fundamental principle in optimization campaigns.

However, these calculations reveal that electrostatics have a “non-linear” influence on binding pose and shape complementarity of the molecular surfaces. The EDA study of repulsion makes such statement clear. Aromatic groups in the ligands create two anchoring points to PEX14, responsible for the lipophilicity prowess. A third anchoring point originates from electrostatics. If improperly placed, ES does not improve shape complementarity, even with suitable interactions (Figure 7). Consequently, weak binders result. Moreover, favorable distributions of anchoring points lead to powerful inhibitors, irrespective of the ligand total charge. This is striking in the case of 5OML against 5N8V. Despite directional salt-bridges and hydrogen-bonds, only in 5OML protein and ligand have opposite total charges. 5N8V sets, however, anchoring points that improve shape complementarity, as supported by EDA calculations and experimental data.

**Figure 7.**
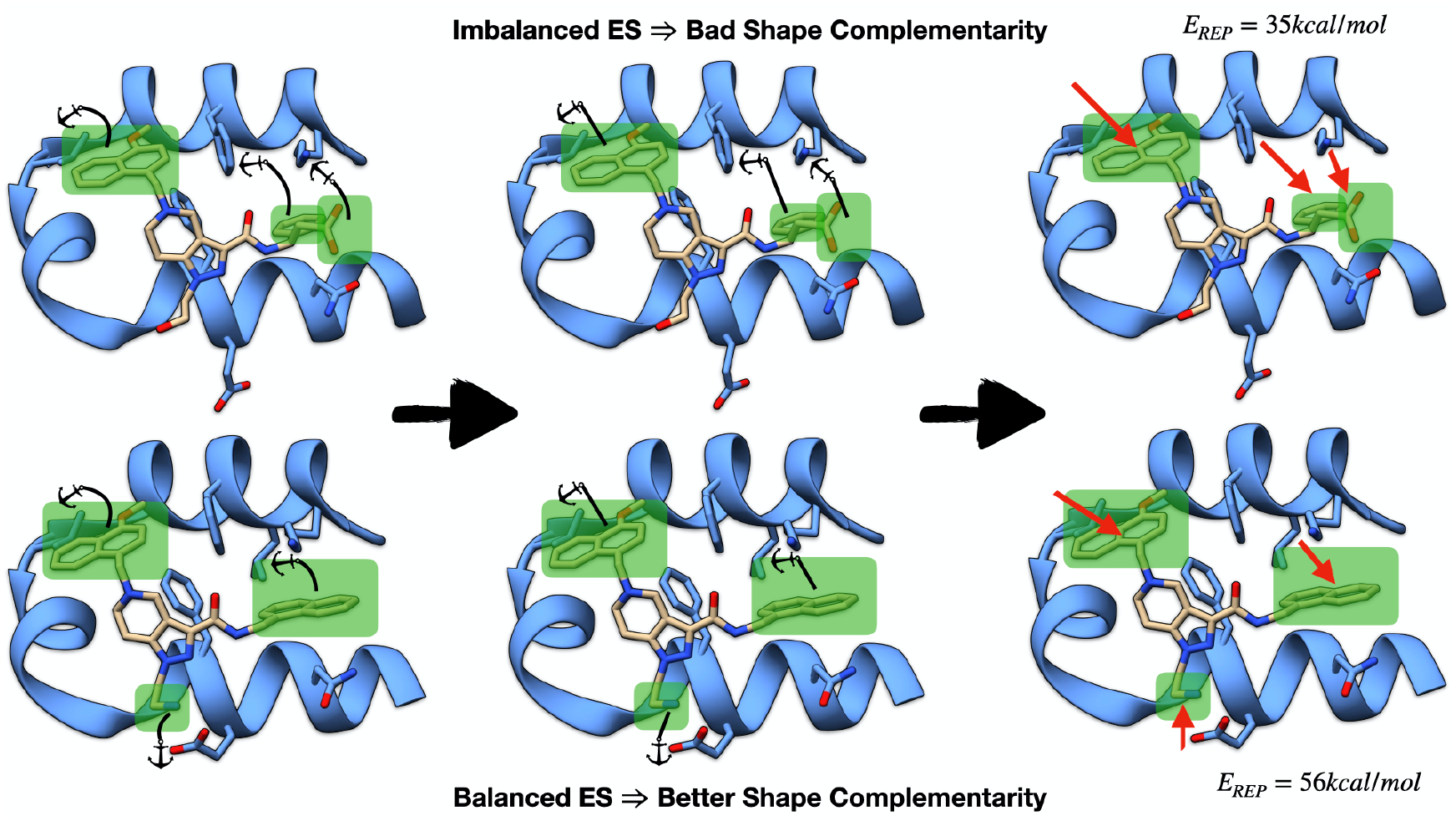
Anchoring points set by 5OML and 5N8V complexes. When anchors are set, an attractive force is created (red arrow on the last panel), which brings the respective groups together.

Two more comments are due on the EDA method herein presented. First, the fact that our algorithm is based on semi-empirical quantum chemistry allows these calculations to be performed on normal workstations affordable by any researcher. Future work will linearize the computational effort, thus extending the applicability of the program. Jorgensen and coworkers explored, in a series of manuscripts, the use of semi-empirical methods in studying enzyme reactivity.^48^ Much like Jorgensen’s approach, all our calculations were formulated in relative terms to increase accuracy. In fact, if we take the matched-molecular-pair 5OML-6RT2, for which entropies of binding are expected to be identical,^24^ then relative affinities might be predicted from binding energies only. Using Boltzmann averaged binding energies, a relative K_d_ of 38.4 is obtained 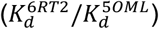. From the experimental K_d_ for 6RT2 we back-calculate a K_d_ of 16.8 μM for 5OML. Despite error cancellation benefitting all computational methods, this agreement with experimental data is an outstanding result. The question lies only on building representative enough models that can efficiently capture the fine details of binding. Additional calculations on SAR data we are currently performing further reinforce these observations, just like other studies of ours.^49^

Throughout this manuscript, several concepts in medicinal chemistry were revisited, like electrostatic-lipophilicity balance and shape complementarity. Our EDA method provided a theoretical basis bridging classical concepts with the fine determinants of inhibitory power. Most information was disclosed from analysis of experimental crystal structures. But complementary data may be retrieved from MD simulations. It is our intention to use EDA in a transformative manner in drug-lead optimization and to shed lights on biologically relevant processes.

## CONCLUSION

An Energy Decomposition Analysis method is proposed and used for calculating the behavior of the PEX14-PEX5 interface as an example of a challenging protein-protein interaction target in medicinal chemistry. EDA allows the derivation of positive guidelines for future drug-lead optimization campaigns, which are far more quantitative and accurate than classical approaches (hydrogen-bond, π-stacking, lipophilicity). Our method provides a physical basis to experimental SAR data. This is accomplished both quantitatively (numerically) and visually, in the form of maps that are intuitively interpreted by non-experts. In addition to showing what drives binding directly from experimental data, EDA eliminates crystal artefacts, simulates in-cell-like environment, and increases the value of MD simulations by revealing underlying electronic effects. Considering also the large molecular sizes our algorithm tackles, the atomistic resolution, and the simple parallelism with terminology used in classical drug discovery renders our EDA is a major advance that overcomes the low predictivity of structure-based drug discovery. The quantum chemical nature of our approach ensures that the value of the tool goes beyond biological systems, like materials science and supramolecular chemistry. We are convinced our method is a new milestone bringing full quantal analysis to biochemical simulation and finally enabling accurate rational, structure-based drug discovery.

## MATERIALS AND METHODS

The semi-empirical EDA was implemented in the ULYSSES package.^50^ Calculations reported in the main text were performed using the tight-binding GFN2-xTB^51^ with ALPB solvation.^52^ A detailed description of the method, implementation details, and in-depth benchmarks are provided in the supplementary material of the present manuscript. Dynamical simulations ran with GROMACS^53^ with the OPLS force field.^54-56^

## ACKNOWLEDGMENT

The authors gratefully acknowledge the help and suggestions of Prof. Dr. Michelle Pavanello in the constrained SCF algorithm.

