## Supplementary Information for "Can Quantum Chemistry Improve the understanding of Protein-Ligand Interactions? Implications for Structure Based Drug Discovery"

### S1 - Theory

The basis of our EDA is the method of van der Vaart and Merz,[1] which is here extended to compute further energy contributions and to include an atom-wise partitioning. The van der Vaart and Merz EDA may be classified as a variational approach because it is based on the construction of several intermediate wavefunctions.[2,3] This contrasts to approaches like SAPT, which are extensions of London's perturbational treatment.[4-6] Besides its conceptual simplicity and elegance, we chose this method because it contains no unaccountable contributions to the interaction energies.[2] Though different schemes partition energies differently, we follow the nomenclature of van der Vaart and Merz, which we complement based on the work of Su and Li.[7]

#### S1.1 The Principles of Energy Decomposition

In the supermolecular approach, the interaction energy between two molecules A and B to form a complex AB is calculated as the difference between the energies of the non-bonded complex and the isolated (and optimized) monomers.

$$\Delta \mathrm{E_{eq}} = \mathrm{E_{AB}} - \mathrm{E_A} - \mathrm{E_B} \tag{1}$$

We use here the extended meaning of the term monomer to specify each partner in the supermolecular complex, which we alternatively call dimer or complex.

EDAs aim at decomposing the above interaction energy in a manner that exposes the different types of interactions between the molecules, namely the electrostatics ($\Delta E_{es}$), polarization ($\Delta E_{pol}$), charge transfer ($\Delta E_{ct}$), exchange ($\Delta E_{exc}$), overlap-repulsion ($\Delta E_{ov}$) and the strain or deformation energy ($\Delta E_{strain}$).

$$\Delta \mathrm{E_{eq}} = \Delta \mathrm{E_{es}} + \Delta \mathrm{E_{pol}} + \Delta \mathrm{E_{ct}} + \Delta \mathrm{E_{exc}} + \Delta \mathrm{E_{ov}} + \Delta \mathrm{E_{strain}} \tag{2}$$

#### S1.2 The Strain Energy

The strain energy is obtained as the difference in energy between the bound and the free monomers, which might be in vacuum or in a solvent. For small molecule complexes, strain energies are small and typically even negligible. For large systems, the inclusion of strain energies may be essential. This is the case of most protein-ligand complexes, but we made similar observations in systems with less than 150 atoms.[8] Typically, strain energy calculations compare the geometry of the monomer in the complex against the closest equilibrium structure of the free species. This is in most cases sufficient unless one wishes to obtain entropies or Gibbs free energies of more flexible structures. In the following, we exclude deformation energies from the treatment and focus instead on the decomposition

$$\Delta \bar{E}_{eq} = \Delta E_{eq} - \Delta E_{strain} \tag{3}$$

$\Delta \bar{E}_{eq}$ contains thus all but the contribution from the deformation energies. The determination of strain energies is tackled in another publication.[9]

#### S1.3 The van der Vaart-Merz Partitioning

The EDA proposed by van der Vaart and Merz[1] partitions the interaction energy $\Delta \bar{E}_{eq}$ in 3 components: electrostatic, polarization and charge transfer. The first component is obtained by bringing the monomers from infinite separation (reference system) to their equilibrium distance without modifying their electronic densities - the electrostatic state or density. Using the definition for the energy at the HF level, the comparison between these two states yields

$$\Delta E_{es} = \frac{1}{2} tr([\boldsymbol{H}^{eq} + \boldsymbol{F}^{es}]\boldsymbol{P}^0) - \frac{1}{2} tr([\boldsymbol{H}^0 + \boldsymbol{F}^0]\boldsymbol{P}^0) + \sum_{A<B}\left(E_{AB}^{nuc,eq} - E_{AB}^{nuc,0}\right) \tag{4}$$

Here, $\boldsymbol{H}$ is the core Hamiltonian (kinetic energy and nuclear attraction), $\boldsymbol{P}$ is the (electronic) density matrix and $\boldsymbol{F}$ is the Fock matrix, which, also contains electron-electron terms. The superscript 0 refers to the infinitely separated monomers. Thus, *e.g.*, $\boldsymbol{P}^0$ is the density matrix of the isolated monomers. The superscript $eq$ refers to the complex at its equilibrium geometry. $\boldsymbol{F}^{es}$, the electrostatic Fock matrix, is calculated when the monomers are at their equilibrium distance, using however the density of the isolated species,

$\boldsymbol{P}^0$. Finally, $tr$ refers to the trace of a matrix and $E_{AB}^{nuc}$ is the nuclear repulsion between atoms A and B. Sections S8 and S9 of this document provide more information on how to construct some these quantities.

The next contribution, polarization, is obtained by optimizing the monomer densities in the presence of each other, while keeping the block-diagonal structure of the density matrix. This permits the relaxation of each molecule's charge densities due to the presence of the other species. The resulting polarization state or density, $\boldsymbol{P}^{pol}$, must be compared against the electrostatic state to yield the polarization energy.

$$\Delta E_{pol} = \frac{1}{2} tr(\boldsymbol{H}^{eq}[\boldsymbol{P}^{pol} - \boldsymbol{P}^0]) + \frac{1}{2} tr(\boldsymbol{F}^{pol}\boldsymbol{P}^{pol}) - \frac{1}{2} tr(\boldsymbol{F}^{es}\boldsymbol{P}^0) \tag{5}$$

Like the electrostatic Fock matrix, $\boldsymbol{F}^{pol}$ is the Fock matrix for the monomers at the equilibrium distance, using however $\boldsymbol{P}^{pol}$ for the calculation. Keeping the block diagonal structure of the density matrix results in a spatial restriction of the electronic densities. This means that the electrons are forced to remain in the Atomic Orbital (AO) space of each molecule in the system. We made an alternative calculation path available, in which the polarization state is obtained by means of constrained Self-Consistent-Field (SCF) and ensuring that the net electronic populations of each species in the complex are that of the isolated molecules.[10-12] Though the Mulliken population analysis might be too permissive and overestimate the polarization terms, this is consistent with the semi-empirical methods we employ. In some situations, these different calculation paths yield significantly different results.[13,14] Typically, one observes weaker charge transfer when constrained-SCF is employed.

Lastly, the charge transfer contribution is obtained by comparing the fully optimized supermolecular density against the polarization one.

$$\Delta E_{ct} = \frac{1}{2} tr(\boldsymbol{H}^{eq}[\boldsymbol{P}^{eq} - \boldsymbol{P}^{pol}]) + \frac{1}{2} tr(\boldsymbol{F}^{eq}\boldsymbol{P}^{eq}) - \frac{1}{2} tr(\boldsymbol{F}^{pol}\boldsymbol{P}^{pol}) \tag{6}$$

Contrary to other EDA-schemes,[7] the charge transfer term in the van der Vaart-Merz method is not directly obtained from the sharing of electrons between occupied and virtual Molecular Orbitals (MOs) of the monomers, but rather as relaxation of the constraints on the electronic populations on the species.

### S1.4 Overlap and Repulsion

A multitude of methods have been developed to evaluate overlap-repulsion.[15-19] Some of the simplest models assume that the overlap-repulsion contribution is proportional to some function of the overlap of electronic clouds.[16] The orbital-overlap is one of those models, and it defines,

$$\Delta E_{ov} = KS^2 = K \sum_{i \in A}^{occ} \sum_{j \in B}^{occ} \left|\langle \psi_i | \psi_j \rangle\right|^2 = K \sum_{i \in A}^{occ} \sum_{j \in B}^{occ} \left(\boldsymbol{C}^0 \boldsymbol{S}_{AO}^{eq} \boldsymbol{C}^{0\dagger}\right)_{ij} \tag{8}$$

$\boldsymbol{C}^0$ is the MO coefficient matrix obtained when the two monomers are isolated and $\boldsymbol{S}_{AO}^{eq}$ is the overlap matrix for the pair at the equilibrium distance. Though easy to compute, this approach is inappropriate for minimal basis methods since it severely underestimates overlap. Using GFN2-xTB, we calculated total repulsion energies for a few complexes in the S22[24] test-set: 2.815 kcal/mol for the water dimer; 0.476 kcal/mol for the parallel displaced dimer of benzene; 1.370 kcal/mol for the T-shaped dimer; 0.139 kcal/mol for the adenine-thymine dimer. Gresh *et al.*[19] suggested using a scaling factor K that depends on the number of occupied orbitals of both monomers, but this may lead to significant overestimation of repulsion in many situations. If we take for instance the water dimer reported above, this means a repulsion of over 45 kcal/mol. The calculations of Su and Li yield a repulsion of 16 kcal/mol.[7] To avoid these problems and empirical expressions, we resorted to the Roothaan-Hall or Pople-Nesbet equations.

### S1.5 Dispersion Contributions

Other EDA schemes resort to higher-level methods like Second Order Møller-Plesset Perturbation Theory (MP2) or Coupled Cluster (CC) to include dispersion.[7] To our knowledge, only the BLW method of Steinmann *et al.*[21] and the GKS-EDA of Su and coworkers[3,22] estimate London forces directly from a dispersion expansion that uses model parameters.

Currently, some of the most used *ad hoc* dispersion corrections for HF and DFT are Grimme's D3[23] and D4[24] methods. These are furthermore also available for most semi-empirical methods. For all NDDO methods available in our program, the dispersion corrections are coupled to hydrogen-bond corrections as well. These have the additional difficulty that they cannot be assigned to any energy term in specific. Furthermore, the dispersion contributions have not been developed in full, since neither D3H4[23,25] nor D3H+[23,26,27] contain contributions from the three-body Axilrod-Teller-Muto terms. Consequently, we currently do not calculate dispersion when NDDO methods are used. In the case of GFN2-xTB,[28] we isolate the contributions of dispersion from the total energy, and we then compare the final adduct against the reference state.

### S1.6 Partition into Atom-wise Contributions

Much like the EDA algorithms themselves, the partitioning of interaction energies into atomic contributions is not unique, as this is also not an observable.[29] For NDDO methods we split the products of $\boldsymbol{H}$ and $\boldsymbol{F}$ with $\boldsymbol{P}$ into pairwise contributions. For GFN2-xTB, though the system's energy is obtained differently, a similar partitioning is possible. We wish to stress that our atomic partitioning differs in nature from the approach followed by Merz and coworkers,[30] which was inspired by the work of Fischer and Kollmar.[31] In our case, atom diagonal terms - specific to one atom - are more influenced by their chemical environment. The consequence thereof is that the one-electron contributions to atom diagonal elements are partially cancelled out by electron-repulsion contributions, leading to more amenable values. The exception is in the electrostatic terms associated when NDDO methods are used. This is caused by the core repulsion contributions.

#### S1.7 Solvation Effects

If the interacting pair of molecules is in solution, the interaction energy $\Delta E_{eq}$ must be additionally corrected for solvation effects

$$\Delta E_{eq}^{sol} = \Delta E_{eq} + \Delta E_{solv} \tag{12}$$

There is an abundance of solvation models, each with its own intricacies. Here, we focus on ALPB,[32] for which the atom-wise partitioning is straightforward. ALPB contains additional terms from the Solvent Accessible Surface Area (SASA) and a free energy shift, a fixed quantity that is only solvent dependent. Both these terms are considered atom-specific and are not shared by pairs of atoms.

Much like the dispersion terms, solvation is estimated using the fully optimized wavefunction for the van der Waals complex and the reference state. Nevertheless, though it is not explicitly accounted for, solvation is present during the calculation of the intermediate states to avoid inconsistencies. This goes hand in hand with what is done in GKS-EDA,[3,22] and it brings several advantages. For instance, the SCF of large molecules with multiple polar groups (like proteins) is usually hard to solve, and the introduction of a dielectric medium has usually a positive impact in the convergence of the wavefunction or of the electronic density. The consequence is however that the polarization and/or charge transfer energies in solution might come out repulsive. This is because the polarization and charge transfer terms are evaluated without the solvation contributions, though the respective states are optimized with solvation included.

Though clear and consistent from the methodology point of view, this partitioning has the disadvantage that charge transfer might become repulsive in certain conditions. This is because the polarization state is determined with solvation included. Since the polarization state is a minimum in energy under a certain population constraint and in the presence of a solvent, this means that by removing the contribution of the solvent from the total energy, it is no longer ensured that the resulting energy is a minimum. Consequently, charge transfer might become repulsive. This is actually noticeable in some of the calculations reported in the main document. Nonetheless, in absence of solvation, charge transfer is expected to be attractive. This may be alternatively seen as the ability of the environment in stabilizing charged species.

### S2 – Implementation and Computational Details

The expressions presented in the previous section are valid primarily for NDDO methods, due to their resemblance to HF theory. An extension to GFN2-xTB is immediate, with the addendum that the atom-wise energies are no longer evaluated using

$$\mathrm{E} = \frac{1}{2} tr([\boldsymbol{H} + \boldsymbol{F}]\boldsymbol{P}) \tag{13}$$

In this case we proceed similarly using however the expressions given by Grimme and coworkers.[28]

The EDA algorithm was implemented in our semi-empirical package ULYSSES.[33] The current version of the program is compatible with all methods available in the package. It is important to stress that dispersion and solvation are not calculated in NDDO methods. Though no particular mechanism was introduced to keep users from running EDA with, *e.g.*, PM6-D3H+, we strongly discourage such calculations.

In the current implementation, the algorithm begins with calculations on the monomers of the van der Waals complex. These calculations take place either by considering each species in isolation (option splitmonomers = true) or by placing both molecules at exaggeratedly large distances (splitmonomers = false). Afterwards, a series of operations determine the necessary quantities for the van der Waals complex: 1) the total interaction energy, 2) electrostatics, 3) polarization, 4) repulsion. Then the actual decomposition analysis is performed.

The minimum input for the program is the van der Waals complex and its total charge. In this case, the program scouts the chemical system defined by the user and attempts to split it in two entities, which are used to start the calculation. This modality is however inconvenient for charged systems since the distributions of total charges between fragments is left unspecified. Note that the same applies to systems with spin multiplicity other than singlet. To cover such situations, it is also possible to run the EDA calculation from 3 starting structures. Here, one defines the van der Waals complex, the fragments, and the charges of each system.

Test calculations were performed using GFN2-xTB[28] and PM6.[34] Structures from the dimers were collected from the S22 and S66 databases.[35,36] The protein-ligand complex used for the calculations in the benchmark was the 5L87 complex as available from the PDB database.[37,38] For other protein-ligand systems, please refer to the main manuscript and references therein. The protein structures were protonated using H++ 4.0[39–41] considering a pH of 7, salinity of 0.15, internal dielectric constant of 10 and external of 80. The protein calculations reported in the main text were run exclusively with GFN2-xTB and using implicit solvation.

Figures were generated with UCSF Chimera, developed by the Resource for Biocomputing, Visualization, and Informatics at the University of California, San Francisco, with support from NIH P41-GM103311.[42]

### S3 – EDA of Simple Molecular Pairs

Here we dissect EDA results for simple dimers from the S22 and S66 databases.[35,36] We begin our benchmark analysis with the benzene complexes (Table S3.1). In the method column we distinguish two types of calculations. On one hand we have the (S) methods, in which the polarization terms are determined with the spatial restraint of the electronic populations. This means that the density matrix is block-diagonal. Calculations with (P) use instead populational restraints, in which the polarization state is obtained by means of constrained SCF. Phipps and coworkers[2] observed that the T-shaped benzene dimer has approximately half the dispersion stabilization of the respective sandwich complex. This is reflected on the GFN2-xTB calculations with quite good accuracy - ratio of 0.56. It is also noteworthy the fact that dispersion dominates the interaction in the $\pi$-stacked/sandwich dimer of benzene. With respect to the overlap-repulsion terms, we observe that these double when going from the sandwich to the T-shaped structure. This difference is easily rationalized when looking at the respective structures (Figure 2): for the sandwich complex, the closest contacts are 3.401 Å between H and C, and 3.375 Å between two carbons of different rings; for the T-shaped system the closest contact is 2.804 Å between one proton of one benzene ring and all the carbons of the other molecule. Despite the smaller electronic cloud around hydrogen atoms, the number of interactions and the respective proximities lead to higher repulsion for the T-shaped complex.

The electrostatic contribution in both benzene complexes is expected to be identical for several EDA analyses, at values of about −2 to −3 kcal/mol, with slightly lower values for the T-shaped complex. Su and Li also predicted similar values for the electrostatics of the T-shaped benzene dimer - $\Delta E_{es} = -3.55$ kcal/mol.[7] In our case, both semi-empirical methods predict the correct ranking of electrostatic contributions. In the case of GFN2-xTB, the relative values for the electrostatic stabilization of these complexes agree well with the literature, though the absolute values are slightly higher. In the sandwich complex, GFN2-xTB even predicts repulsive electrostatics. This might be caused by an imbalance in charge penetration contributions.[43] However, in all due fairness, the electrostatics according to DFT methods depend strongly on the method and basis set employed and may deviate by more than 15 kcal/mol.[44] In the case of PM6, the interactions are all attractive, though for the T-shaped complex the electrostatic stabilization is too favorable, at least in comparison to literature values. The reason for the discrepancy between PM6 and GFN2-xTB electrostatics stems from the fact that in the former we subtract the overlap repulsion contributions, which might lead to the overestimation of electrostatics at too short range.

**Table S3.1. Energy decomposition analysis for some model complexes taken from the S22[35] - PhH:PhH, H2O:H2O and the base-pair complexes - and S66 databases.[36] PhH stands for benzene, MeOH for methanol, A for adenine and T for thymine. In the column of structures, (S) stands for sandwich structure, (T) stands for the T-shaped complex and (WC) for Watson-Crick. Associated to methods, (S) stands for the algorithm that spatially restricts electrons, whereas (P) stands for the constrained SCF calculation. All energies in units of kcal/mol.**

| Complex | Method | $\Delta E_{ES}$ | $\Delta E_{POL}$ | $\Delta E_{CT}$ | $\Delta E_{EXC}$ | $\Delta E_{OV}$ | $\Delta E_{DISP}$ |
|---|---|---|---|---|---|---|---|
| PhH:PhH (S) | PM6 (S) | -4.236 | -0.074 | -0.263 | 0.076 | 4.623 | 0.000 |
| | PM6 (P) | -4.236 | -0.262 | 0.001 | 0.001 | 4.623 | 0.000 |
| | GFN2-xTB (S) | 0.595 | -0.344 | -0.008 | 0.000 | 0.752 | -4.813 |
| | GFN2-xTB (P) | 0.595 | -0.352 | 0.000 | 0.000 | 0.752 | -4.813 |
| PhH:PhH (T) | PM6 (S) | -8.343 | -0.050 | -1.117 | 0.264 | 8.492 | 0.000 |
| | PM6 (P) | -8.343 | -0.691 | -0.241 | 0.029 | 8.492 | 0.000 |
| | GFN2-xTB (S) | -0.329 | -1.545 | -0.601 | 0.000 | 3.940 | -2.698 |
| | GFN2-xTB (P) | -0.329 | -1.780 | -0.365 | 0.000 | 3.940 | -2.698 |
| $H_2O$:$H_2O$ | PM6 (S) | -7.987 | -0.111 | -6.759 | 2.324 | 8.596 | 0.000 |
| | PM6 (P) | -7.987 | -3.588 | -1.230 | 0.271 | 8.596 | 0.000 |
| | GFN2-xTB (S) | -4.400 | -9.517 | -2.118 | 0.000 | 11.391 | -0.316 |
| | GFN2-xTB (P) | -4.400 | -10.112 | -1.523 | 0.000 | 11.391 | -0.316 |
| $H_2O$:MeOH | PM6 (S) | -9.767 | -0.200 | -7.231 | 2.283 | 10.768 | 0.000 |

| | | | | | | | |
|---|---|---|---|---|---|---|---|
| | PM6 (P) | -9.767 | -4.193 | -0.940 | -0.105 | 10.768 | 0.000 |
| | GFN2-xTB (S) | -4.291 | -10.148 | -2.610 | 0.000 | 12.819 | -0.567 |
| | GFN2-xTB (P) | -4.291 | -11.100 | -0.658 | 0.000 | 12.819 | -0.567 |
| MeOH $H_2O$ | PM6 (S) | -6.510 | -0.134 | -0.626 | 2.155 | 7.558 | 0.000 |
| | PM6 (P) | -6.510 | -3.754 | -0.599 | -0.067 | 7.558 | 0.000 |
| | GFN2-xTB (S) | -3.964 | -9.403 | -1.577 | 0.000 | 10.853 | -0.512 |
| | GFN2-xTB (P) | -3.964 | -9.737 | -1.242 | 0.000 | 10.853 | -0.512 |
| MeOH:MeOH | PM6 (S) | -8.798 | -0.192 | -7.344 | 2.260 | 10.582 | 0.000 |
| | PM6 (P) | -8.798 | -4.431 | -0.505 | -0.339 | 10.582 | 0.000 |
| | GFN2-xTB (S) | -4.023 | -10.802 | -2.283 | 0.000 | 13.214 | -0.876 |
| | GFN2-xTB (P) | -4.023 | -11.514 | -1.571 | 0.000 | 13.214 | -0.876 |
| A:T (WC) | PM6 (S) | -32.087 | -1.321 | -20.572 | 5.611 | 39.310 | 0.000 |
| | PM6 (P) | -32.087 | -15.704 | 0.535 | -1.113 | 39.310 | 0.000 |
| | GFN2-xTB (S) | -9.474 | -36.233 | -2.570 | 0.000 | 35.570 | -3.224 |
| | GFN2-xTB (P) | -9.474 | -37.942 | -0.861 | 0.000 | 35.570 | -3.224 |
| A:T (S) | PM6 (S) | -15.968 | -0.386 | -1.039 | -0.073 | 12.524 | 0.000 |
| | PM6 (P) | -15.968 | -1.463 | 0.045 | -0.080 | 12.524 | 0.000 |
| | GFN2-xTB (S) | -4.082 | -2.463 | -0.055 | 0.000 | 3.327 | -9.020 |
| | GFN2-xTB (P) | -4.082 | -2.428 | -0.089 | 0.000 | 3.327 | -9.020 |

When it comes to polarization, contributions are usually attractive, though quite small in magnitude. The exception is the polarization for the T-shaped complex obtained from GFN2-xTB, which mismatches the rest of the calculations. Nevertheless, the value is not exceedingly large. This is because for this method we are calculating polarization mixed with exchange. As discussed by Hayes and Stone,[15] exchange contributions have no explicit dependency on overlap. Thus, when we calculate the GFN2-xTB repulsion terms, we are not estimating the electronic exchange, which is then not subtracted from the GFN2-xTB polarization. Though for correctness we should name these terms exchange-polarization, we decided to keep the nomenclature consistent between all methods we employ. Except for the PM6 (S) calculations, the two semi-empirical models employed agree on the fact that polarization is stronger in the T-shaped complex. This contrasts with other calculations.[2] Though not investigated in detail, we believe this is a consequence of the minimal basis set used in the calculations. In the case of charge transfer, the semi-empirical calculations mirror the higher-level analyses:[2] low values in magnitude and larger for the T-shaped complex.

Next, we look at the complexes involving water and methanol. It has been previously observed[2] that water-water and water-methanol complexes have roughly the same electrostatic contributions, which may deviate by at most 0.5 kcal/mol. This is fully reflected in the GFN2-xTB analysis, and we furthermore extend this prediction to the methanol dimer. We stress that such a result captured by the semi-empirical EDA is not entirely intuitive: though methanol is not a fantastic nucleophile, it is expected to be superior to water due to charge donation from the methyl group to the oxygen atom. One would therefore expect that electrostatics are more favorable when one water is replaced by one molecule of methanol. In the case of the PM6 analysis there is more variation in the values, which arises from the subtraction of overlap-repulsion from the electrostatics.

Our GFN2-xTB polarization terms in the water dimer seem to be exceedingly large. As discussed above, this is caused by the contamination of these terms by exchange. This may easily be traced on the data reported by Su and Li:[7] their summed contribution of exchange-polarization (-11.23 kcal/mol) matches quite well our summed polarization and charge transfer terms (-11.64 kcal/mol). This is because Su and Li do not separate charge transfer from polarization. Though the PM6 values differ significantly according to the path used to calculate the polarization state, the relative values are in good agreement with the literature.[2] Note that in the PM6 calculation exchange and polarization are properly separated from one another. Again, charge penetration effects may lead to imbalances in some terms. Regarding charge transfer, the agreement also seems to be good with other higher-level terms, especially for GFN2-xTB. As noted by van der Vaart and Merz,[1] the PM6 (S) and GFN2-xTB (S) calculations are closer to Natural Bond Orbital EDA.[45-48]

Though their EDA scheme differs from ours in several key points, Su and Li were,[7] to the best of our knowledge, the only ones that performed calculations on the Adenine-Thymine Watson-Crick (WC) pair. In their calculations, a strong stabilization by electrostatics (−30.35 kcal/mol) and by exchange (−40.21 kcal/mol) is predicted. Polarization and dispersion are minor contributions - respectively −14.02 and −6.68 kcal/mol. These are however essential for binding, since the two most prominent attractive interactions are overcompensated by overlap-repulsion: 74.13 kcal/mol. Compared to all other case studies, we also verify a steep increase in the repulsion terms, and this is due to the 3 short contacts from hydrogen bonding. In PM6 calculations, overlap repulsion contributes with as much as 44.29 % to the interaction, whereas in GFN2-xTB it amounts to 40.81 %. This is in excellent agreement with the calculations of Su and Li, who estimated 44.82 %. The main interactions compensating the overlap-repulsion terms are however electrostatics and polarization, and this is an effect of how the EDA-methods are constructed: PM6-based calculations place more emphasis on electrostatics, whereas in GFN2-xTB there is a stronger accent on (exchange-) polarization. It is also worth noting that the dispersion contribution calculated with GFN2-xTB (3.70 %) also matches quite well the higher-level calculations of Su and Li (4.04 %). Charge transfer typically plays a minor role, except for the PM6 (S) and GFN2-xTB (S) calculations. The latter value seems however to be exaggeratedly large.

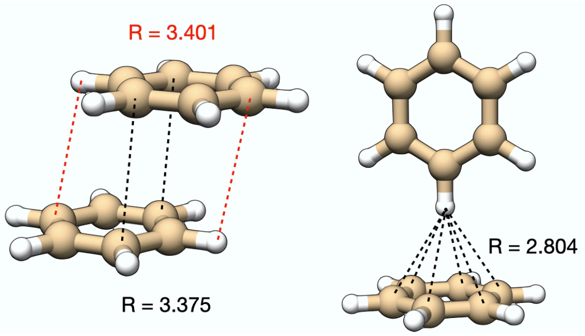


**Figure S3.1. Some interatomic distances in the benzene complexes.**

We also performed the energy decomposition for the sandwiched Adenine-Thymine pair. In this case there is a significant percentual decrease in most contributions. Repulsion and charge transfer drop by up to a factor of 10, electrostatics are halved and the drops in polarization are even more pronounced. On the other hand, dispersion becomes 3 times stronger. The interaction energy in the WC pair is nevertheless still stronger. These results are in good agreement with the analysis done above for the benzene dimers.

### S4 – EDA on Energy Surfaces: S66×10

To benchmark the semi-empirical EDA for a larger range of interaction patterns and configurations, we ran calculations on the S66×10 database.[49] The whole set of calculations is available in additional files (*c.f.* below for more details). Here, we focus on general observations and on specific complexes of biological interest, namely the complexes involving the amide (peptide) - with water, methanol, and itself - and Uracil-Uracil complexes (the respective selection of PESs are also available in S6, due to the extensiveness of the material).

In most cases, the GFN2-xTB curves seem to be simple continuations of the results obtained for the equilibrium structures. Though no complex was optimized in this work, we still try to verify how well GFN2-xTB reproduces the reference energy surfaces. In broad terms, interactions with $\pi$ bonds are the least qualitative in the systems studied. A closer look at the energy surfaces reveals that overall, the short-range domain of the Potential Energy Surfaces (PESs) either lack repulsion or are too attractive. This is particularly affecting the less polar systems, since the benzene dimer shows more deviations than the pyridine dimer, which itself shows more deviations than the Uracil dimer. Similar observations were made in a previous study of ours,[50] where equilibrium structures were too close. An analysis of the fractions for the different contributions at different intermolecular distances seems to indicate that in the short-range there is an overestimation of dispersion (*c.f.* S5), which is still steeply decreasing (with decreasing distance). This contrasts the situation observed in the systems where the agreement is good. Similar observations are made for alkanes, in particular the acyclic ones. When it comes to hydrogen-bonds, we observed a good reproduction of the reference data, though when nitrogen atoms act as acceptors there are larger deviations. More polar systems seem to be improved.

Though the overall agreement using PM6 based methods is good, there are a few cases in which PM6-D3H4 is more attractive after the equilibrium distance than PM6-D3H+ and the reference data. There are however many cases, for which PM6-D3H+ lacks repulsion in the short-range domain. The most graving case would be the dimer of cyclopentane, which also severely impacts PM6-D3H4. Here, no PM6-based method shows a proper and well-defined minimum on the energy surface. Though at first sight it might be tempting to blame dispersion, a closer analysis reveals that this is a happens already with the native method (the

PM6 interaction energy at $R_{AB}/R_{eq}$ = 0.7 is −0.226 kcal/mol). A thorough analysis is beyond the scope of this work. Nevertheless, the data we collect suggests lack of repulsion between atoms of carbon.

**Table S4.1. PES descriptors for the semi-empirical methods. The main value in each cell is the average value, in parenthesis the standard deviation.**

| Descriptor | GFN2-xTB | PM6-D3H4 | PM6-D3H+ |
|---|---|---|---|
| Matching $R_{eq}$ | 30 % | 58 % | 15 % |
| $\overline{\Delta E}_{min}$ | 0.532 (0.704) | 0.113 (0.614) | -0.046 (0.882) |
| $\overline{\|\Delta E\|}_{min}$ | 0.734 (0.487) | 0.470 (0.406) | 0.672 (0.568) |
| $\overline{\Delta E}_{Rref}$ | 0.596 (0.691) | 0.172 (0.617) | 0.125 (0.800) |
| $\overline{\|\Delta E\|}_{Rref}$ | 0.751 (0.515) | 0.469 (0.433) | 0.562 (0.579) |
| $\Delta \int PES$ | 0.128 (1.022) | 0.170 (0.689) | -0.065 (0.952) |
| $\Delta \left\|\int PES\right\|$ | 0.733 (0.718) | 0.550 (0.448) | 0.761 (0.581) |

In order to understand how well the semi-empirical methods reflect the reference PES, several descriptors were calculated. These are gathered in Table 2. The first descriptor is the ability to identify the reference equilibrium structure, *i.e.*, whether the minimum of the method matches the minimum of the reference data. PM6-D3H4 is the best performing method, where the correct minimum is captured 58% of the times. Then follows GFN2-xTB with 30%, and then PM6-D3H+, where only in 15% of the cases the minimum matches the reference data. To evaluate the quality of energy prediction, we look at the average difference between minima ($\overline{\Delta E}_{min}$ and $\overline{|\Delta E|}_{min}$) and the average energy difference at the reference equilibrium structure (energy at the minimum of reference curve, $\overline{\Delta E}_{Rref}$ and $\overline{|\Delta E|}_{Rref}$). The mean absolute deviations show, once again, that PM6-D3H4 is the best performing method. Though the mean absolute deviation for PM6-D3H+ is smaller than for GFN2-xTB, the former also shows significantly larger standard deviation. This means that the deviations in GFN2-xTB are more systematic, an observation further reinforced when comparing the mean deviation and the mean absolute deviation for PM6-D3H+. We also evaluated the root mean square deviation for PM6-D3H4, for which we obtained the value of 0.636 kcal/mol. This compares well with the value of Hostaš and coworkers (0.68).[51] Lastly, we analyze the ability of the methods to reproduce the reference energy surfaces. By means of the trapezoidal rule, the difference of the integrals of the PESs against the reference ones were calculated ($\Delta \int PES$ and $\Delta|\int PES|$). Again, PM6-D3H4 seems to be superior to the other two methods, followed by GFN2-xTB and only then PM6-D3H+. We note that this analysis is based on pre-optimized higher-level structures and does not reflect how the methods behave stand-alone. Here we refer the reader to the work of Hostaš and coworkers,[51] who verified that PM6-D3H4 may indeed be used for structure optimization of the complexes in the S22 and S66 databases.

Based on our calculations, it seems that, as is, PM6-D3H+ can only be recommended for energy evaluation but not for structural optimization. Though apparently PM6-D3H4 yields higher quality PESs than GFN2-xTB, there are some potential theoretical limitations in PM6 which suggest at least caution when using the method.[52,53]

Below we present a series of selected energy surfaces for a few van der Waals pairs that model interactions of biological relevance. More energy surfaces from the S66×10 database are found in the excel files PM6_3mol.xlsx, GFN2_3mol.xlsx and GFN2_3mol_solv.xlsx.

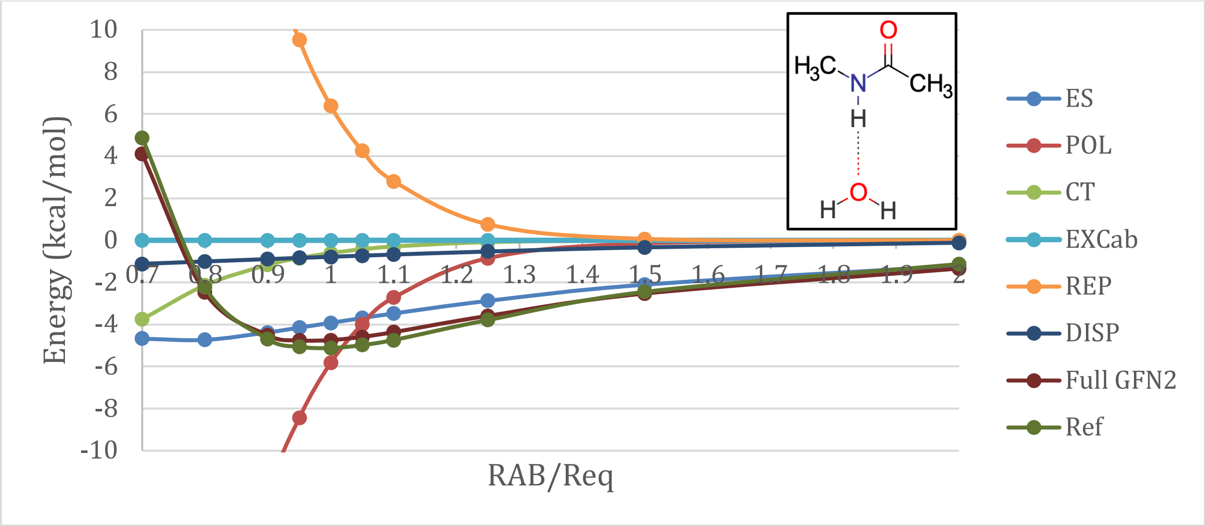
Energy (kcal/mol)
RAB/Req
10 8 6 4 2 0 -2 -4 -6 -8 -10
0.7 0.8 0.9 1 1.1 1.2 1.3 1.4 1.5 1.6 1.7 1.8 1.9 2
ES
POL
CT
EXCab
REP
DISP
Full GFN2
Ref
H3C N CH3 O H
H O H

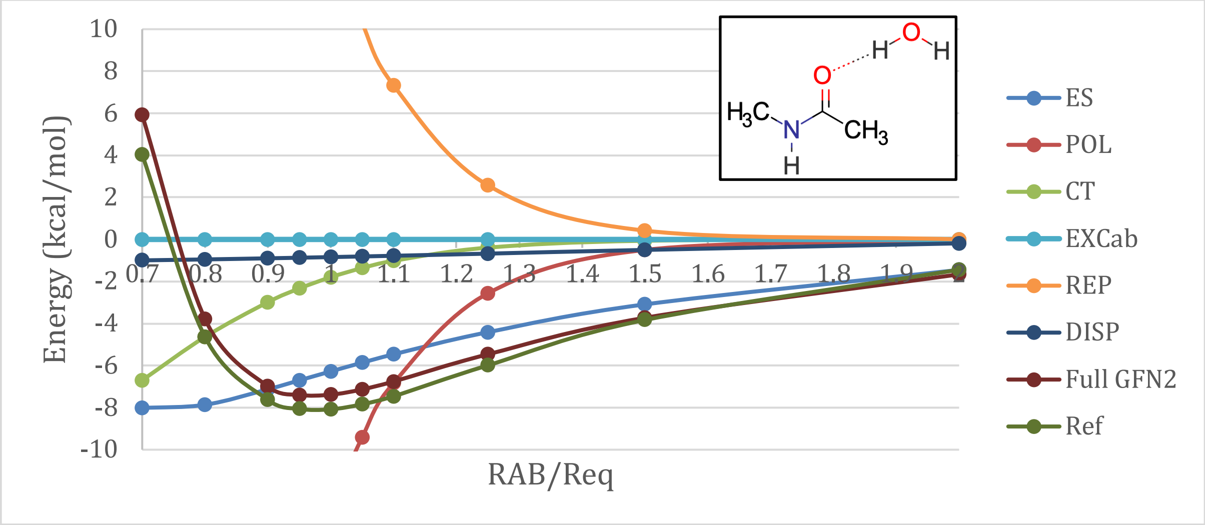
Energy (kcal/mol)
RAB/Req
10 8 6 4 2 0 -2 -4 -6 -8 -10
0.7 0.8 0.9 1 1.1 1.2 1.3 1.4 1.5 1.6 1.7 1.8 1.9 2
ES
POL
CT
EXCab
REP
DISP
Full GFN2
Ref
H O H
O
H3C N CH3
H

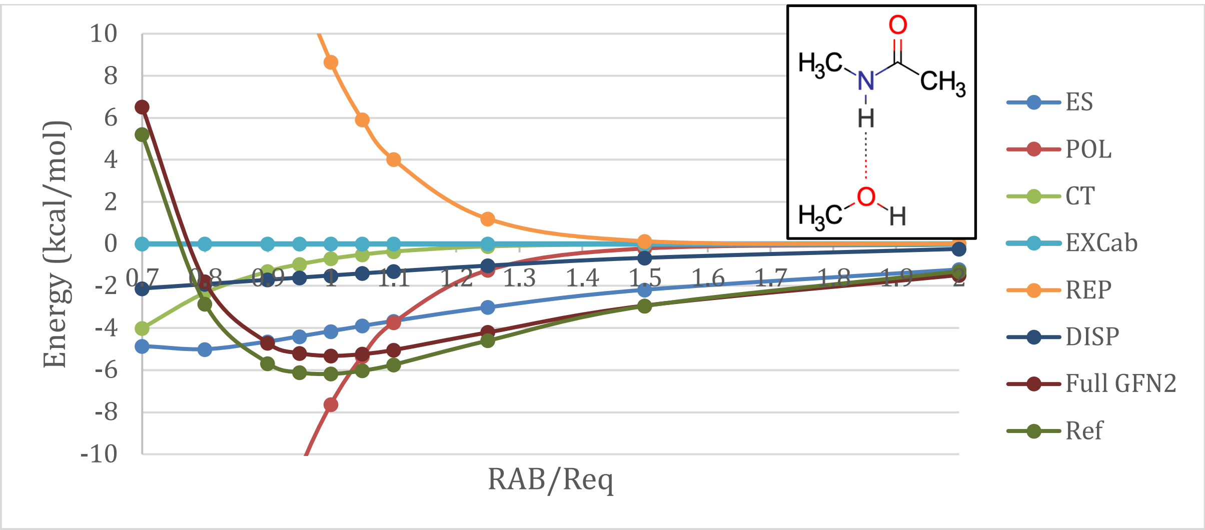
Energy (kcal/mol)
RAB/Req
10 8 6 4 2 0 -2 -4 -6 -8 -10
0.7 0.8 0.9 1 1.1 1.2 1.3 1.4 1.5 1.6 1.7 1.8 1.9 2
ES
POL
CT
EXCab
REP
DISP
Full GFN2
Ref
H3C N CH3 O H
H3C O H

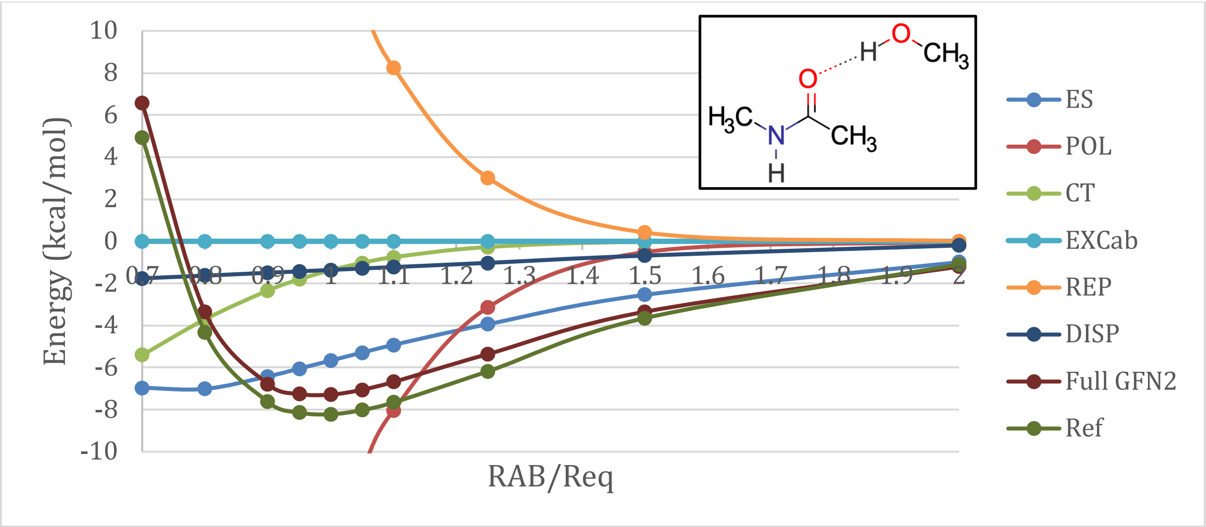
Energy (kcal/mol)
RAB/Req
ES
POL
CT
EXCab
REP
DISP
Full GFN2
Ref

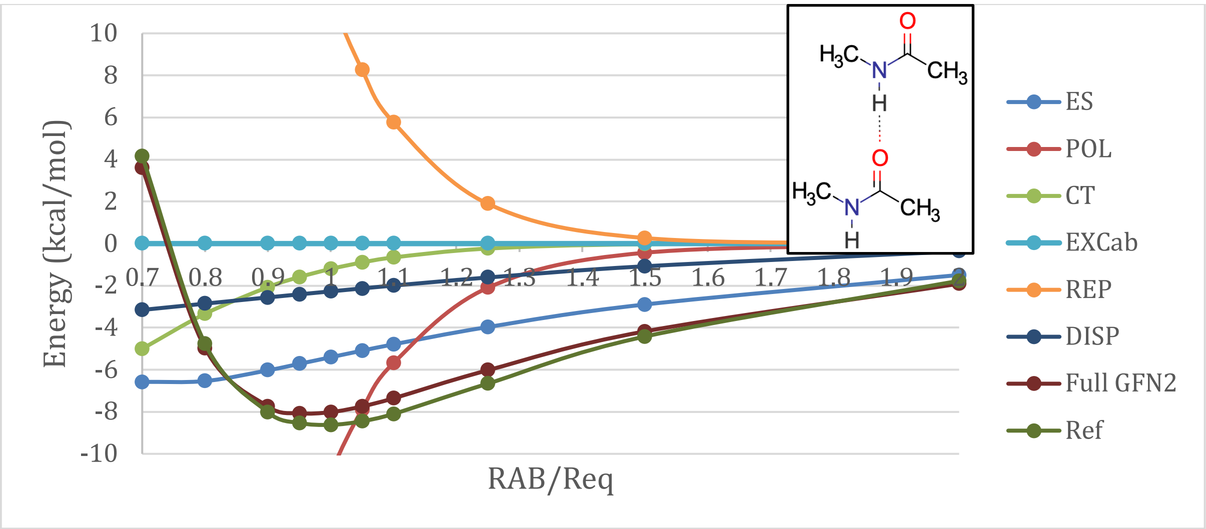
Energy (kcal/mol)
RAB/Req
ES
POL
CT
EXCab
REP
DISP
Full GFN2
Ref

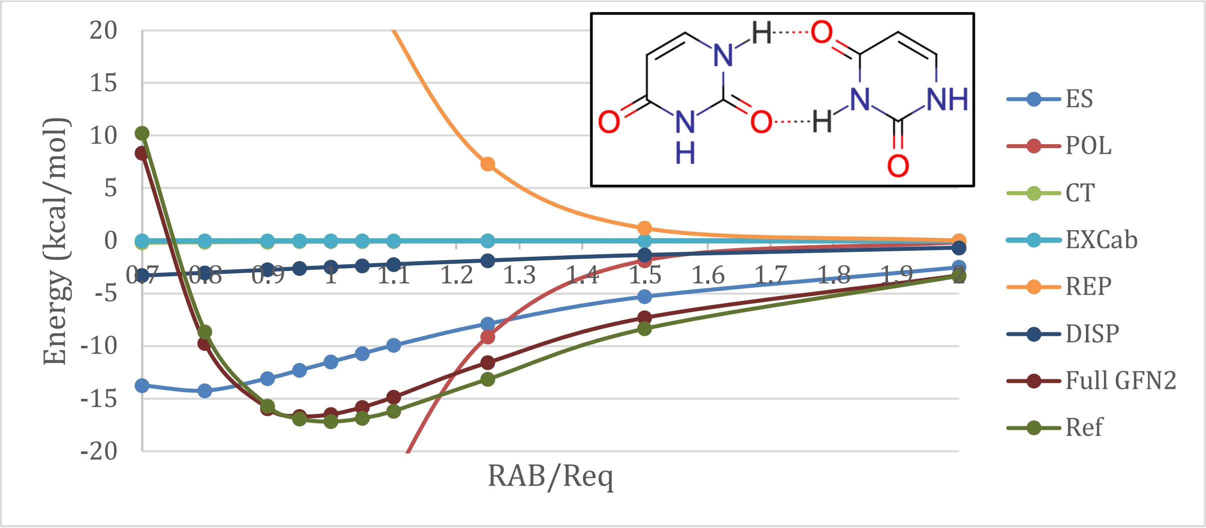
Energy (kcal/mol)
RAB/Req
ES
POL
CT
EXCab
REP
DISP
Full GFN2
Ref

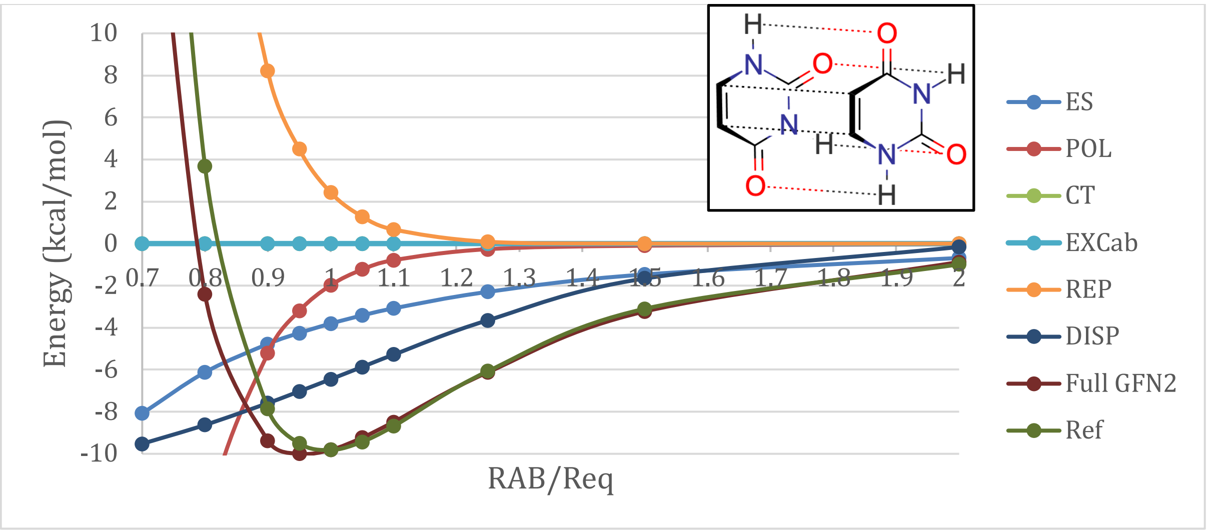


**Figure S4.1. Selected energy surfaces for GFN2-xTB in gas phase.**

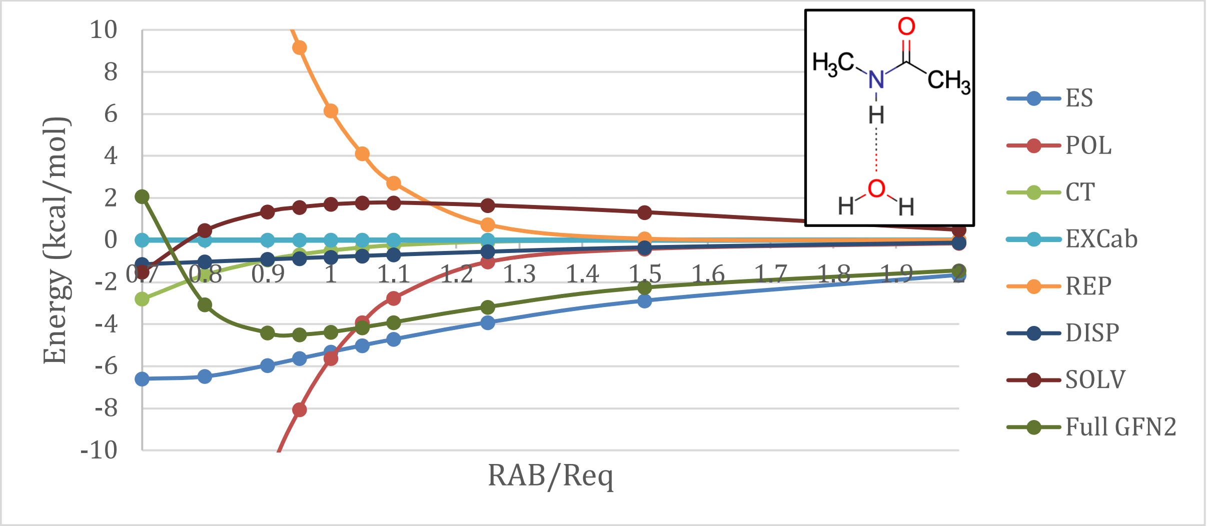


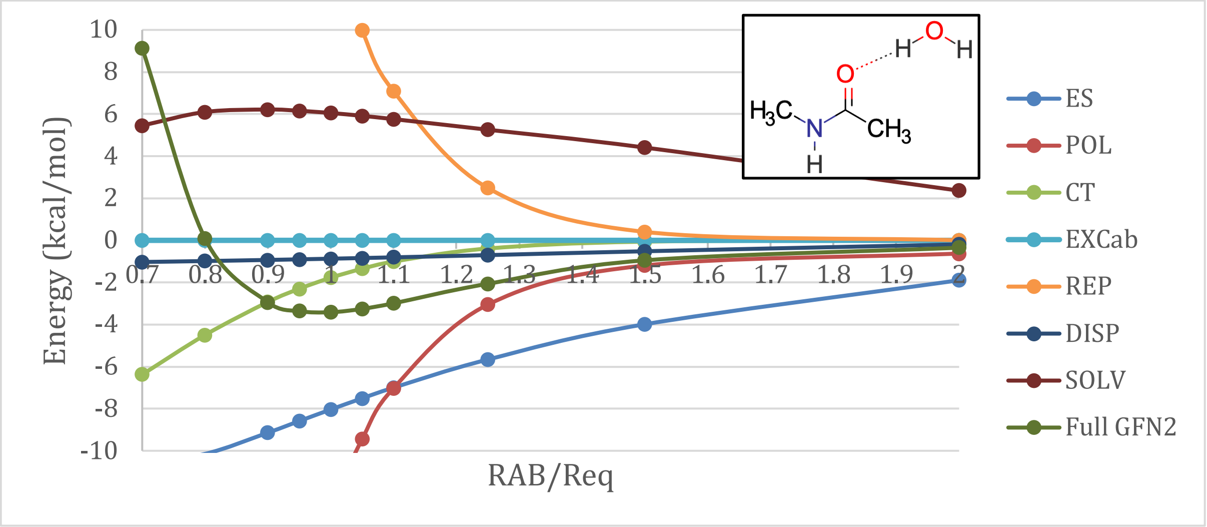

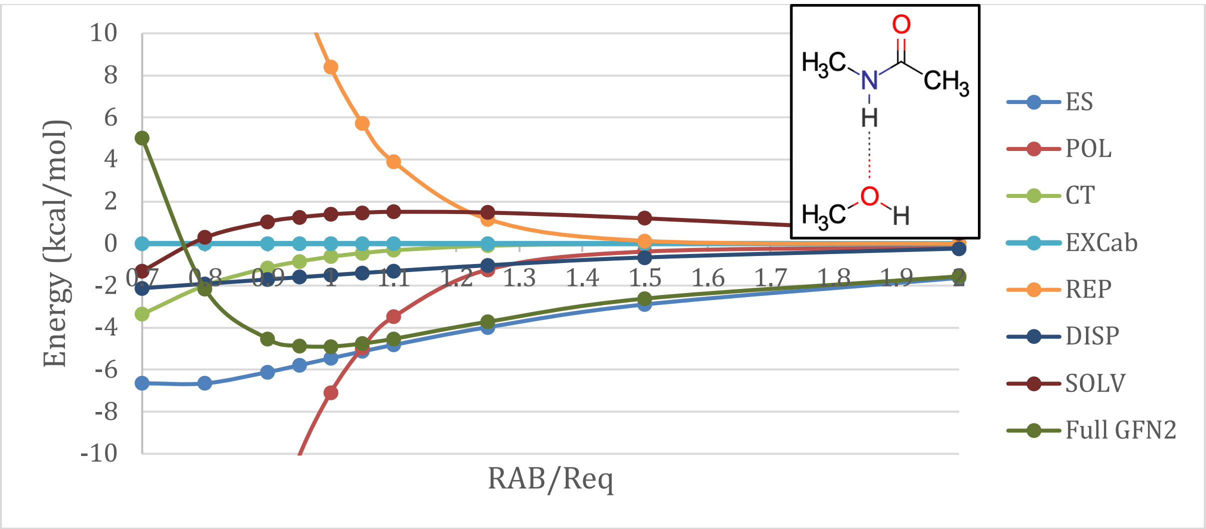
Energy (kcal/mol)
RAB/Req
10
8
6
4
2
0
-2
-4
-6
-8
-10
0.7
0.8
0.9
1
1.1
1.2
1.3
1.4
1.5
1.6
1.7
1.8
1.9
2
ES
POL
CT
EXCab
REP
DISP
SOLV
Full GFN2

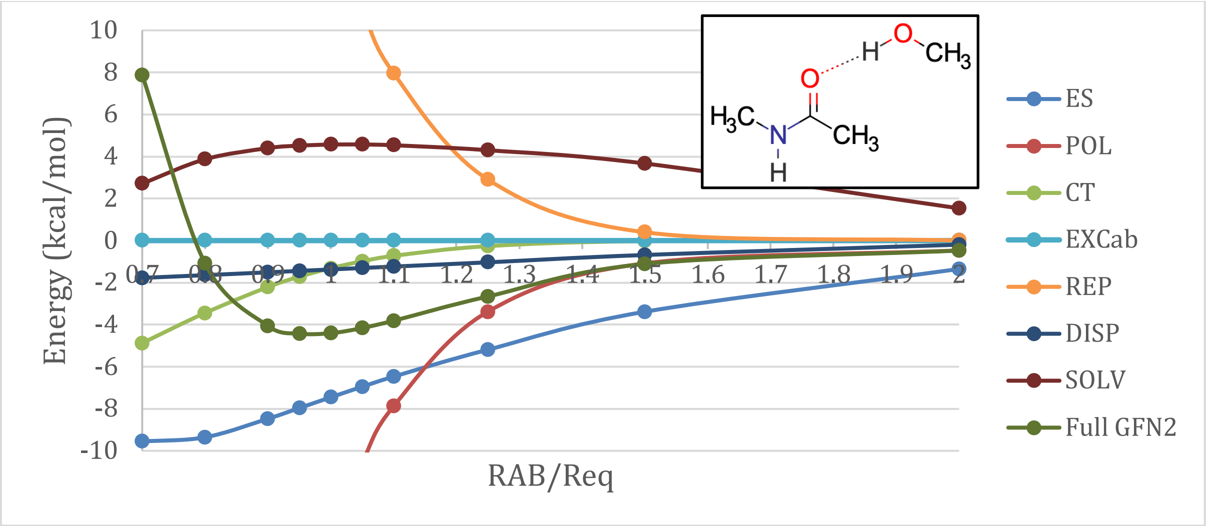
Energy (kcal/mol)
RAB/Req
10
8
6
4
2
0
-2
-4
-6
-8
-10
0.7
0.8
0.9
1
1.1
1.2
1.3
1.4
1.5
1.6
1.7
1.8
1.9
2
ES
POL
CT
EXCab
REP
DISP
SOLV
Full GFN2

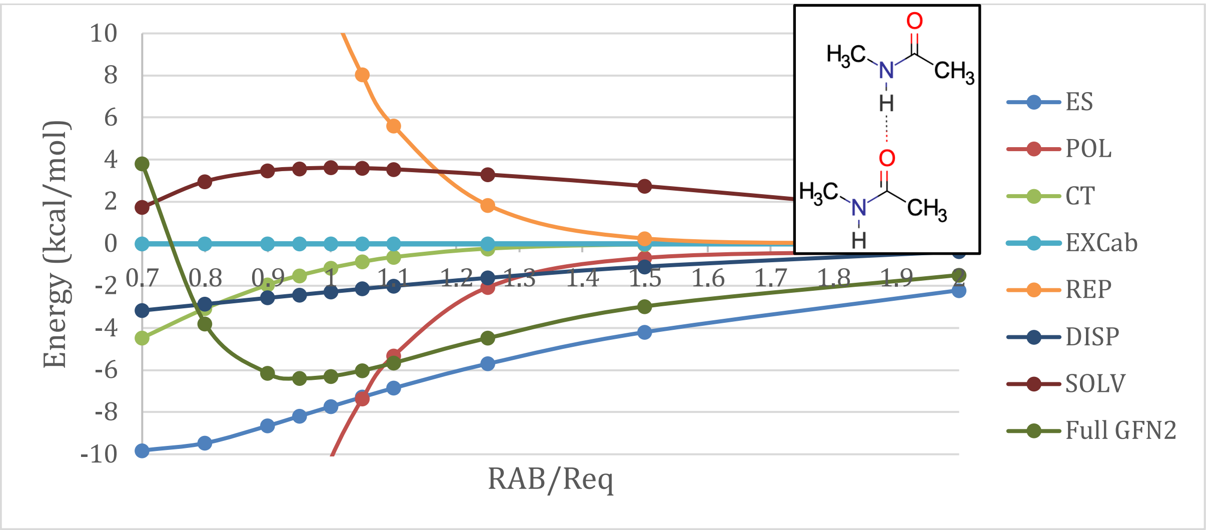
Energy (kcal/mol)
RAB/Req
10
8
6
4
2
0
-2
-4
-6
-8
-10
0.7
0.8
0.9
1
1.1
1.2
1.3
1.4
1.5
1.6
1.7
1.8
1.9
2
ES
POL
CT
EXCab
REP
DISP
SOLV
Full GFN2

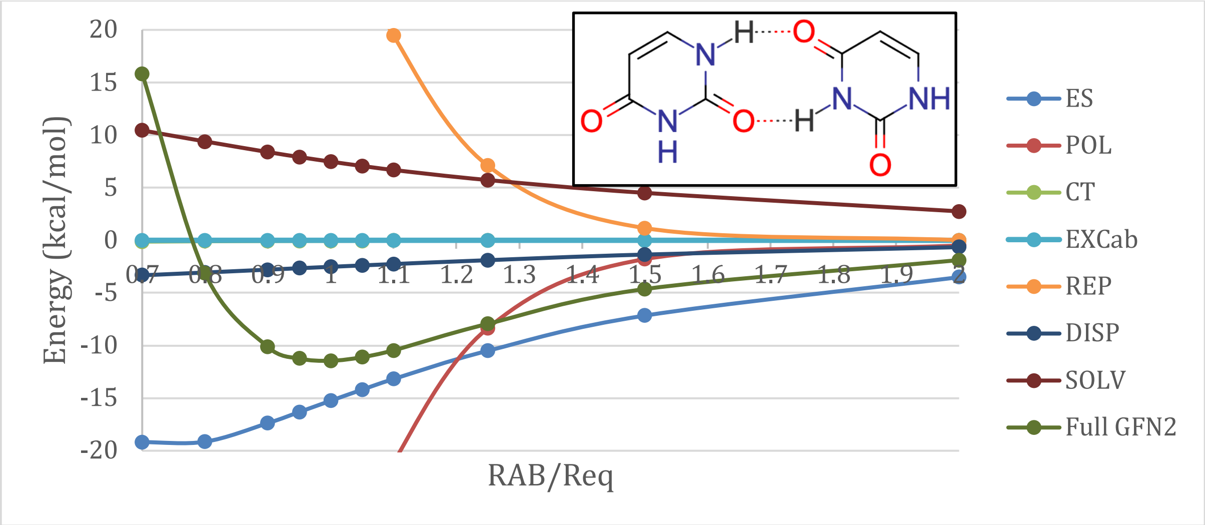


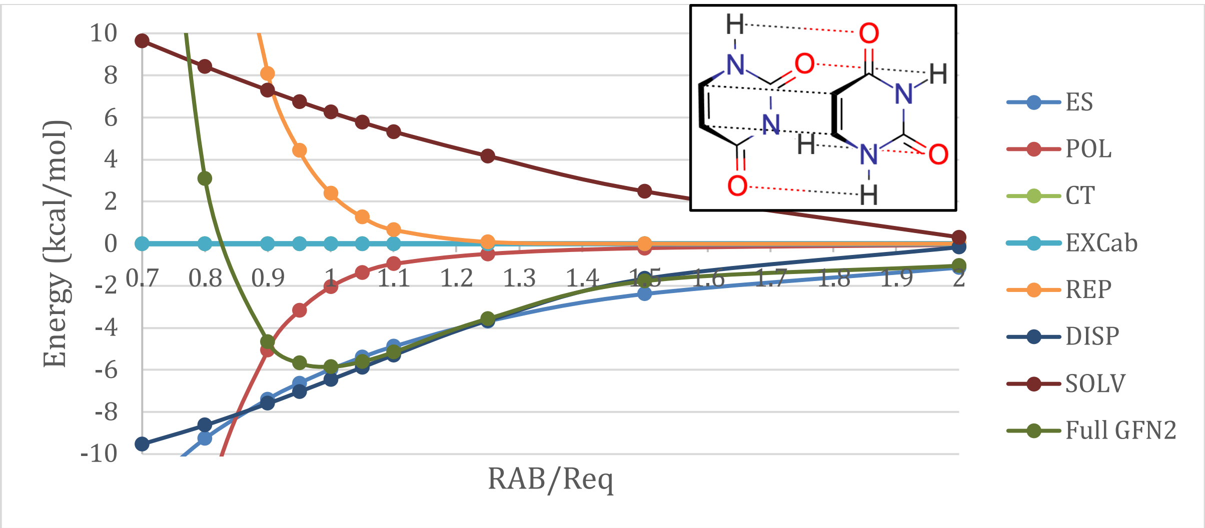


**Figure S4.2. Selected energy surfaces for GFN2-xTB in water's dielectric.**

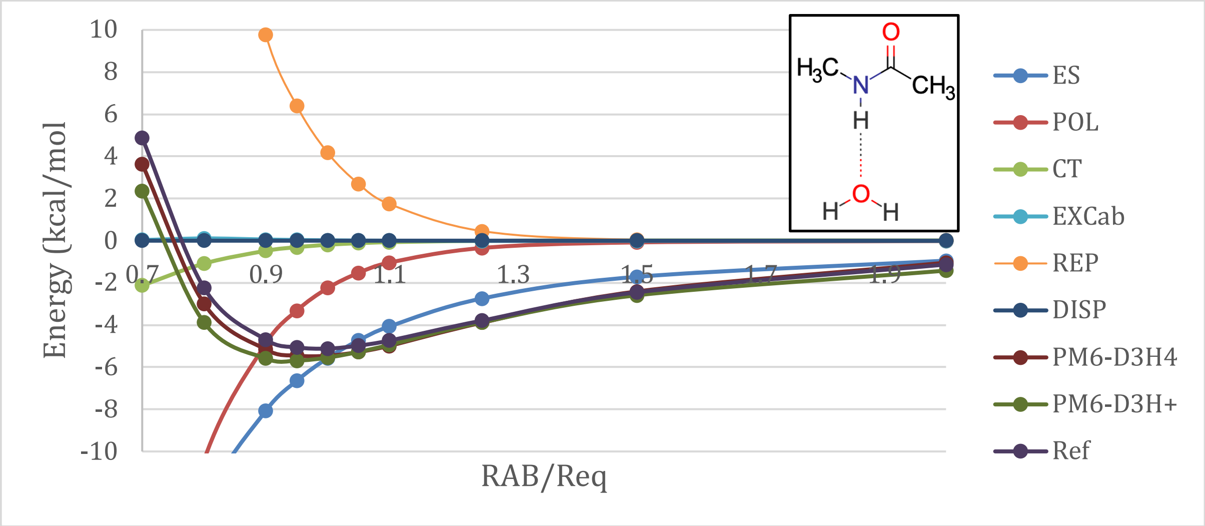

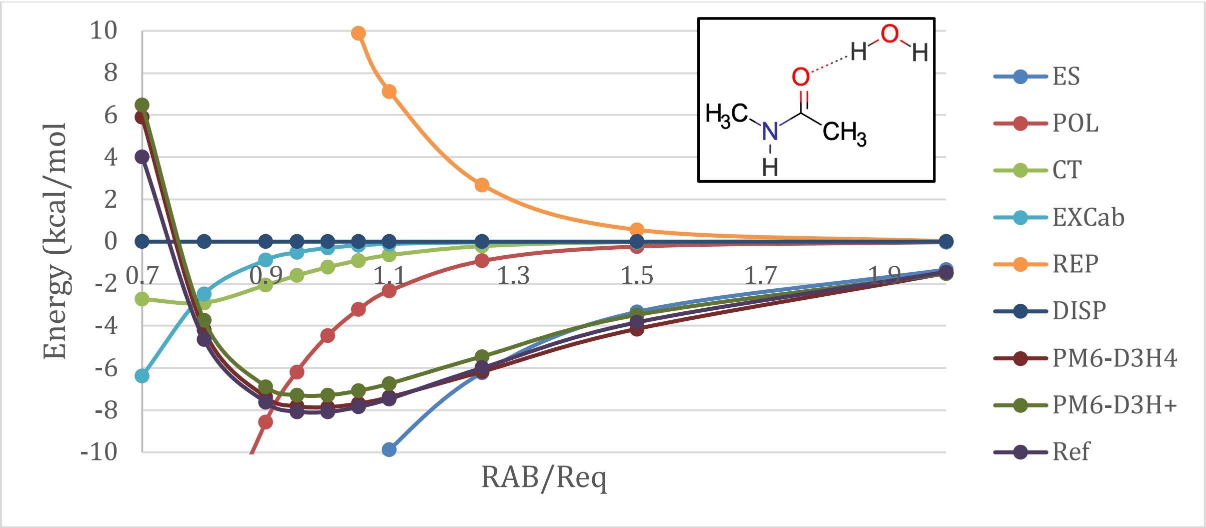
Energy (kcal/mol
RAB/Req
10
8
6
4
2
0
-2
-4
-6
-8
-10
0.7
0.9
1.1
1.3
1.5
1.7
1.9
ES
POL
CT
EXCab
REP
DISP
PM6-D3H4
PM6-D3H+
Ref
O
H
H
H3C
N
H
CH3

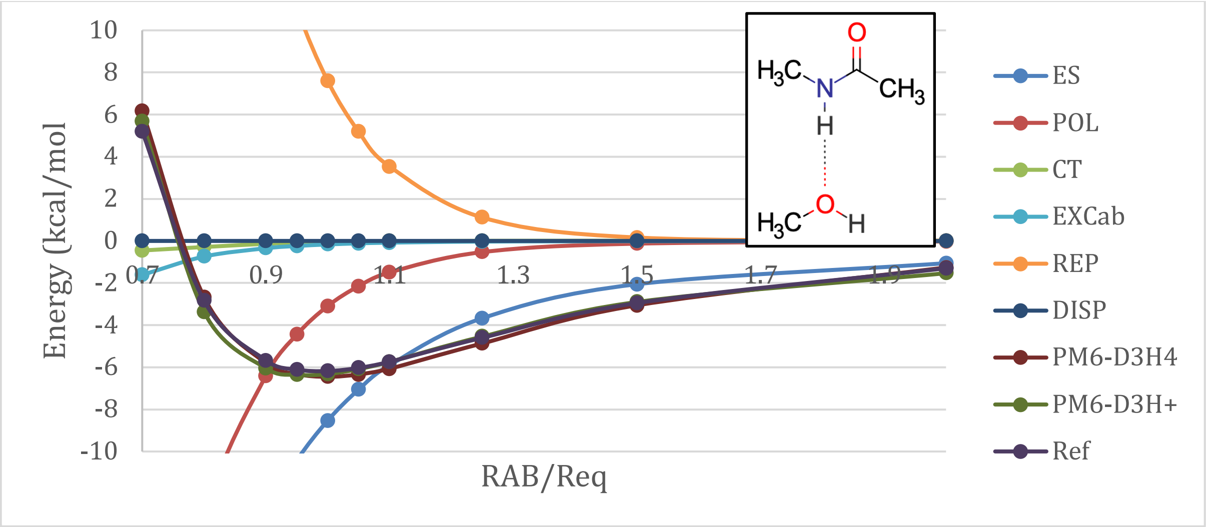
Energy (kcal/mol
RAB/Req
10
8
6
4
2
0
-2
-4
-6
-8
-10
0.7
0.9
1.1
1.3
1.5
1.7
1.9
ES
POL
CT
EXCab
REP
DISP
PM6-D3H4
PM6-D3H+
Ref
O
H3C
N
CH3
H
H3C
O
H

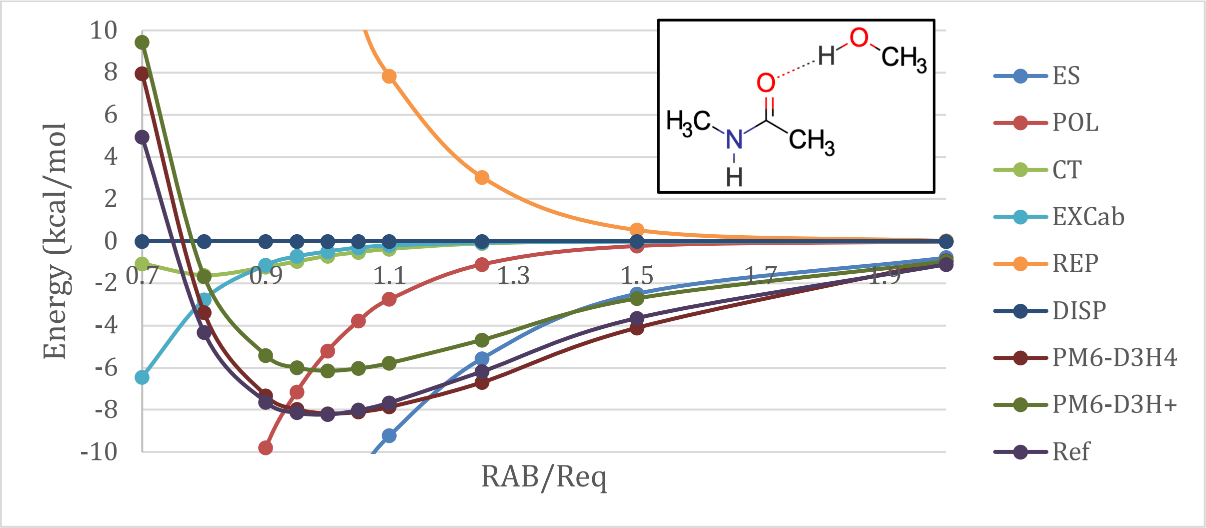
Energy (kcal/mol
RAB/Req
10
8
6
4
2
0
-2
-4
-6
-8
-10
0.7
0.9
1.1
1.3
1.5
1.7
1.9
ES
POL
CT
EXCab
REP
DISP
PM6-D3H4
PM6-D3H+
Ref
H
O
CH3
O
H3C
N
CH3
H

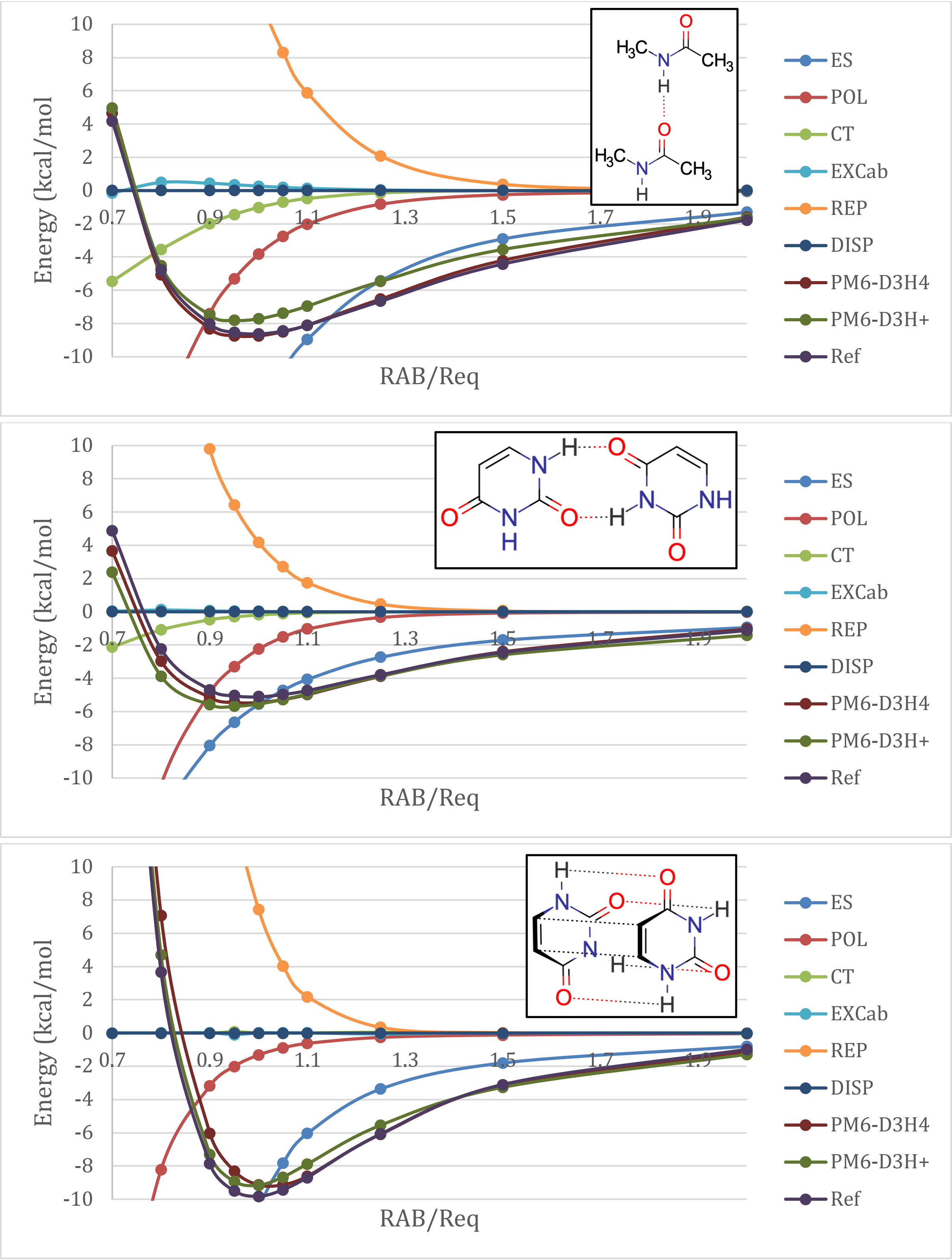


**Figure S4.3. Selected energy surfaces for PM6-based methods in gas phase.**

As for the selected PESs on model systems for biological interactions, we verify that GFN2-xTB can generally reproduce well the reference data. The largest deviations come perhaps for the already discussed π-stacked dimer of Uracil. Together with the amide dimer, the Uracil dimers show larger deviations at the equilibrium geometry. Hydrogen-bonds are typically well described, particularly close to the equilibrium complex. Except for the sandwich dimer of Uracil, intermolecular interactions are systematically shifted by up to +1 kcal/mol. PM6-D3H4 shows significantly smaller deviations than the related PM6-D3H+, and also than GFN2-xTB. The reproduction of the reference energy surfaces is also far better using PM6-D3H4. Though there are situations where the agreement with reference data is excellent, there are also some interactions which are improperly described by PM6-D3H+. This is for instance the case of the amide(A)-water(D) complex against the amide(A)-methanol(D) one. We find no obvious reason for this observation.

Irrespective of the method used for EDA, the long-range domain of the PESs is dominated by electrostatics. Dispersion is also an important contribution, particularly for the sandwich dimer of Uracil. In fact, for this π-stacked dimer, dispersion effects even dominate over electrostatics for intermolecular distances up to 1.5-2 times the equilibrium geometry. In the short-range domain, polarization dominates the attractive forces due to strong orbital overlap. It is also in the region of small intermolecular distances that charge transfer gains weight. Nevertheless, this interaction rapidly zeroes as the intermolecular distances increase. In some cases, the PM6 EDA shows a decrease in the magnitude of charge transfer at very short molecular contacts. Our tests revealed thus far that this only happens at very short distances, where there is no practical relevance (the interaction between molecules is already repulsive). Finally, we observe that the in most cases the overlap-repulsion between molecules starts increasing at $R_{AB} = 1.25R_{eq}$, though in some cases the deviation is very soft.

### S5 – Analysis of π-Stacking

Below we plot the fraction of the binding energy for each energy contribution at several intermolecular distances for the pyridine (P:P) and Uracil (U:U) dimers. Calculations are done with GFN2-xTB. To simplify the analysis, each energy contribution was assigned a color. The pyridine dimer curves have full lines, whereas the Uracil dimer is represented by dashed lines.

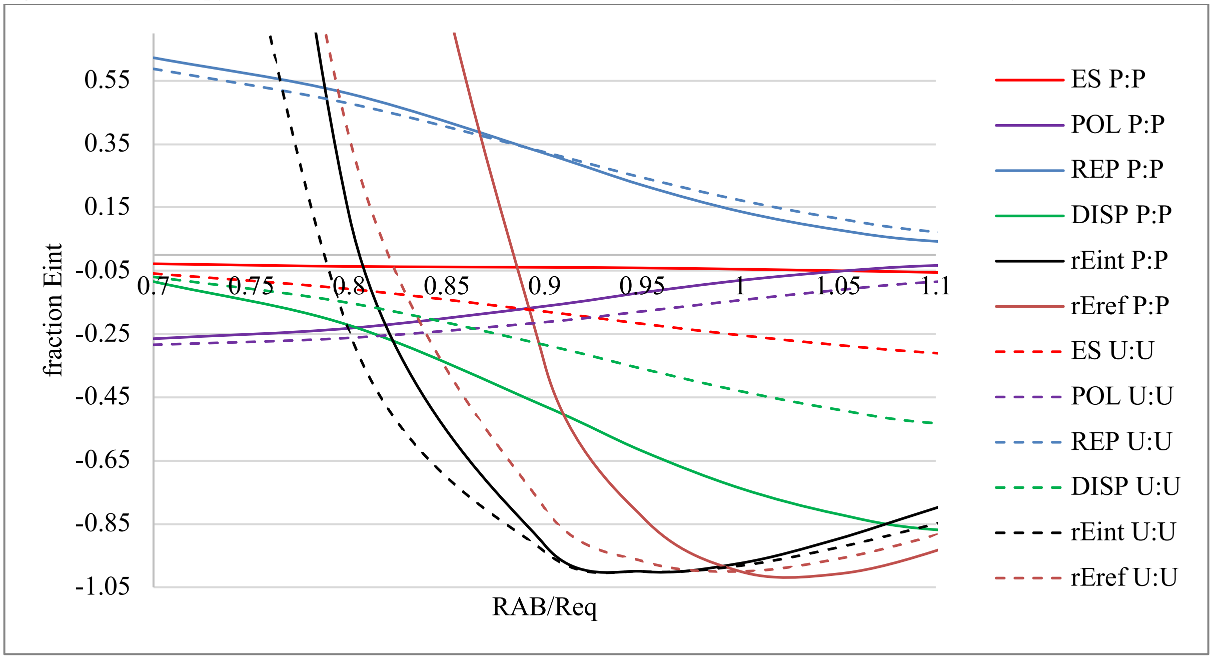


**Figure S5.1. Energy Decomposition Analysis of the uracil and pyridine dimers using GFN2-xTB.**

Acknowledging that the GFN2-xTB curves are too attractive in the short-range domain, and that this overestimation of attraction is more pronounced for the pyridine dimer than for the Uracil dimer, then we should look for the energy contribution that is stronger in the P:P pair than in the U:U one. This is only dispersion. We note that even repulsion is stronger for the pyridine dimer than it is for the Uracil one.

### S6 – Truncations of a Protein-Ligand Complex

One driving force for the development of the semi-empirical EDA was to better understand the interactions in protein-ligand complexes. For the initial benchmark study we chose the complex 5L87.[37,38] To evaluate the effects of truncation on the interaction energies and on the EDA, we performed calculations on the full protein and on the pocket. We furthermore split the pocket in two and performed the decomposition analysis of the resulting complexes. One of the pocket's halves was further partitioned in two fragments, on which we also ran calculations. The structures are available in section S7 and in Figure S6.1.

Expectedly, the magnitude of each energy contribution to the protein-ligand interaction increases with the number of protein atoms considered. Electrostatics differ significantly except for the two largest structures (protein and pocket). This is due to imbalances brought by the removal of residues in the most reduced versions of the pocket. We expect therefore that, if the ligand is charged or for pockets not reflecting the charge distributions in the protein, deviations could be more pronounced. Charge transfer is also impacted, though not visible in Figure S6.1 due to its minimal contribution - it accounts for less than 0.5% of the interaction energy. The weight of each contribution to the interaction energy also changes according to the number of atoms of the protein included. The change is also strongly dependent on the nature of the interactions probed in the calculation.

Using fragments to reconstruct a larger complex yields estimates of each contribution within 5% deviation. Such relative deviation decreases for larger blocks. However, absolute deviations increase: the addition of complexes 5 and 6 to reconstruct 4 (diamonds) yields better results than using 1 and 4 to rebuild 2 (circles). Dispersion and solvation are the contributions most affected by the cutoff, and in the case of the former the absolute deviation can go up to 3 kcal/mol. Deviations in the estimation of dispersion due to structural cutoffs are caused by missing pairwise interactions, but also due to the three body effects included in our calculations. This observation alone is enough to justify using the semi-empirical EDA to decompose protein-ligand interactions instead of splitting the complex into several small sub-blocks and to make use of higher-level methods.

Taking advantage of the atom-specific analysis at our hands, we built three-dimensional interaction maps (Figure S6.2), which indicate the relative contribution of different atoms to each term in the energy decomposition. These maps permit the classification of atoms according to attractive (blue) and repulsive (red) interactions with the protein, similar to what is done in SeeSAR's HYDE package.[54-56] Based on these, we determine that oxygen and nitrogen atoms dominate the attractive electrostatics with the protein. The Mulliken populations reveal that these are the atoms with lowest partial charge (*i.e.*, more electrons). The protein's pocket used for the calculation has a total charge of +6, which explains the results. On the other hand, repulsive electrostatics are dominated by the hydrogen atoms and the carboxyl's carbon. This is also easily understood when considering the Mulliken populations on the centers.

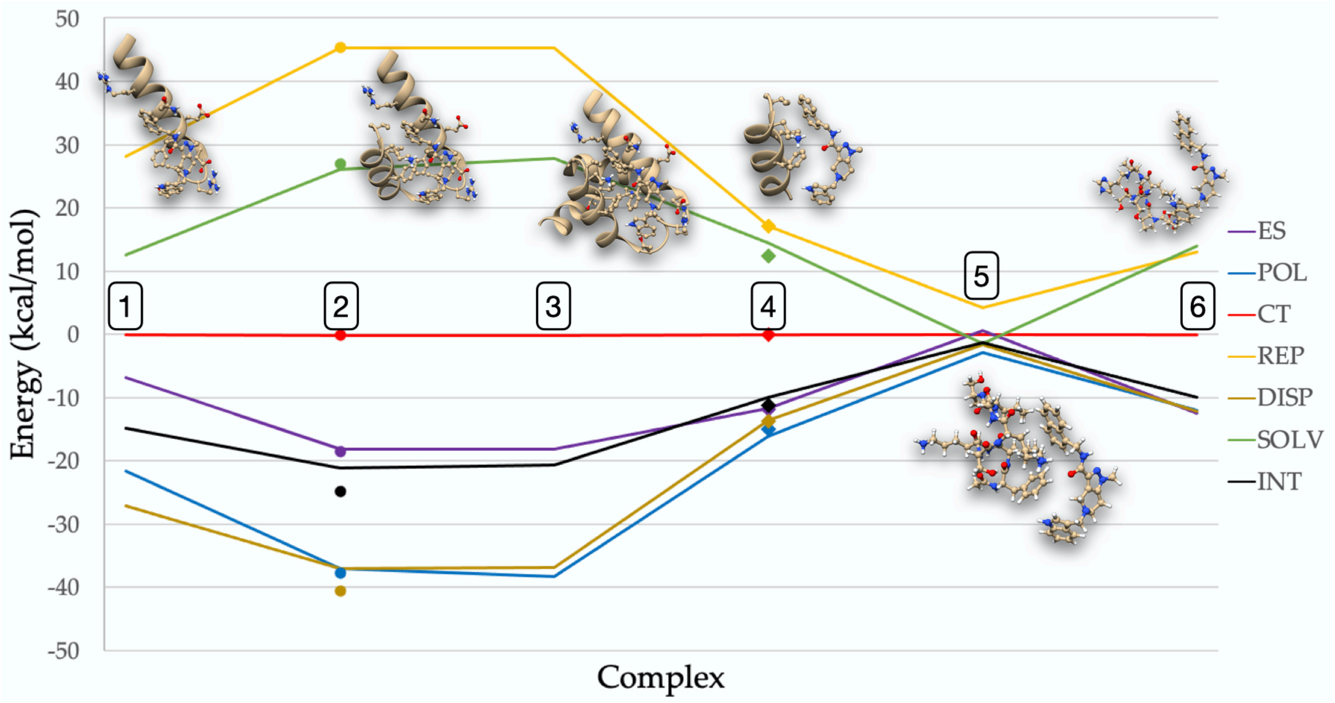


**Figure S6.1. Energy decomposition for several calculation boundary cutoffs of the same protein-ligand complex: half pocket A (1), pocket (2), full complex (3), half pocket B (4), quarter pocket BA (5) and quarter pocket BB (6). Vertices along the lines represent the results of the calculations, whereas markers alone stand for estimation of data based on subcomplexes.**

Stabilizing polarization interactions with the protein are mostly determined by protons, though some other atoms may have significant contributions too. An example would be the carbon from the methyl group in the ligand. The strongest polarization destabilization comes from the protons closest to the protein or deeper in the pocket. These are the atoms most susceptible to deformation of their electronic densities by the protein, shielding the rest of the ligand's atoms. Similarly, these centers contribute mostly to charge transfer. In the complex we are studying, charge transfer is overall attractive. A functional group that is particularly affected by charge transfer is the amide in the ligand, in specific the oxygen atom. The Mulliken populations are once more helpful in understanding the physics behind the observation: when the ligand binds to the pocket, the electronic population on this group decreases (oxygen: −0.55 *e* → −0.51 *e*), which leads to destabilization. The amide's nitrogen also suffers electronic depletion upon binding, the population is however less affected: −0.18 *e* → −0.17 *e*. Contrary to what happens to the amide's nitrogen and oxygen atoms, the carboxyl's carbon has a net gain of 0.004 electrons. This counteracts the charge depletion suffered by this atom due to its neighbors. Other non-hydrogen atoms that significantly participate in charge transfer are one of pyrazole's nitrogens and the methyl group bonded to the pyrazole. These have however opposing effects: pyrazole's nitrogen gains 0.01 electrons upon binding, which destabilizes the atom; the $C_{Me}$ has a loss of 0.04 electrons and is stabilized, as it goes closer to neutrality. Though the ligand has a net loss of 0.01 electrons when it goes into the protein's pocket, there is an exchange of almost 0.60 *e* between subsystems. If we take Reed *et al.*'s observations regarding charge transfer,[57] we verify that there are many effects cancelling each other on the complex.

Analysis of repulsion is identical to the analysis of polarization. This is again meaningful, since the atoms that should contribute most to stereo hindrance are the ones closest to the protein. A non-obvious atom that significantly contributes to the protein-ligand repulsion is the amide's oxygen atom. Interestingly, the nitrogen of this very same functional group has a negligible contribution. Comparing the CN bond distance in the 5L87 ligand against acetamide,[58] we observe that the bond is shorter in the former. The CO bond distance however increases in the ligand. This would indicate larger CN and smaller CO bond orders in the ligand than in acetamide. We stress that this is in very good agreement with the rest of the analysis done above.

Regarding the dispersion interactions, the dominant center is the naphthyl group. Though there is a phenyl ring from a nearby phenylalanine, this is not the main reason for naphthyl's dominance of this force because the two rings form a T-shape interaction. Comparing the dispersion interaction map with the binding pose shows that the strongest dispersion contributions come from the naphthyl centers penetrating deeper in the protein's pocket, as expected from the lipophilic nature of the interaction. This is particularly obvious in Figure S6.2 f). Second to the naphthyl ring is indole and only then the pyrazole ring. Finally, we have the solvation contributions. These are clearly dominated by the amide, in particular the oxygen atom, whose contribution goes against binding. The nitrogen atoms also have repulsive contributions, and the carboxyl's carbon gives a positive impact.

Summing up all contributions shows that key to binding is the ligand's amide group (mostly electrostatics), the naphthyl (mainly by dispersion) and the external hydrogen atoms on the indole ring (mainly due to polarization). Dispersion on the indole and pyrazole rings is also not to neglect. However, such an effect is mostly cancelled. The pyrazole ring also offers a good interaction from one of the nitrogen atoms, which rivals in magnitude one the naphthyl's carbons. Nevertheless, in terms of intensity, this is of course reduced due to the number of atoms in each functional group. In fact, we performed a functional group based analysis and we observed that the naphthyl substituent binds stronger to the protein than the tetrahydro-pyrazolopyridine core by almost 5 kcal/mol. Among other factors, this is determined by the strength of dispersion forces and a positive impact in removing the group from water's dielectric medium.

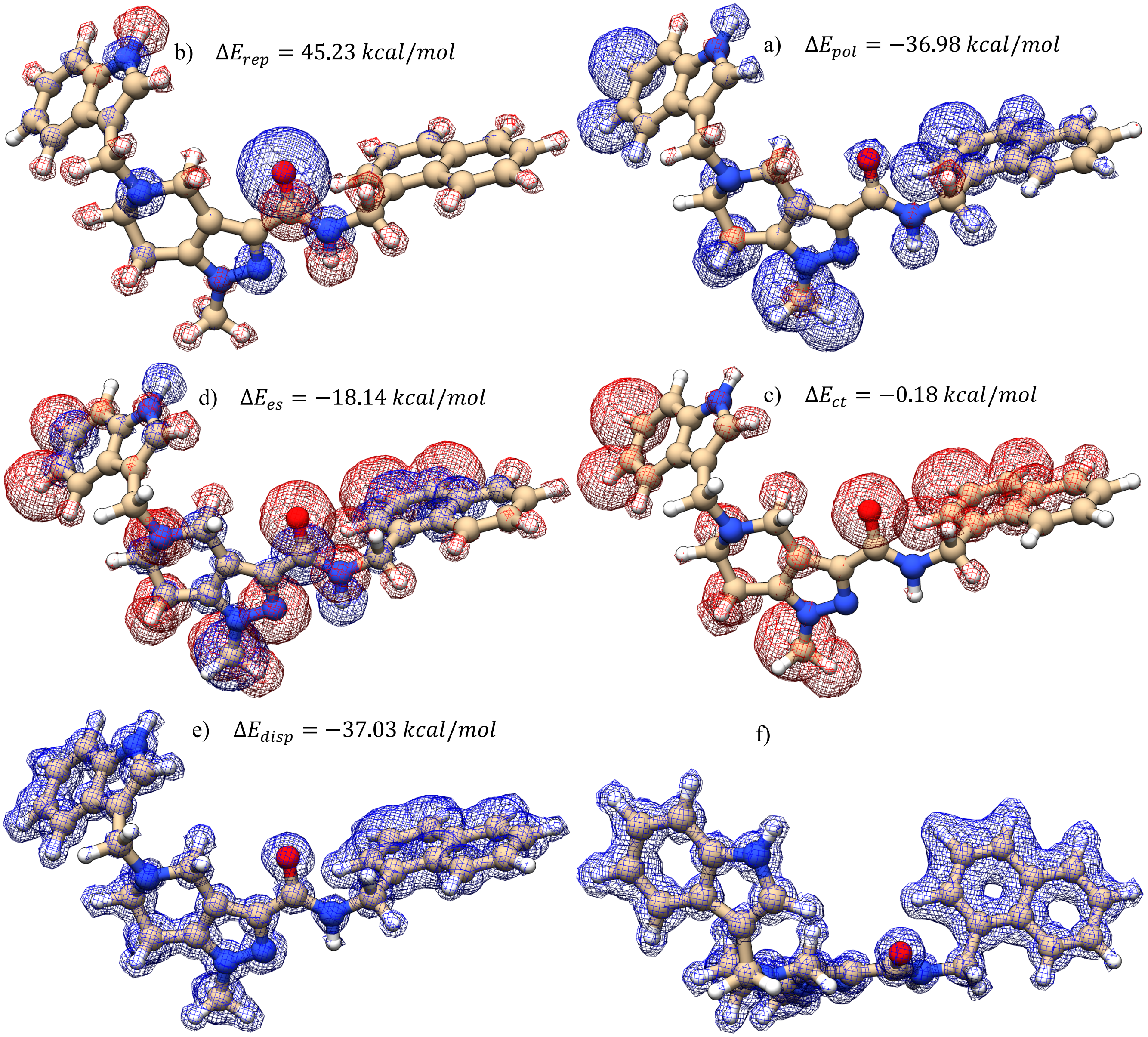

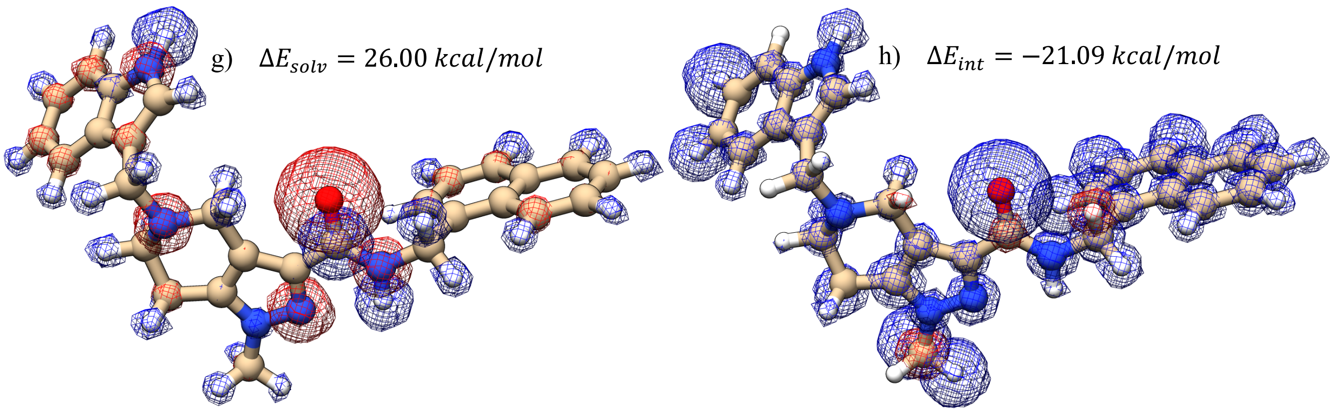


**Figure S6.2. Interaction maps for the ligand in 5L87 with the respective protein: a) electrostatics; b) polarization; c) charge transfer; d) repulsion; e) dispersion, view 1; f) dispersion, view 2; g) solvation; h) total interaction. Red represents repulsion, whereas blue stands for attractive forces.**

At this point it is important to mention that considering reduced versions of the protein's pocket may have a profound impact on the interaction maps described above. For instance, using the complex 1 to estimate interactions leads to an incomplete electrostatic and polarization picture (missing key interactions) and a completely wrong dispersion map: the naphthyl, indole and pyrazole rings become all equivalent in strength. Similar observations take place for the analysis based on the pocket subcomplex 4.

When we perform the EDA on the full protein-ligand complex we observe that there are minimal changes in the magnitude of the different contributions to the interaction energy. In fact, even the total binding energy differs by less than 0.5 kcal/mol. Repulsion is, expectedly, unaffected, since the additional blocks of the protein are not in direct contact with the ligand. The same applies to charge transfer, though this is so minimal in the complex that any change in this contribution has virtually no impact. The effect on electrostatics is, in a first instance, unexpectedly minimal. A closer look at the structures reveals that in the full protein complex there is essentially extension of the scaffold, and the formally charged groups participate in ionic bridges. In all due fairness, the pocket we cut is not much smaller than the full protein. On the other hand, the solvation contribution increases by almost 2.0 kcal/mol, which is compensated by a similar decrease in polarization. The former may be justified by the change in the exposed surface from the protein. The latter is a cumulative effect of the charge distribution of the protein on the ligand. Dispersion is also barely impacted.

We note that this study did not include any explicit molecule of water. As is seen in the main document, this may contribute to accentuate differences caused by structural truncation. Before we close this section, we feel it is still important to comment on the timings involved in these calculations. Using a personal computer (iMac 3.6 GHz, quad-core i7) and running the calculations in serial (our ULYSSES package is still not parallelized), it required 15 minutes to run the EDA on the pocket and 50 minutes the EDA on the full protein complex. These timings will reduce significantly once we make the code parallel and linear scaling techniques are introduced.

### S7 – Enlargement of protein-ligand structures

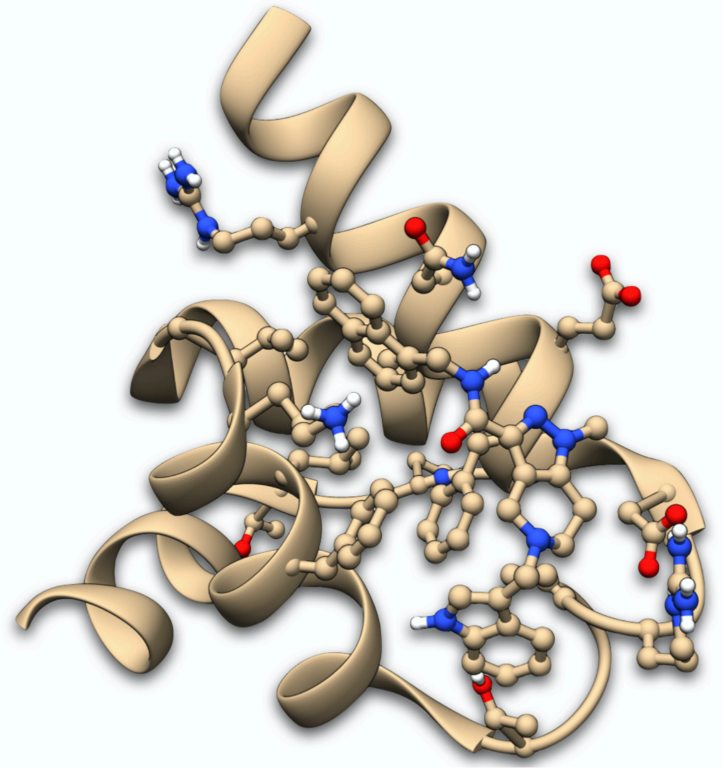

**Figure S7.1. Complex 3, full protein-ligand.**

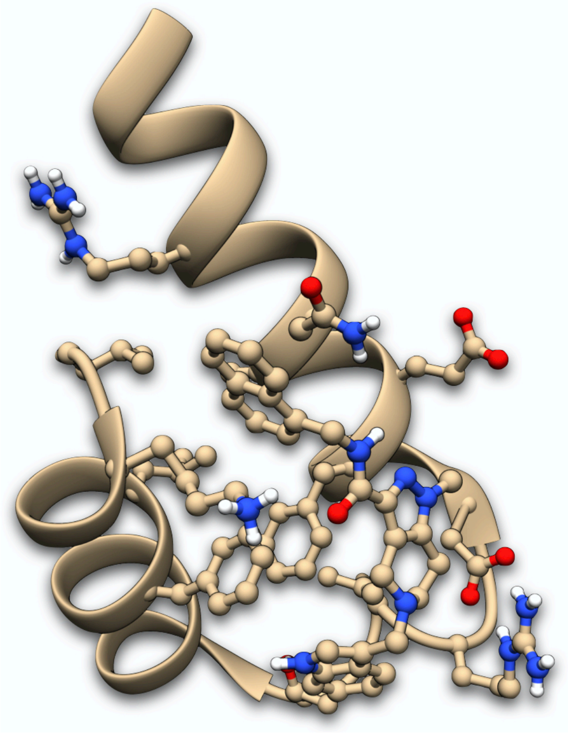

**Figure S7.2. Complex 2, ligand in protein's pocket.**

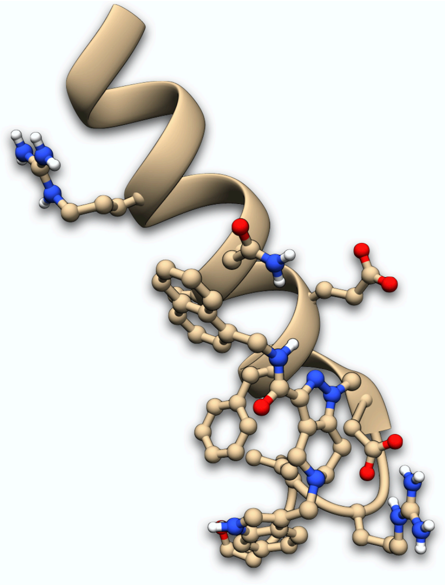

**Figure S7.3. Complex 1, pocket half A.**

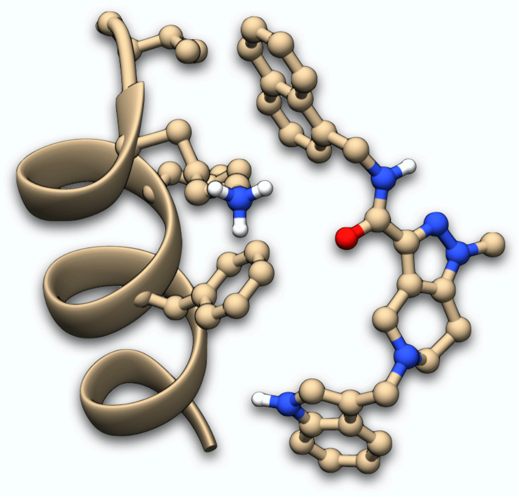

**Figure S7.4. Complex 4, pocket half B.**

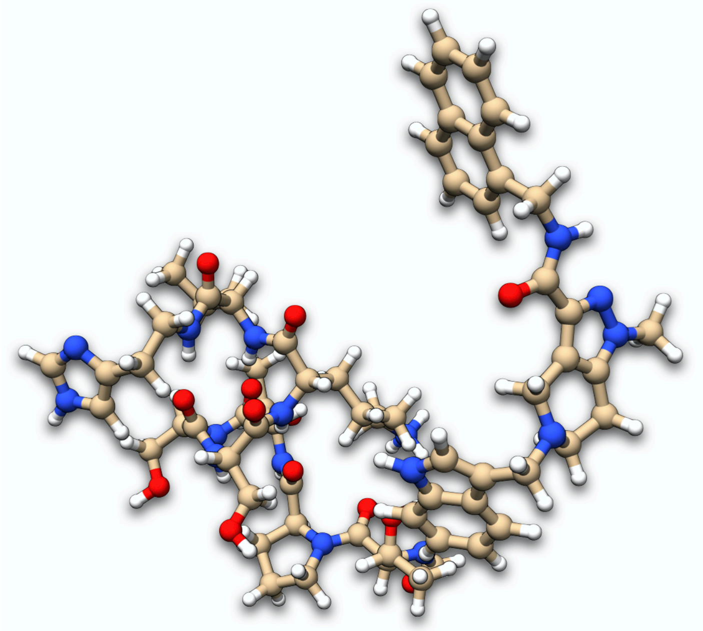

**Figure S7.5. Complex 5, pocket quarter BA.**

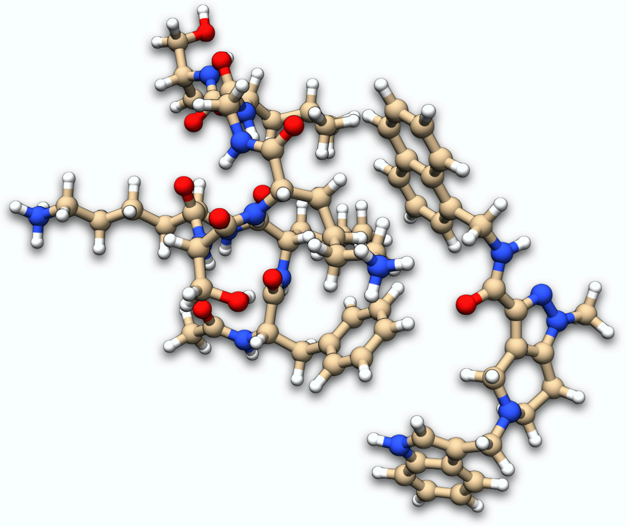

**Figure S7.6. Complex 6, pocket quarte BB.**

### S8 – Redefinition of matrices in the van der Vaart-Merz notation using block-explicit density matrices

The electrostatic state is obtained by comparing the reference state (isolated molecules) against the system at the equilibrium distance using the density of the isolated molecules. This is thus given by

$$\Delta E_{es} = E\left(R_{AB}, \begin{bmatrix} AA(+\infty) & AB(+\infty) \\ BA(+\infty) & BB(+\infty) \end{bmatrix}\right) - E\left(+\infty, \begin{bmatrix} AA(+\infty) & AB(+\infty) \\ BA(+\infty) & BB(+\infty) \end{bmatrix}\right)$$

In the expression above, $R_{AB}$ is the distance between monomers in the supermolecule. The density matrix for the system is made explicit in terms of blocks describing either monomer (AA and BB) or the interaction between these. Since we use the density of the separated monomers ($+\infty$), then the AB and BA blocks are necessarily zero and the density used in the calculation above is block diagonal. The polarization and charge transfer states are determined using

$$\Delta E_{pol} = E\left(R_{AB}, \begin{bmatrix} AA(R_{AB}) & AB(+\infty) \\ BA(+\infty) & BB(R_{AB}) \end{bmatrix}\right) - E\left(R_{AB}, \begin{bmatrix} AA(+\infty) & AB(+\infty) \\ BA(+\infty) & BB(+\infty) \end{bmatrix}\right)$$

and

$$\Delta E_{ct} = E\left(R_{AB}, \begin{bmatrix} AA(R_{AB}) & AB(R_{AB}) \\ BA(R_{AB}) & BB(R_{AB}) \end{bmatrix} - E\left(R_{AB}, \begin{bmatrix} AA(R_{AB}) & AB(+\infty) \\ BA(+\infty) & BB(R_{AB}) \end{bmatrix}\right)\right)$$

Note that AB($R_{AB}$) and BA($R_{AB}$) are no longer block diagonal. In our algorithm, because constrained-SCF is used to calculate the polarization state, then the terms AB($+\infty$) and BA($+\infty$) are not necessarily zero, *i.e.*, the polarization density is not necessarily block diagonal.

### S9 – Definition of Matrices

$$\mathbf{H}^{int} = \mathbf{H}(R_{AB})$$

$$\mathbf{H}^{0} = \mathbf{H}(R_{AB} = +\infty)$$

$$\mathbf{F}^{0} = \mathbf{F}(R_{AB} = +\infty, \mathbf{P}^{0})$$

$$\mathbf{F}^{es} = \mathbf{F}(R_{AB}, \mathbf{P}^{0})$$

$$\mathbf{F}^{pol} = \mathbf{F}(R_{AB}, \mathbf{P}^{pol})$$

$$\mathbf{F}^{int} = \mathbf{F}(R_{AB}, \mathbf{P}^{int})$$

$$\mathbf{P}^{0} = \begin{bmatrix} AA(+\infty) & AB(+\infty) \\ BA(+\infty) & BB(+\infty) \end{bmatrix} = \begin{bmatrix} AA(+\infty) & 0 \\ 0 & BB(+\infty) \end{bmatrix}$$

$$\mathbf{P}^{pol} = \begin{bmatrix} AA(R_{AB}) & AB(+\infty) \\ BA(+\infty) & BB(R_{AB}) \end{bmatrix} = \begin{bmatrix} AA(R_{AB}) & 0 \\ 0 & BB(R_{AB}) \end{bmatrix}$$

$$\mathbf{P}^{int} = \begin{bmatrix} AA(R_{AB}) & AB(R_{AB}) \\ BA(R_{AB}) & BB(R_{AB}) \end{bmatrix}$$

Note that the definition used above for the polarization density is valid in the van der Vaart-Merz notation, and not when constrained SCF if used.

### S10 – Constrained SCF

When trying to find wavefunctions for a system, such that electronic populations are subject to some constraint, constrained Self-Consistent Field (SCF) is perhaps the most widely used algorithm.[10] In the case of our Energy Decomposition Analysis (EDA) program, we wish to find a relaxed wavefunction, such that the electronic populations of two monomers are the same as if the molecules were isolated, *i.e.*, without any sort of charge transfer. The total number of electrons on each monomer must then be $N_A^0$ and $N_B^0$. We need thus to find a set of MOs, a Lagrange multiplier λ and a set of weights $w_A$ and $w_B$ such that

$$E_{SCF}^{\lambda} = E_{SCF} + \lambda\left[tr\left(\mathbf{P}\mathbf{w}_A\right) - N_A^0\right] - \lambda\left[tr\left(\mathbf{P}\mathbf{w}_B\right) - N_B^0\right]$$

is a stationary point. Note that we seek a stationary point because the energy of the system is minimized with respect to the set MOs but maximized with respect to the constraint defined by the Lagrange multiplier. This is easily grasped, since the natural state of the system will necessarily include charge transfer, which is here inhibited.

Differentiating the equation above yields a modified Fock matrix

$$\mathbf{F}^{\lambda} = \mathbf{F} + \lambda\left(\mathbf{w}_A - \mathbf{w}_B\right)$$

which must be used to obtain the respective set of MOs, under the above specified constraints.

In solving this problem, we follow the algorithm proposed by Wu and Van Voorhis.[10,11] Here, the SCF procedure is split in two cycles: a set of macro-iterations, which is identical to the typical SCF procedure; the micro-iterations, in which the optimal (maximum) value for λ is to be found. Like suggested,[10] we use the Newton method to obtain the solution to the micro-iterations. This requires the first and second derivatives of $E_{SCF}^{\lambda}$ with respect to λ:

$$\frac{\partial E_{SCF}^{\lambda}}{\partial \lambda} = 2\left[tr\left(\mathbf{P}\mathbf{w}_A\right) - N_A^0\right]$$

The factor of 2 in the equation above comes from the fact that the electrons we remove from one system are given to the other. This is because our constrained-SCF is specifically suited for charge transfer related problems. The second derivatives are given by

$$\frac{\partial^2 E_{SCF}^{\lambda}}{\partial \lambda^2} = 4\sum_{i\in occ}\sum_{a\in vir}\frac{A_{ia}^2}{\epsilon_i - \epsilon_a}$$

where

$$\mathbf{A} = \mathbf{C}^{o\dagger}\left(\mathbf{w}_A - \mathbf{w}_B\right)\mathbf{C}^{v}$$

In the expressions above, $\epsilon$ are the MO energies, $\boldsymbol{C}^o$ is the occupied block of the MO coefficient matrix and $\boldsymbol{C}^v$ is the respective virtual block. Despite the decoupling, solving the SCF allows one to find the optimal set of orbitals for a fixed λ. At the end of each iteration, one ends up with a set of constrained MOs and the respective ideal Lagrange multiplier.

To apply the above-described algorithm, one still requires the weights $\boldsymbol{w}_A$ and $\boldsymbol{w}_B$. The fact that we wish to restrict electronic populations on specific parts of the system suggests that these two quantities are somehow related to population analysis. Though we presented an energy expression using Mulliken populations, other schemes are also possible.[10,11] However, since our focus is semi-empirical EDA, we defined our algorithm exclusively based on the Mulliken electronic populations of each monomer. This yields

$$w_{A,\mu\nu} = \begin{cases} S_{\mu\nu}, & \mu, \nu \in A \\ \frac{1}{2} S_{\mu\nu}, & \mu \in A \quad \vee \quad \nu \in A \\ 0, & \mu, \nu \notin A \end{cases}$$

and a similar expression applies to $w_B$. In the case of GFN2-xTB, we may set additionally the middle term of the expression above to zero in order to get spatially restricted polarization states.

Typical constrained SCF algorithms require the user to define a set of starting values for the Lagrange multipliers (λ in our case) or they simply use default values. To optimize the efficiency of the Newton steps and to make the algorithm black-box like, we use the bisection method to make an initial search of starting values for λ. Also, for robustness, we use the bisection step, whenever the Newton one overshoots or is expected to be qualitatively poor. Though this choice increases the initial overhead (due to the bisection search), it effectively minimizes the number of bisection steps during the rest of the calculation, which is important in protein calculations. Furthermore, in most cases, we observe that the number of micro-iterations within each macro-iteration is typically very small, around 2-3. To adapt the initial bisection search, we start by scouting in both directions from $\lambda = 0$, *i.e.*, we gradually modify λ for positive and negative values. Using the derivative $\partial E_{SCF}^{\lambda} / \partial \lambda$, we determine which direction has the potential change of sign. After the first bisection search iteration, this becomes the only scouting direction.

### S11 – Conceptual Problems in the Calculation of Electrostatics with NDDO Methods

It has been pointed that the EDA of van der Vaart and Merz, coupled with semi- empirical methods, yields positive electrostatic interactions.[20] Due to the disagreement with other theoretical data,[59] this and other results led researchers to state that NDDO methods are very poor in describing electrostatics.[60] Though of course semi-empirical charges lack the quality of *ab initio* ones, we believe that the panorama offered by semi-empirical methods is not as dire. As mentioned above, the results of the van der Vaart-Merz EDA are, despite the limitations of NDDO methods to describe electrostatics, a result of a theoretical inconsistency.

Electrostatic interactions should be formally calculated when the AO-monomer overlap is inexistent.[7] By construction, NDDO methods consider that the AO-basis of the quantum chemical system is orthonormal, though in reality there are (significant) deviations. If for instance one calculates the wavefunctions for the monomers at infinite distance and separately, *i.e.*, isolated from one another, leads technically to two theoretically inconsistent states. This is because in the former the AOs of the monomers are orthogonal to one another, whereas in the latter they are independent and technically potentially non-orthogonal. The results are however numerically indistinguishable because the overlap between the AOs of the monomers zeroes due to the very large distance. The practical impact of this observation takes place when calculating the electrostatic state defined in the main text. This is because the charge distributions of the monomers are obtained in a context that retains the formal orthogonality of the AO-bases, however, by construction, the zeroth-order approximation to the binding energy considers that the orbitals from the monomers are orthogonal. When we calculate the electrostatic state using NDDO methods, by default construction of these, overlap effects are taken in consideration. The repulsive electrostatics obtained with these methods are then a theoretical inconsistency instead of a limitation of the methods.[59]

### S12 – Interaction Maps for Protein-Ligand Complexes

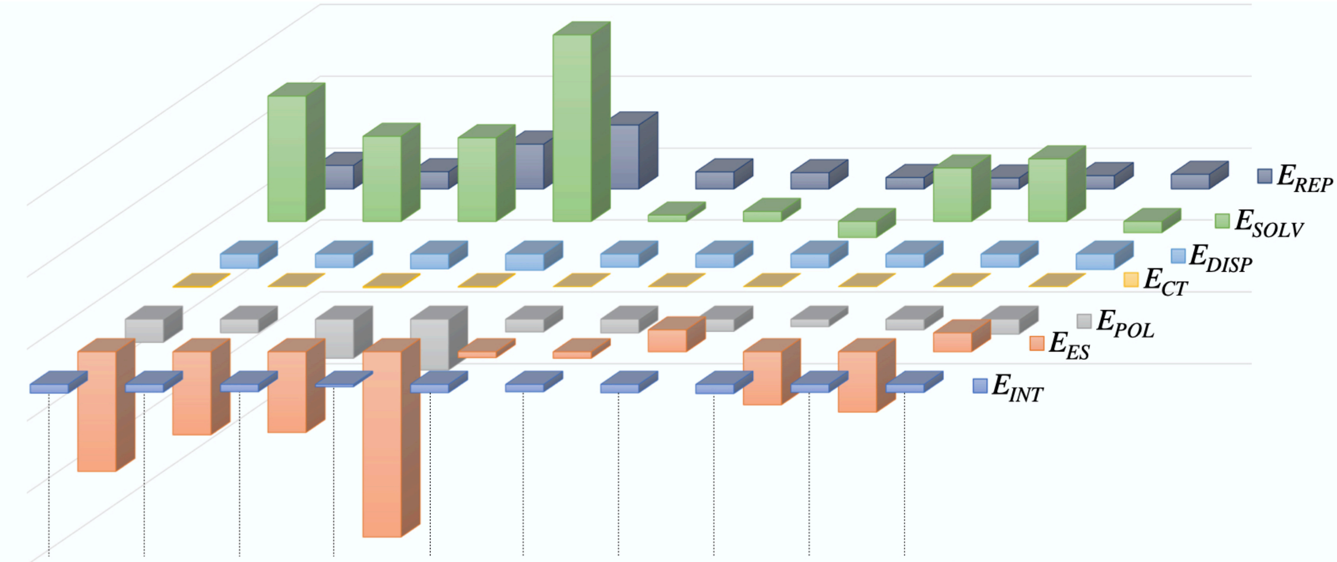


| 2W84 | 2W84’ | 2W85 | 4BXU | 5L8A | 5L87 | 5N8V | 5OML | 6RT2 | 6SPT | |
|---|---|---|---|---|---|---|---|---|---|---|
| 65.35 | 47.77 | 124.42 | 177.71 | 47.38 | 45.26 | 31.75 | 30.42 | 37.02 | 40.73 | $E_{REP}$ |
| 348.89 | 236.97 | 232.88 | 519.17 | 17.66 | 27.75 | -44.36 | 148.63 | 174.26 | -31.27 | $E_{SOLV}$ |
| -39.51 | -37.35 | -41.09 | -45.29 | -36.42 | -36.96 | -37.93 | -36.41 | -36.27 | -42.80 | $E_{DISP}$ |
| -1.82 | -0.07 | -3.10 | -1.39 | -0.08 | -0.18 | -0.01 | 0.04 | 0.00 | -0.06 | $E_{CT}$ |
| -64.53 | -37.87 | -108.78 | -141.41 | -35.32 | -38.37 | -33.82 | -20.72 | -29.45 | -41.43 | $E_{POL}$ |
| -332.85 | -231.13 | -224.64 | -515.46 | -16.48 | -18.13 | 60.94 | -147.51 | -168.03 | 52.45 | $E_{ES}$ |
| -24.49 | -21.69 | -20.32 | -6.67 | -23.25 | -20.63 | -23.44 | -25.55 | -22.46 | -22.38 | $E_{INT}$ |

**Figure S12.1. Energy decomposition for the protein-ligand and protein-peptide complexes considered in this work. The embedded table gives the respective contributions in units of kcal/mol.**

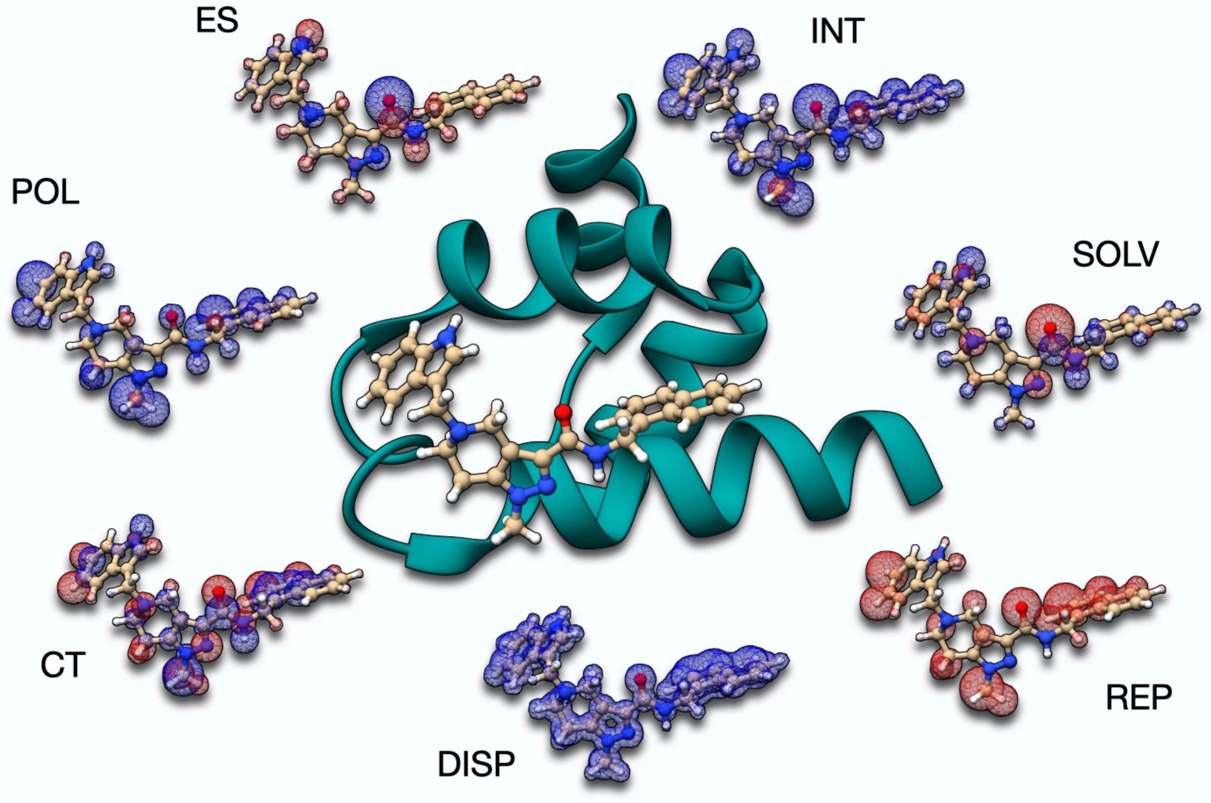


**Figure S12.2. Interaction maps for 5L87. Blue stands for attractive interactions, whereas red stands for repulsion.**

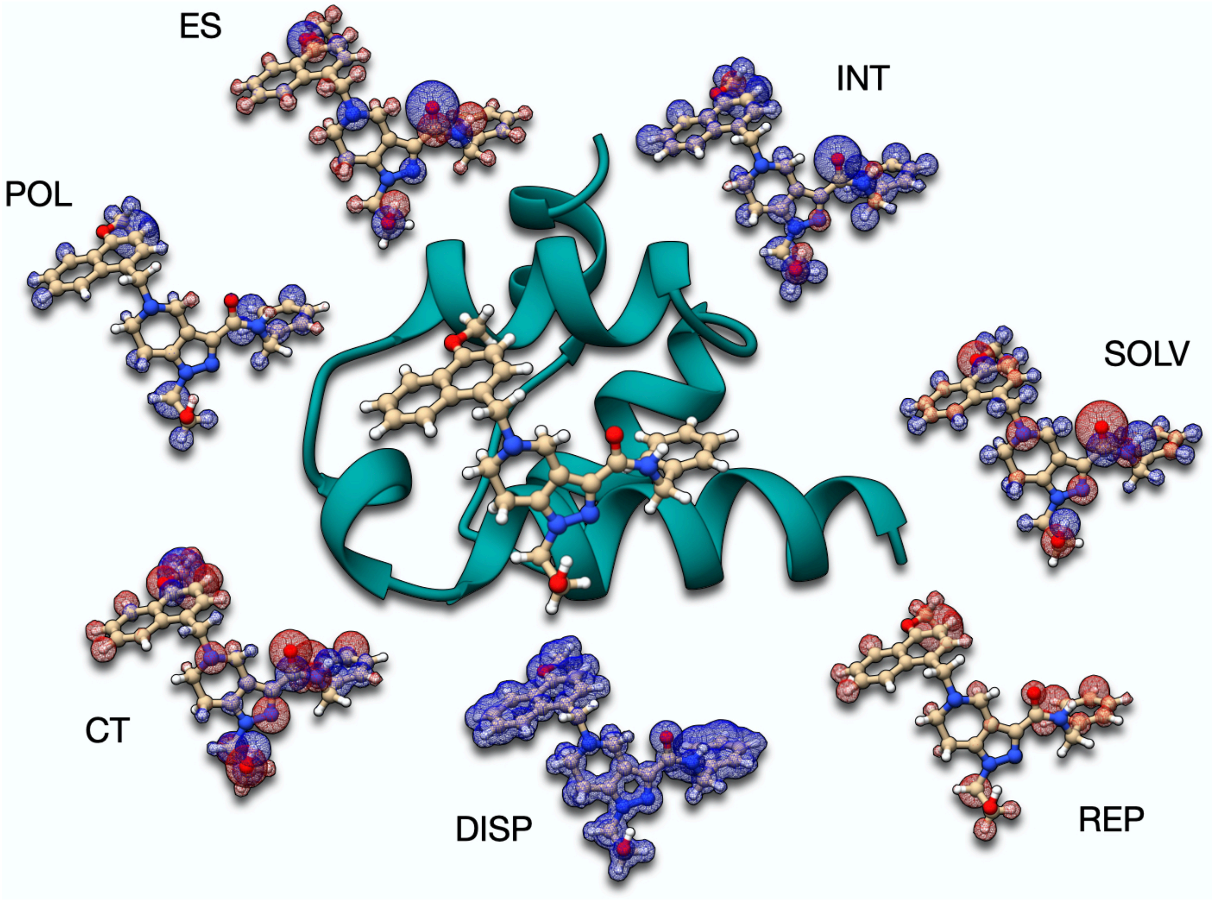

**Figure S12.3. Interaction maps for 5L8A. Blue stands for attractive interactions, whereas red stands for repulsion.**

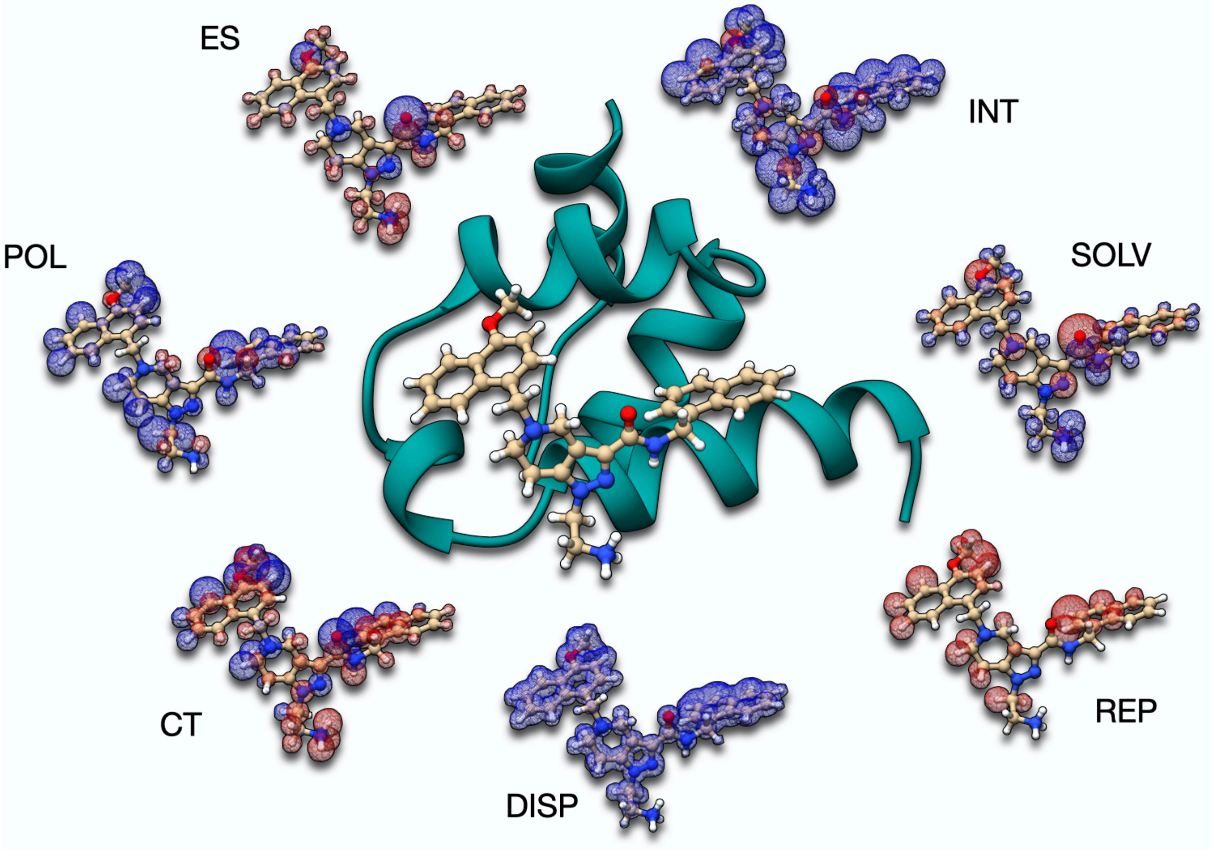

**Figure S12.4. Interaction maps for 5N8V. Blue stands for attractive interactions, whereas red stands for repulsion.**

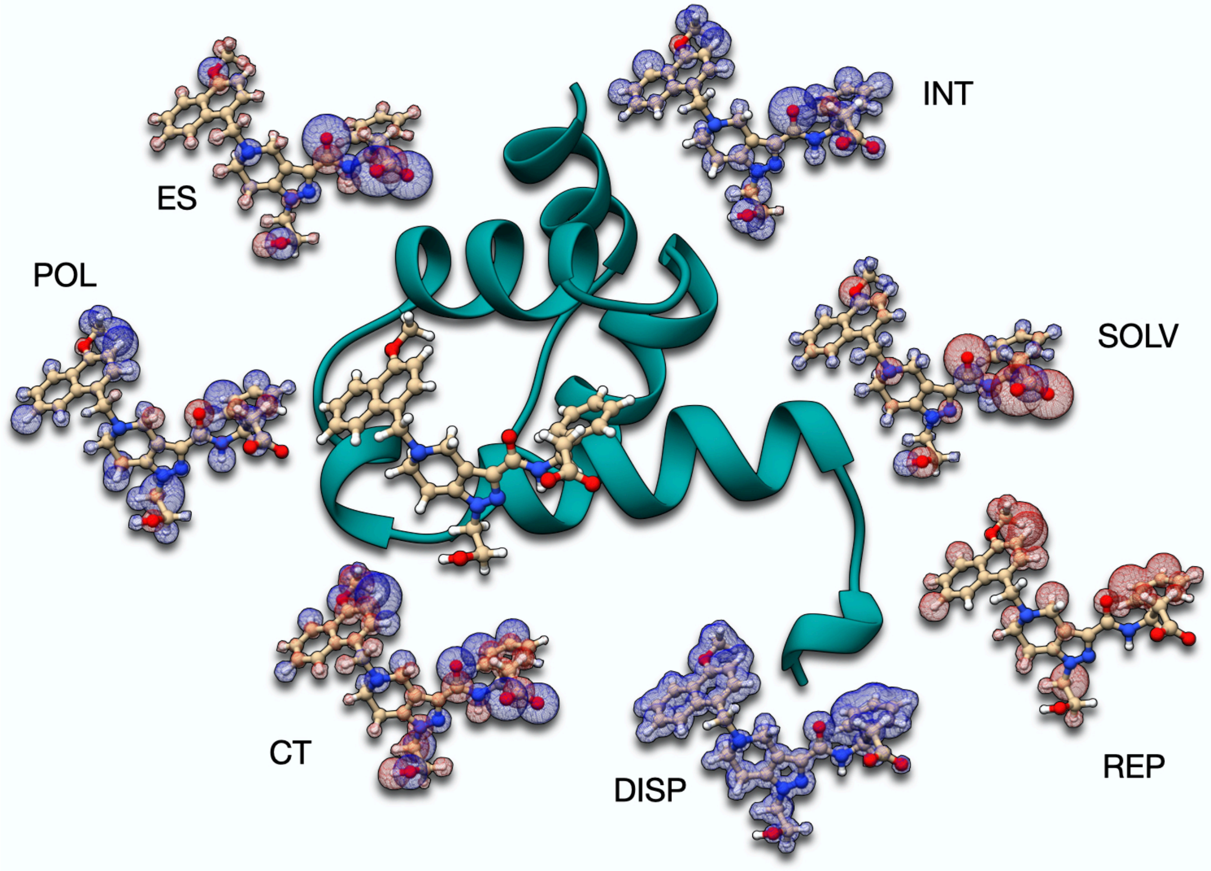

**Figure S12.5. Interaction maps for 5OML. Blue stands for attractive interactions, whereas red stands for repulsion.**

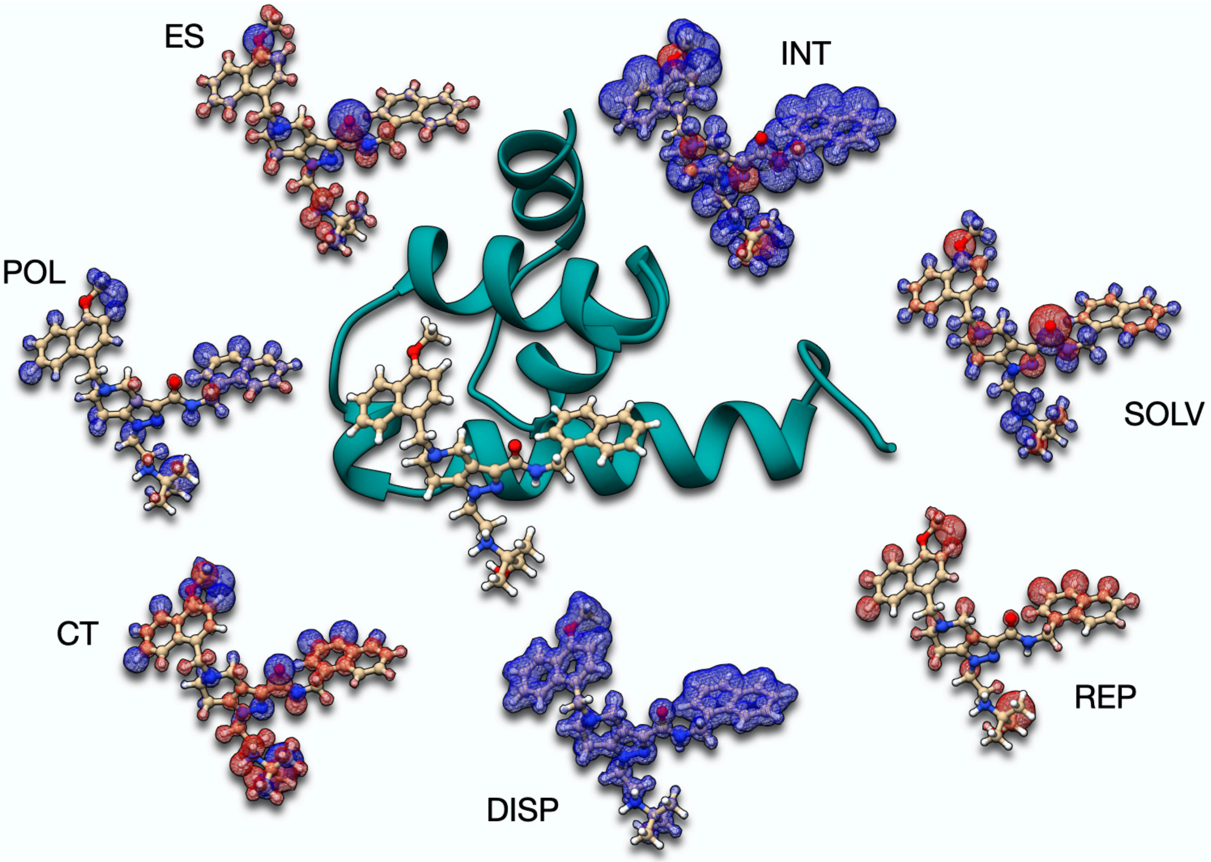

**Figure S12.6. Interaction maps for 6SPT. Blue stands for attractive interactions, whereas red stands for repulsion.**

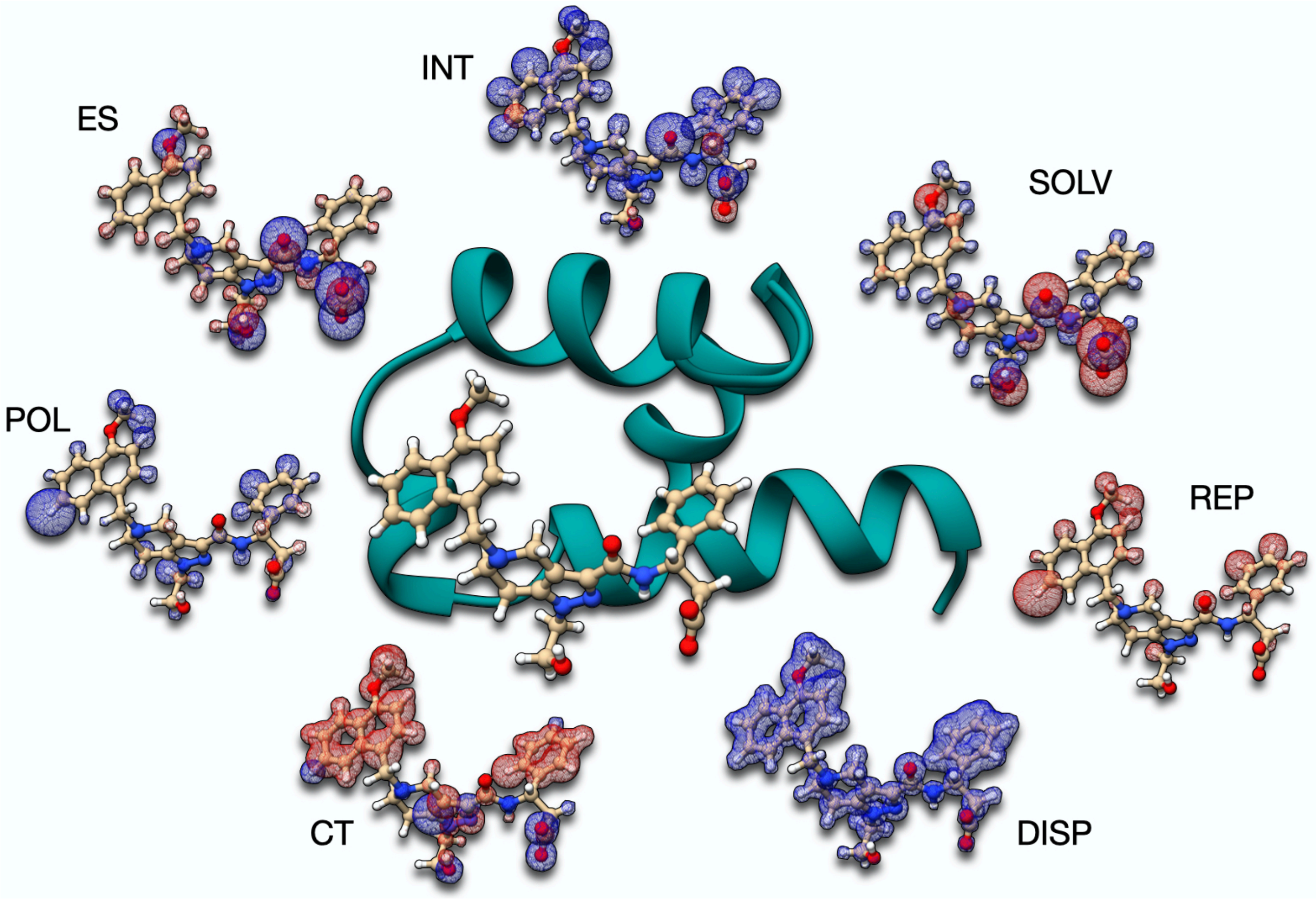


**Figure S12.7. Interaction maps for 6RT2. Blue stands for attractive interactions, whereas red stands for repulsion.** We stress that the red surface in the charge transfer maps is exacerbated due to the equivalently small contribution from aromatic rings.

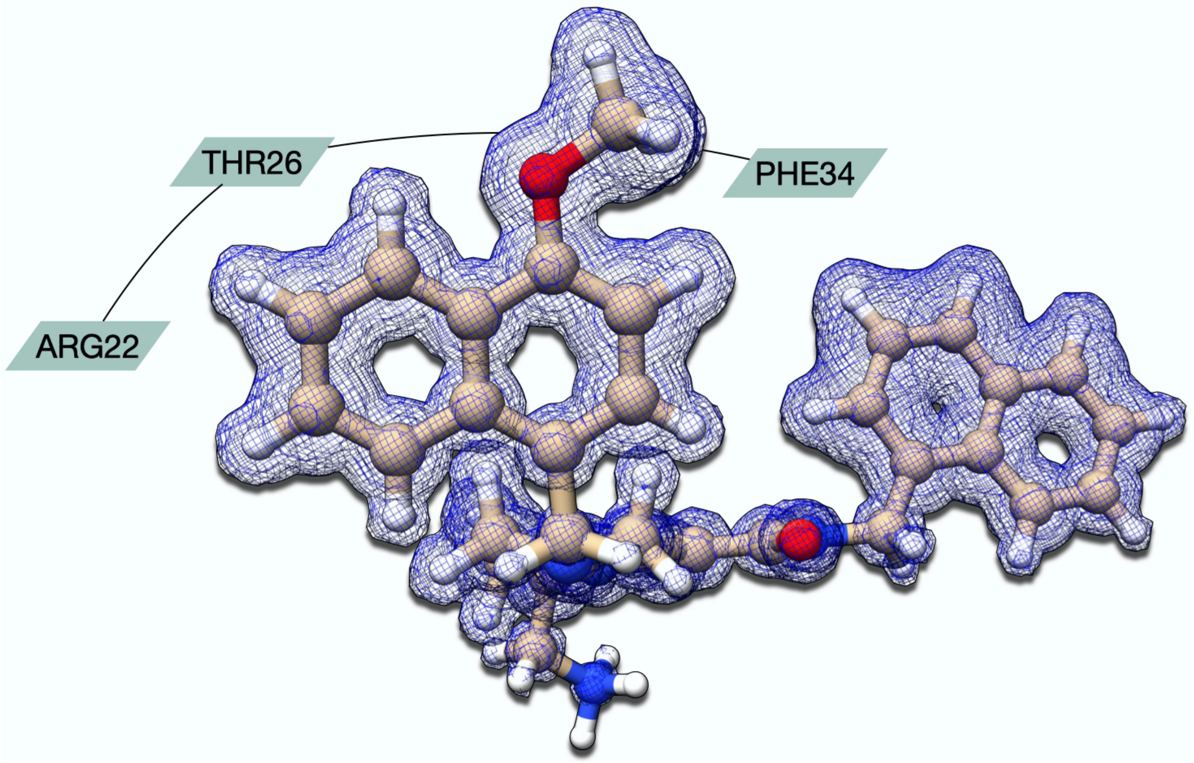


**Figure S12.8. Dispersion Maps for 5N8V with a special focus on the direct residue interactions surrounding the methoxynaphthyl group.**

**Of the structures with inhibitors, complexes with charge neutral ligands, like 5L87 and 5L8A, show slightly stabilizing electrostatics. On the other hand, for negatively charged ligands, like 5OML and 6RT2, ES contributions are strongly attractive (Figure 3). This last case is particularly interesting because the two ligands are enantiomers. Nevertheless, the EDA calculations expect significantly different ES stabilizations. Curiously, the stronger binder shows weaker ES stabilization (*c.f.* discussion below). For positively charged inhibitors, 5N8V and 6SPT, ES contributions are already repulsive. For 5N8V we also considered the neutral form of the ligand, and in this case, electrostatics become attractive. The magnitude of the stabilization is on par with the other charge neutral ligands herein studied, *i.e.*, 5L87 and 5L8A.**

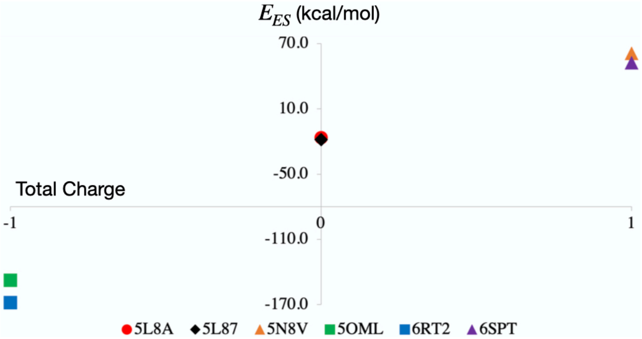


**Figure S12.9. ES contributions (in kcal/mol) for the ligands considered as a function of the ligand's total charge.**

Of the structures with inhibitors, complexes with charge neutral ligands, like 5L87 and 5L8A, show slightly stabilizing electrosta

### S13 – Interaction Maps for Protein-Protein Complexes

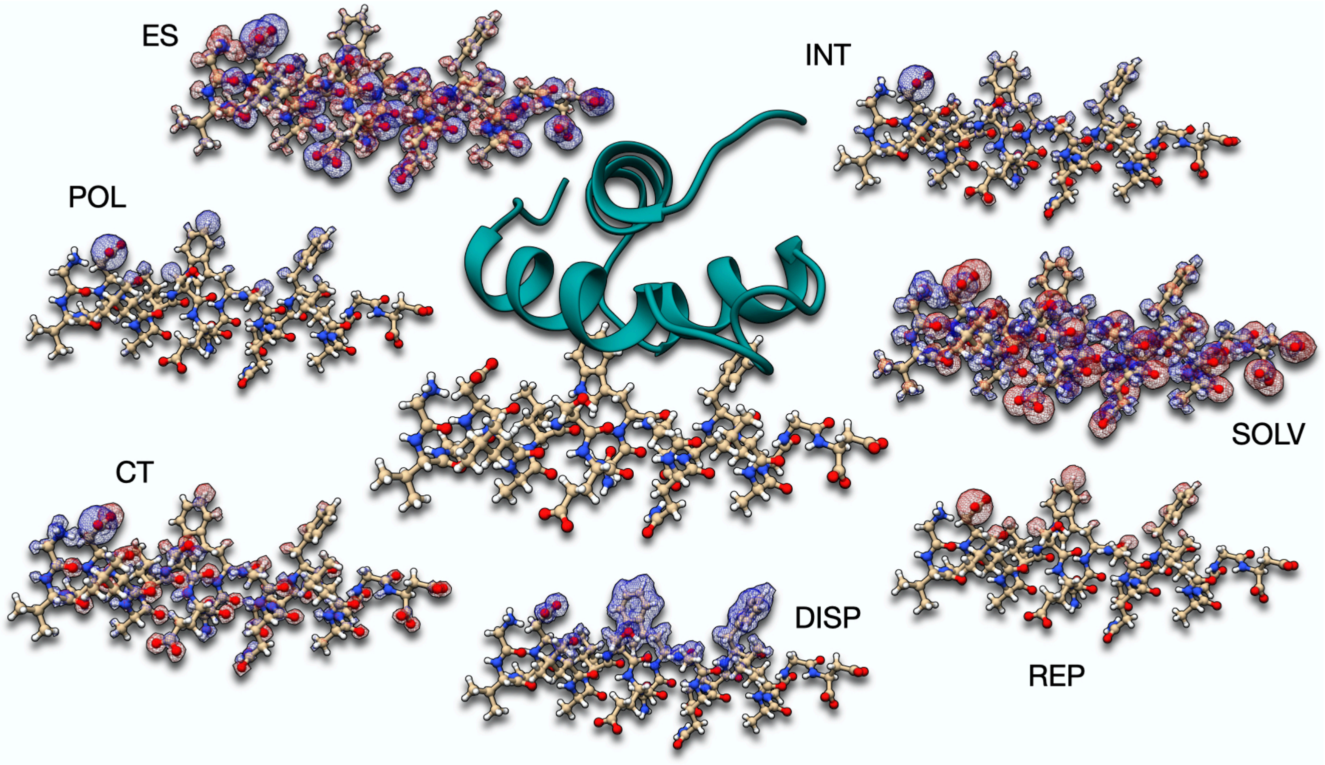


**Figure S13.1. Interaction maps for 2W84. Blue stands for attractive interactions, whereas red stands for repulsion.**

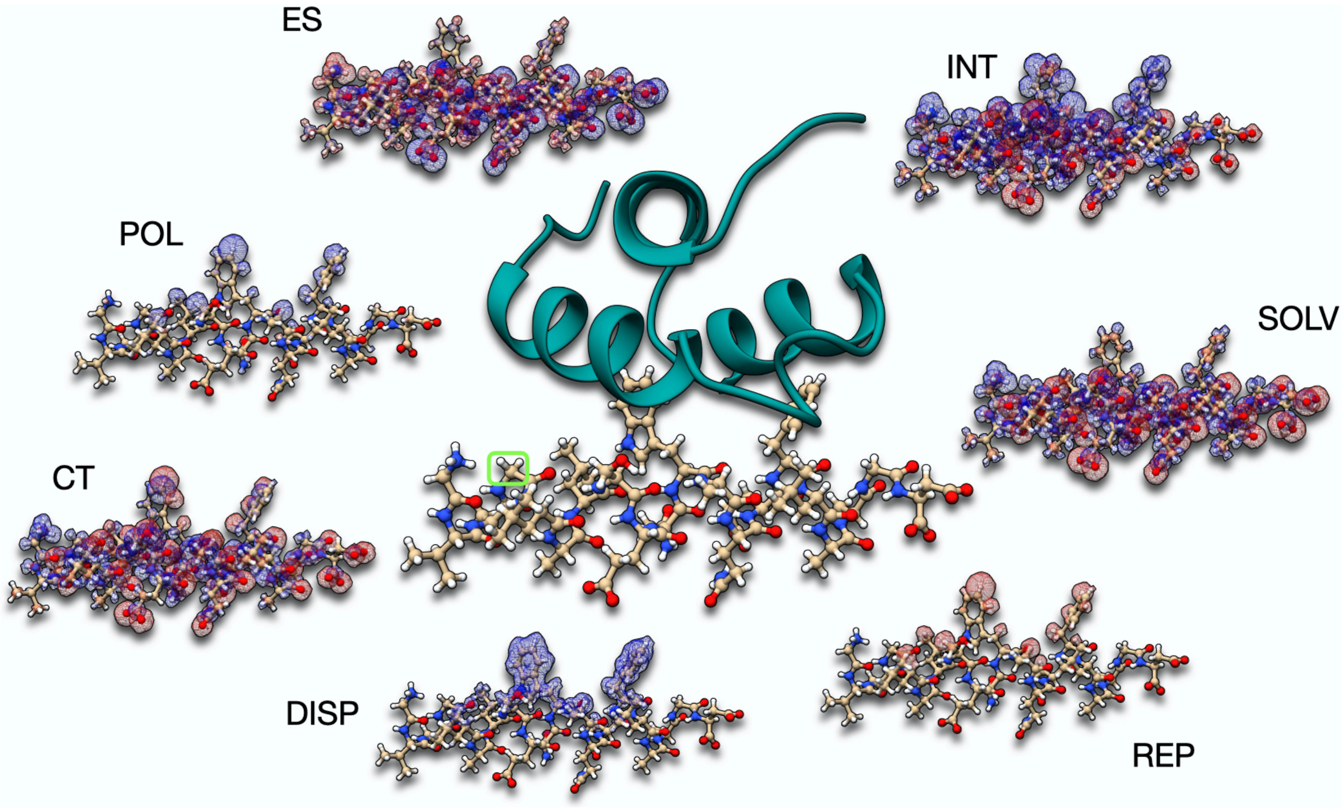


**Figure S13.2. Interaction maps for 2W84'. Blue stands for attractive interactions, whereas red stands for repulsion.**

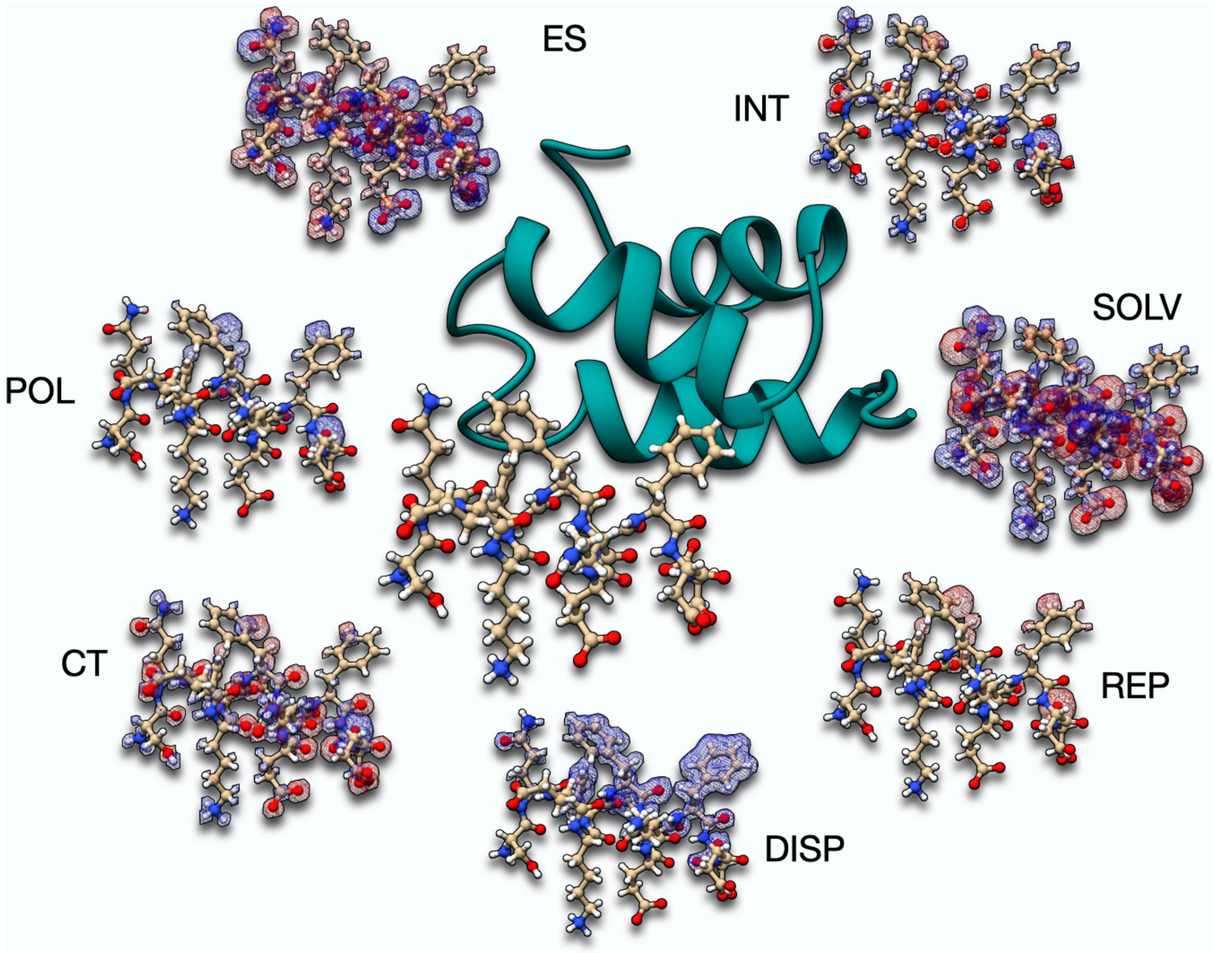


**Figure S13.3. Interaction maps for 2W85. Blue stands for attractive interactions, whereas red stands for repulsion.**

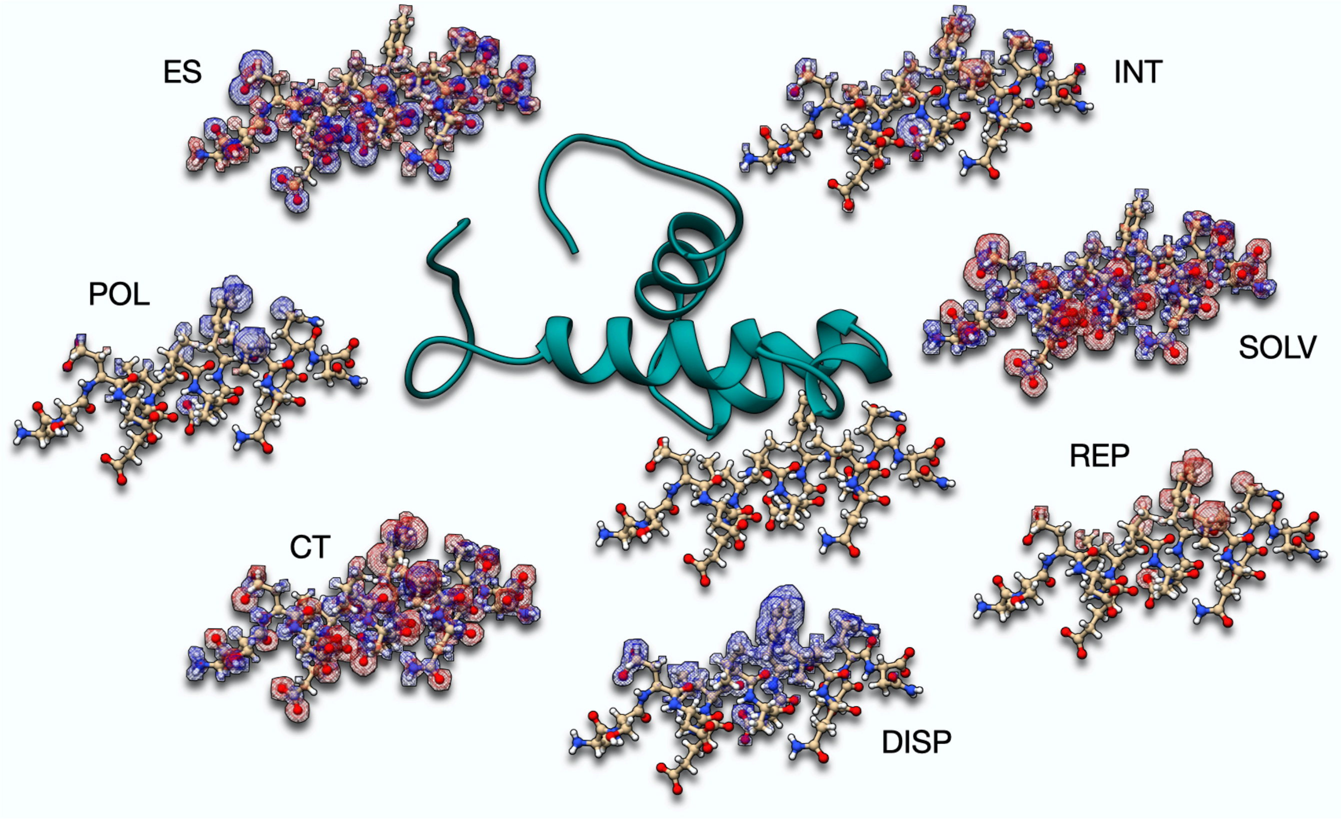


**Figure S13.4. Interaction maps for 4BXU. Blue stands for attractive interactions, whereas red stands for repulsion.**

### S14 – Energy Decomposition Analysis for different poses of 5N8V

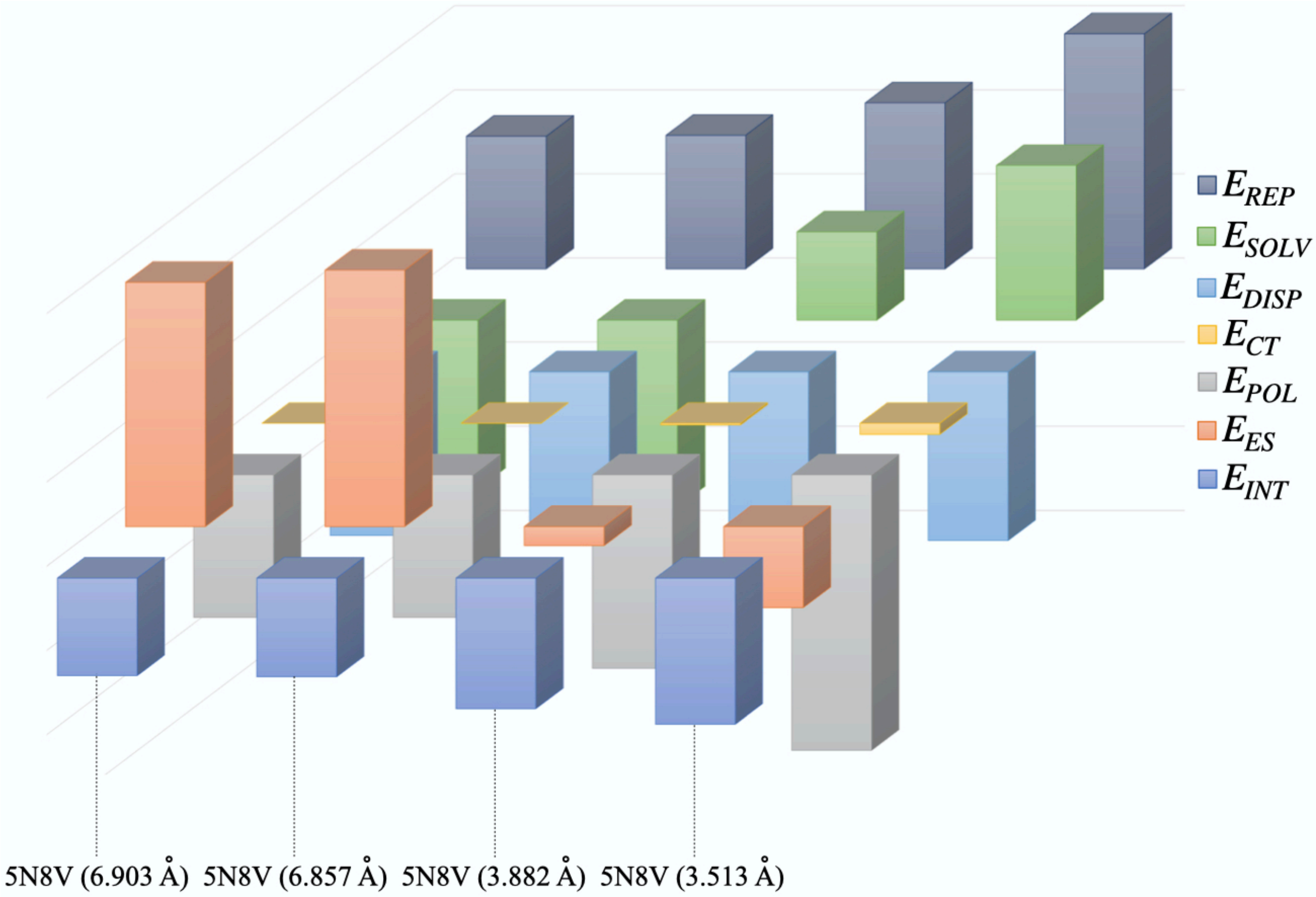


**Figure S15.1. Column plot for the EDA calculations on different poses of 5N8V.**

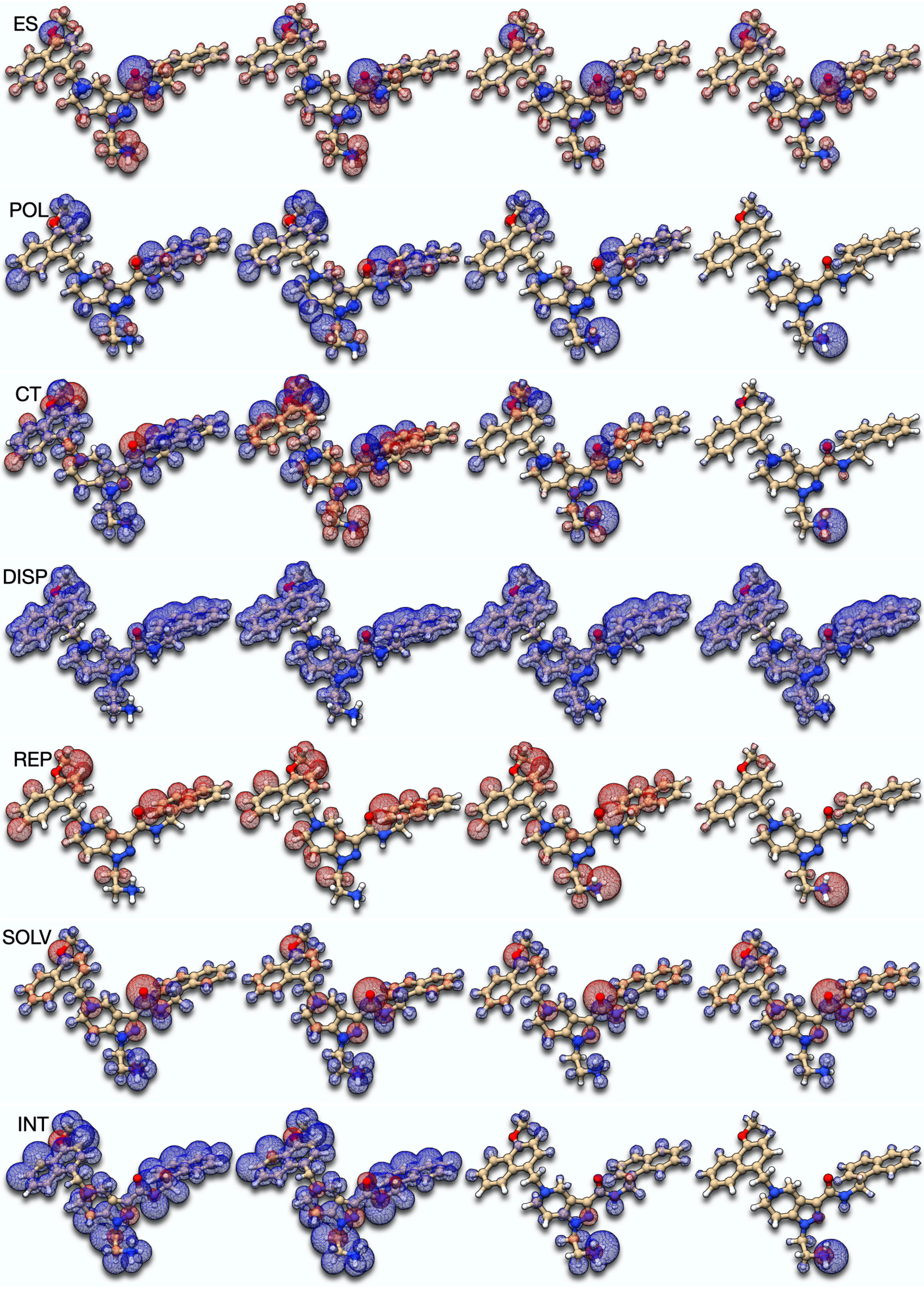
ES
POL
CT
DISP
REP
SOLV
INT

**Figure S14.2. Comparison of interaction maps for different poses of 5N8V. From left to right, the complexes 6.903, 6.857, 3.882 and 3.513 Å.**

**Table S14.1. Summary of different EDA contributions to the 4 poses of 5N8V. Energies in kcal/mol.**

| | poses | | | |
|---|---|---|---|---|
| | 1 | 2 | 3 | 4 |
| $R_{NC}$ (Å) | 6.903 | 6.857 | 3.882 | 3.513 |
| $E_{INT}$ | -23.155 | -23.435 | -31.007 | -34.700 |
| $E_{ES}$ | 58.041 | 60.942 | -4.542 | -19.251 |
| $E_{POL}$ | -33.839 | -33.819 | -45.923 | -65.394 |
| $E_{CT}$ | 0.001 | -0.014 | -0.388 | -2.640 |
| $E_{DISP}$ | -38.872 | -37.928 | -40.628 | -40.101 |
| $E_{SOLV}$ | -40.014 | -44.363 | 21.024 | 36.815 |
| $E_{REP}$ | 31.528 | 31.746 | 39.451 | 55.871 |

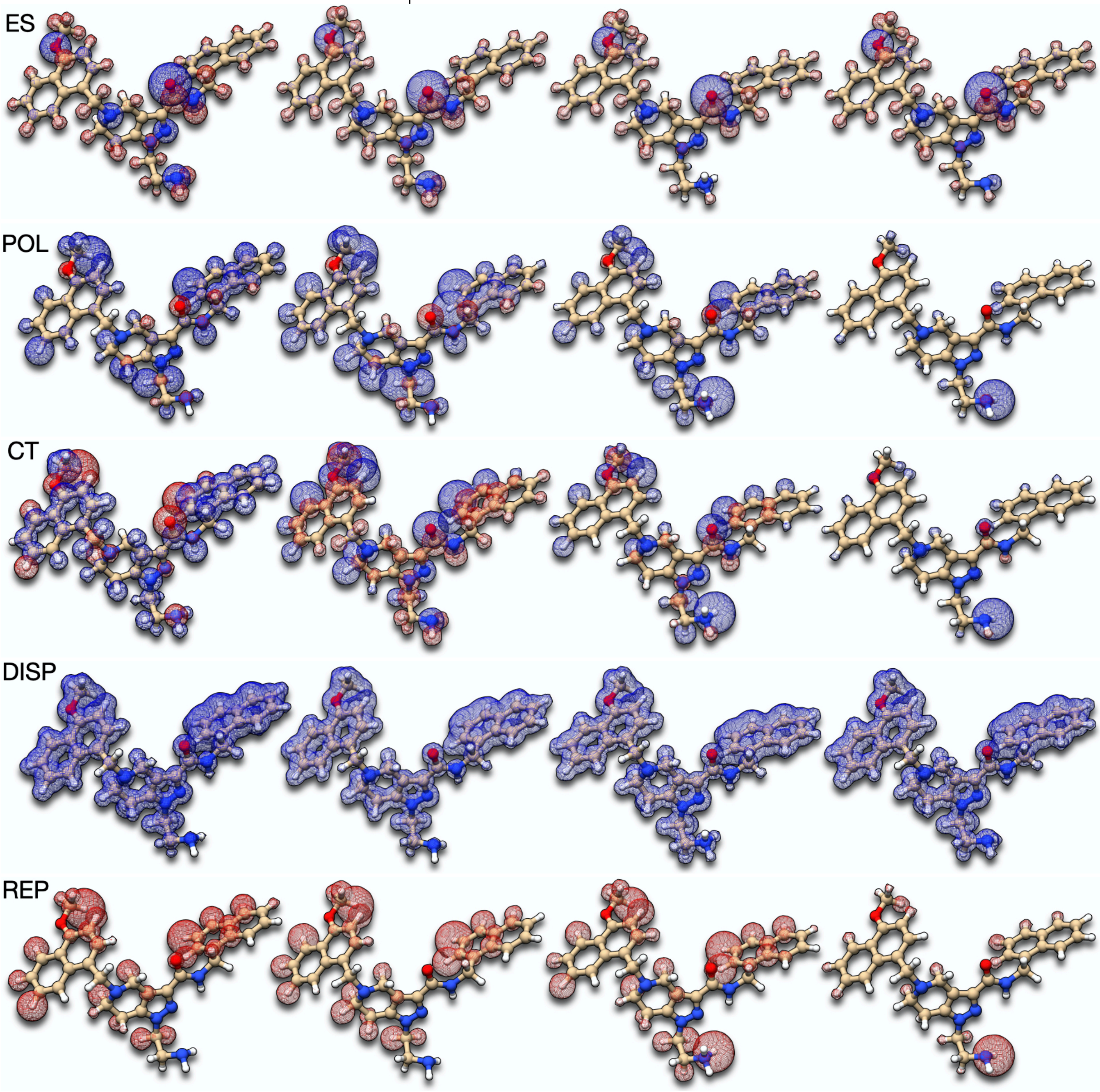

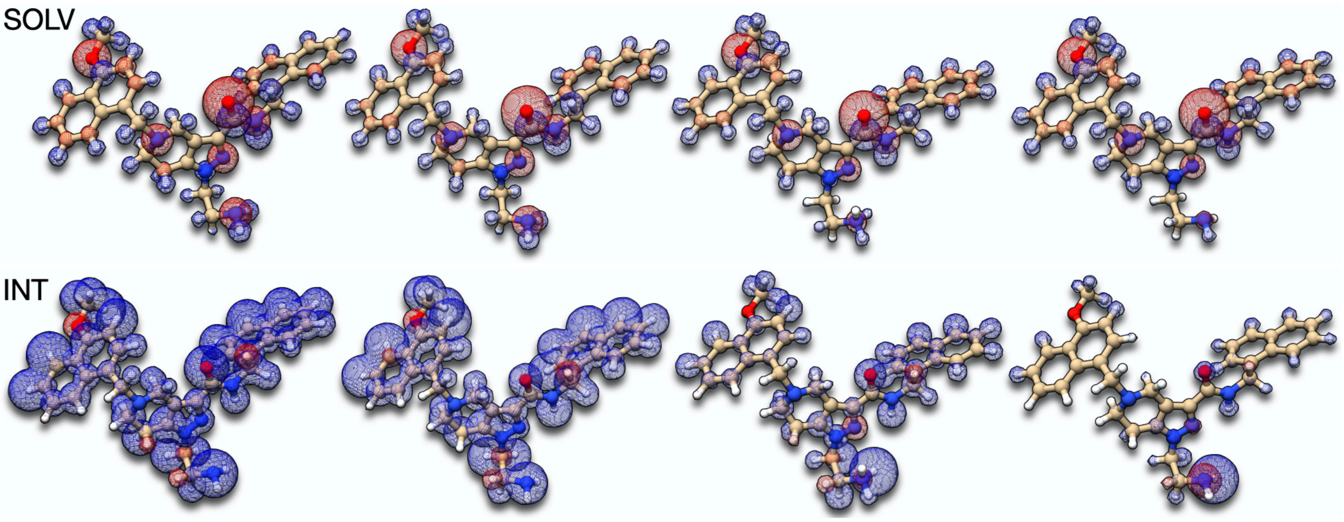


**Figure S14.3. Comparison of interaction maps for different poses of 5N8V using the neutral form of the ligand. From left to right, the complexes 6.903, 6.857, 3.882 and 3.513 Å.**

**Table S14.2. Summary of different EDA contributions to the 4 poses of 5N8V using the neutral form of the ligand. Energies in kcal/mol.**

| | poses | | | |
|---|---|---|---|---|
| | 1 | 2 | 3 | 4 |
| $R_{NC}$ (Å) | 6.903 | 6.857 | 3.882 | 3.513 |
| $E_{INT}$ | -27.199 | -26.699 | -31.756 | -30.798 |
| $E_{ES}$ | -19.867 | -20.390 | -31.295 | -31.838 |
| $E_{POL}$ | -26.800 | -27.145 | -31.073 | -48.167 |
| $E_{CT}$ | -0.007 | 0.007 | -0.114 | -0.881 |
| $E_{DISP}$ | -39.419 | -38.447 | -41.309 | -40.808 |
| $E_{SOLV}$ | 27.294 | 27.425 | 32.055 | 33.255 |
| $E_{REP}$ | 31.601 | 31.851 | 39.979 | 57.641 |

Crystal structures often contain several protein-ligand complexes in the asymmetric unit. One may therefore ask which unit to choose for further analysis. We selected, separately, the four protein-ligand complexes available in the PDB file 5N8V and ran the respective energy decomposition analyses. Each complex is identified uniquely by the distance between the ammonium's nitrogen in the ligand and the carboxylate's carbon in Glu16 ($R_{NC}$). We order the poses according to $R_{NC}$. Figure S14.4 contains the EDA results as a function of the $R_{NC}$ distance. The bar plots, table with the actual values, and the maps for the four poses of the ligand were presented above. Data for other complexes and a brief discussion is available in S15.

Electrostatics and solvation contributions are the most sensible to variations of $R_{NC}$. It is particularly interesting to note that somewhere between the two sets of structures available, both terms reverse their sign, such that at short contacts electrostatics become attractive, even though protein and ligand are both positively charged. This corresponds to the formation of a hydrogen bond with ionic bridge. Other contributions to the interaction of the two molecules show softer variations, though at very short distances, polarization and repulsion also exhibit marked differences, which attest the short contact.

The interaction maps for the four poses are quite instructive. When decreasing $R_{NC}$, electrostatics become more favorable. In particular, long-range repulsive electrostatics with PEX14 are overshaded by the hydrogen bond and ionic bridge with Glu16. To support this statement, the rest of ES maps remain largely unaffected in the four poses and similar considerations apply to all other maps. The exception is of course the total interaction terms, which shows the clear dominance of the above-mentioned hydrogen bond and ionic bridge. Charge transfer is increased when $R_{NC}$ is shortest, and the polarization and electronic density repulsion maps also reflect the increased strength of the hydrogen bond. This means that, matching our intuition and expectations, the hydrogen bond between ammonium and carboxylate only contributes for short enough $R_{NC}$ (below 6 Å). For large $R_{NC}$, only the attractive ionic contribution between groups remains. A closer look at the crystal structure of 5N8V reveals that the complexes with larger $R_{NC}$ have Glu16 involved in ionic bridges with the neighboring protein-ligand complex from the asymmetric unit, which renders the group unavailable to interact with the inhibitor. This is of course not directly reflected in the EDA calculations, because we only take one complex from the crystal for calculations. However, we note that the CT terms are negligible in the complexes with large $R_{NC}$. This is not so, when carboxyl and ammonium are close together. Since, our calculations reflect exclusively the protein-ligand interactions (and the internal statics of the protein), we hypothesize that charge transfer could be a good indicator of the extent the structure is affected by crystal artefacts.

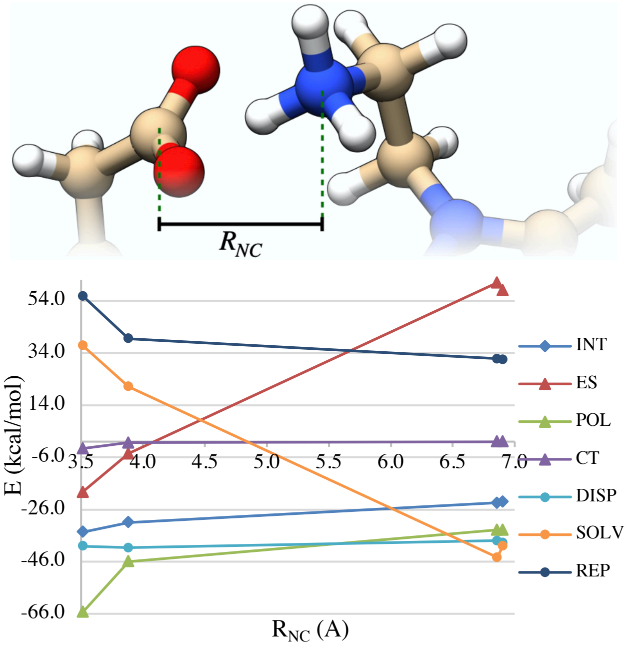


**Figure S14.4. Variation of the decomposed energies for the different poses of 5N8V available from different complexes of an assymetric unit of the crystal structure. $R_{NC}$ is the distance between 5N8V's ammonium (in position R2) and the protein's Glu16.**

These examples show furthermore how dangerous it is to select the improper protein-ligand complex for static studies, as incorrect conclusions might easily result from crystal artifacts. According to the results in Figure 2 of the main manuscript, the difference in binding between 5N8V and 5L87 is 2.8 kcal/mol, which compares well with the experimental value of 2.4 kcal/mol. However, as shown in this section, our analysis is strongly biased by crystal artefacts, and the difference in binding unreliable. This is in part because the nature of interactions between ligands and protein are intrinsically different: in place of an ethyl ammonium, 5L87 has a methyl group. Consequently, the contribution from binding entropies is not negligible.[19] The EDA results point however that crystal artefacts are at play.

Another question of relevance in computational studies is the protonation state of the ligand. Though amines are expected to be cationic due to their $pK_b$, the protonation state is not always unambiguously assigned. For that purpose, we re-ran the EDA calculations for all binding poses of 5N8V assuming a neutral form of the inhibitor. Results are gathered in Figure S14.5.

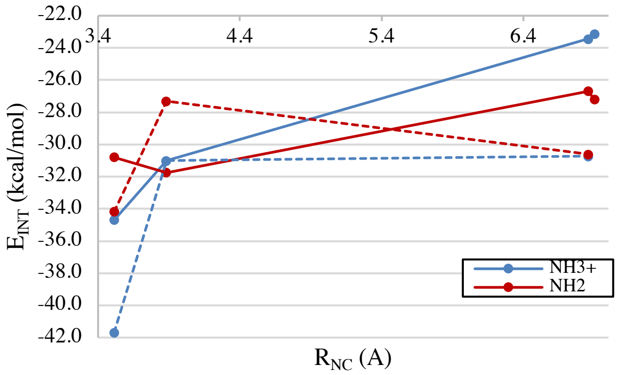


**Figure S14.5. Binding energy for the different poses of 5N8V at different protonation states of the ligand. $R_{NC}$ is the same as in figure 7. Full lines correspond to calculations with implicit solvation only, dashed lines consider additionally explicit waters.**

We start with the results using implicit water only (full lines), where it is easily observed that only at very short $R_{NC}$ the ammonium is favored over the neutral amine. The driving forces for the change in the nature of the interaction can be followed from the EDA contributions: the ammonium group shows less electronic density repulsion and a gain in charge transfer (formation of hydrogen bond with ionic bridge). The amine on the other hand shows systematically more favorable electrostatics and, only at the very shortest contact, also a smaller solvation penalty. It is interesting and simultaneously counterintuitive to observe that the amine form of the ligand offers better electrostatics than the respective ammonium, when probing the interactions of those groups with a glutamate. We stress however that these numbers are a global characterization of the complex and completely ignore local effects (like most other EDA algorithms). The introduction of the atom-specific partitioning and of the interaction maps aims precisely at overcoming such limitations and offering a clear picture of the locality of some interactions, which is actually useful for drug discovery.

The difference in the summed ES and SOLV terms decreases strongly with the distance $R_{NC}$. The lipophilic stabilization is quite conserved over all structures, which matches the expected behavior. However, when the ligand's ammonium in R2 dissociates from the carboxylate of Glu16, the ammonium is expected to be exposed to the aqueous phase. Even if water mediated hydrogen bonds are not formed with the protein, the $pK_b$ of primary amines indicates that the group is expected to be in the protonated form (ammonium). Re-running the calculations including explicit water (dashed lines) permits us to recover the expected behavior at short distances: the amine becomes less stable than the respective conjugate acid. At large $R_{NC}$ the ammonium is slightly more favorable than the amine. For practical purposes, however, we consider both protonation states of the amine to be isoenergetic. As discussed in the main manuscript (S16), this is in part a limitation of using an unrelaxed structure for the calculations. This is evident from the respective MD traces.

### S15 – Energy Decomposition Analysis for different poses of 5L8A, 5OML and 6RT2.

Since we study here several poses of different ligands inside PEX14, we need a measure of the relative position of the ligand in the pocket. In the main text and in the section above we define $R_{NC}$ as the distance between the carboxylate in Glu16 and a nitrogen of the ligand in an ammonium group. This is a suitable measure for 5N8V, since we are primarily interested in tracking the hydrogen bond between the two groups. In this section however, we study only ligands with no ammonium in R2. Therefore, we replace the ammonium's nitrogen by the pyrazole's nitrogen that is bound to R2.

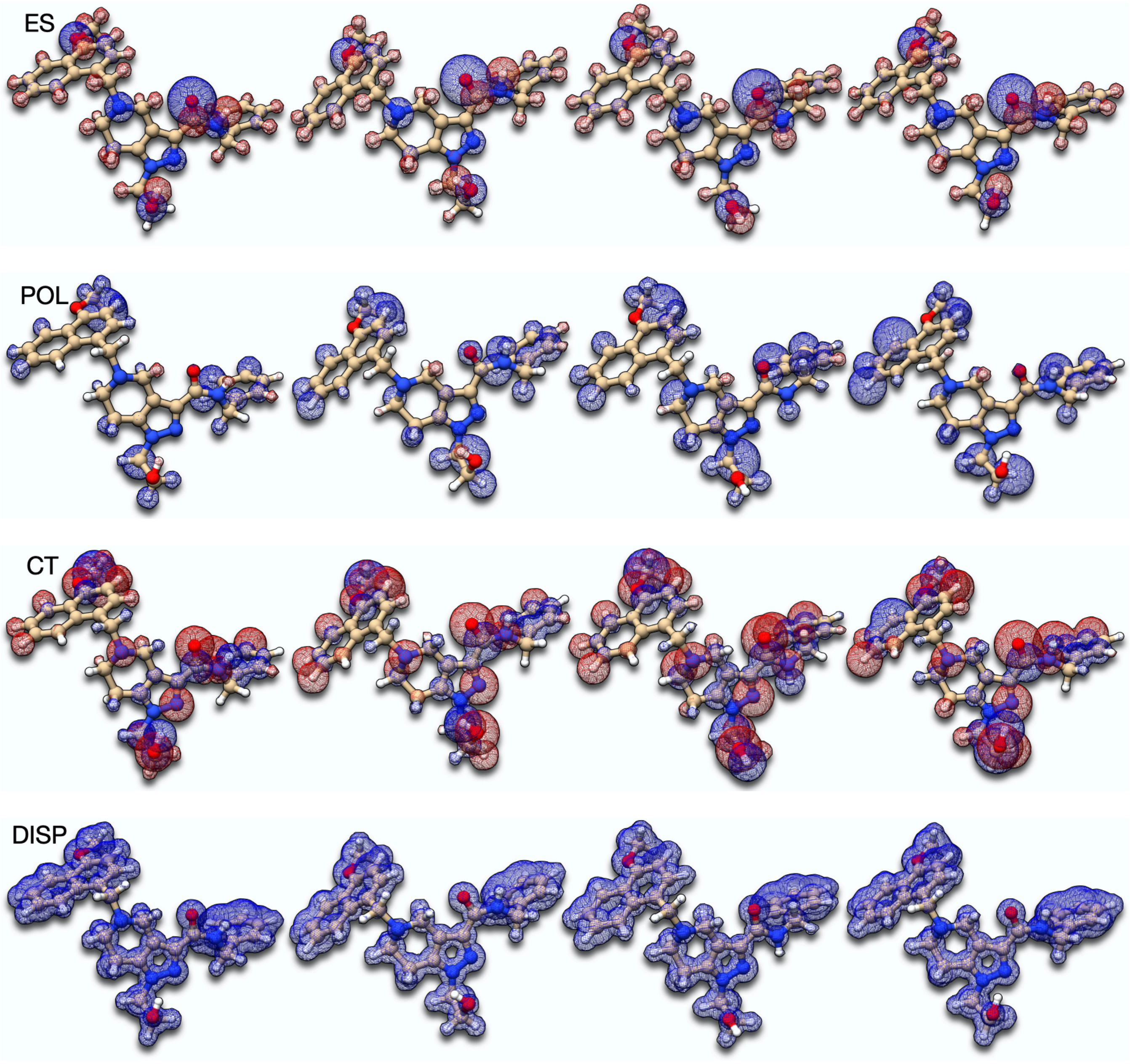

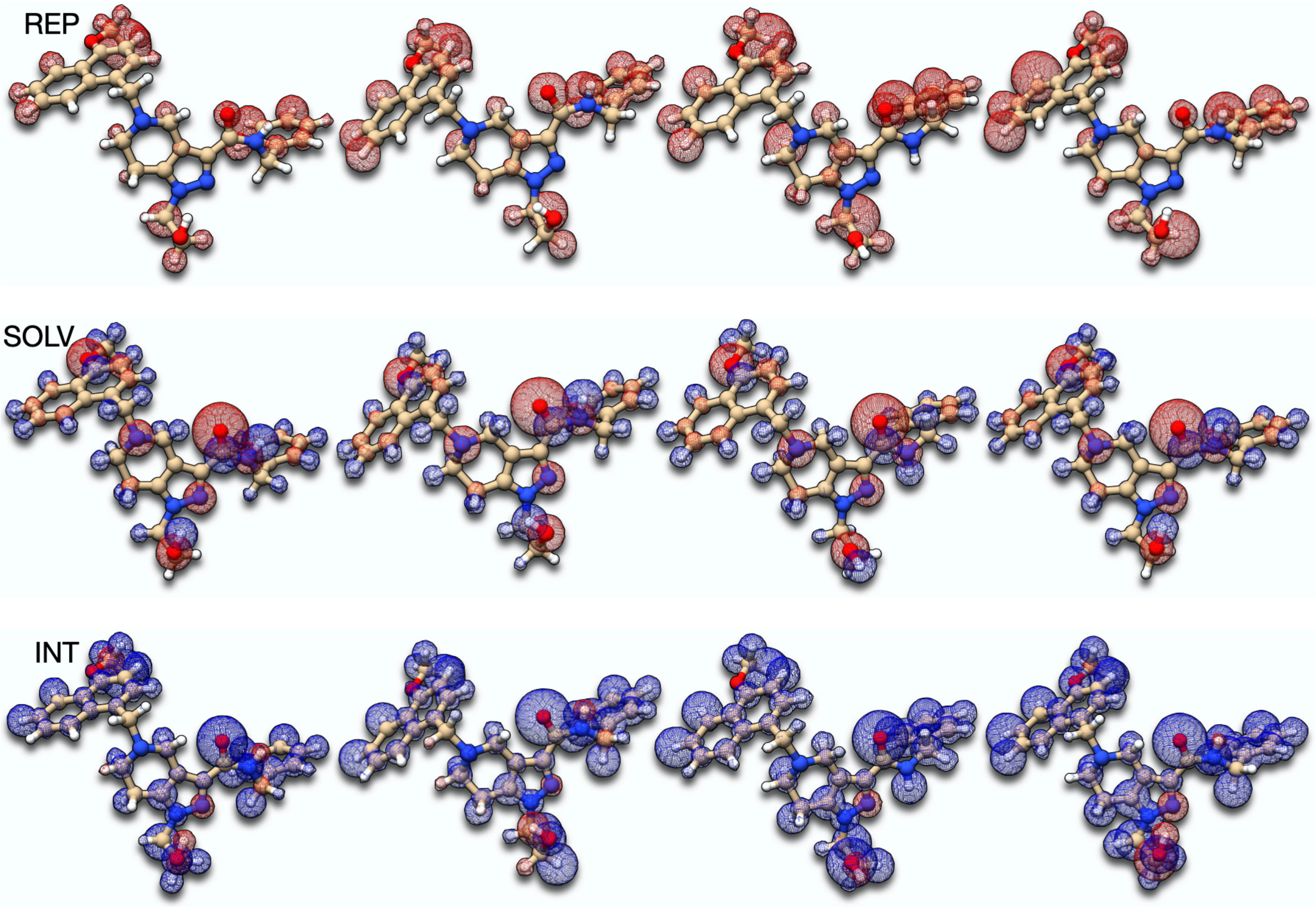


**Figure S15.1. Comparison of interaction maps for different poses of 5L8A. From left to right, the complexes 5.379, 5.204, 5.252 and 5.263 Å.**

**Table S15.1. Summary of different EDA contributions to the 4 poses of 5L8A. Energies in kcal/mol.**

| | poses | | | |
|---|---|---|---|---|
| | main | A | B | C |
| $R_{NC}$ (Å) | 5.379 | 5.204 | 5.252 | 5.263 |
| $E_{ES}$ | -16.479 | -9.043 | -24.023 | -10.560 |
| $E_{POL}$ | -35.315 | -25.426 | -31.799 | -40.988 |
| $E_{CT}$ | -0.078 | -0.193 | -0.230 | -0.295 |
| $E_{REP}$ | 47.377 | 35.518 | 42.667 | 57.248 |
| $E_{DISP}$ | -36.415 | -35.419 | -37.637 | -38.755 |
| $E_{SOLV}$ | 17.657 | 11.747 | 24.172 | 11.535 |
| $E_{INT}$ | -23.253 | -22.816 | -26.852 | -21.814 |

The maps for 5L8A are all very similar and there are only minor discrepancies in the nature of interactions involving some protons. On the other hand, the binding poses are also quite identical. This is primarily seen in the distance-measure used to track the position of the ligand inside the protein.

Similar considerations apply to 5OML. However, the maps for the fourth pose (C in Table S16.2) are different from the rest. This is the case where the ligand's carboxylate and Lys 38 interact by direct hydrogen bond. Differences are particularly noticeable in the polarization, charge transfer and repulsion maps. Comparing the repulsion maps for the different binding poses of 5OML one sees that the stereo hindrance between the methoxy-naphthyl group and the protein is not minimized in the last binding pose. This indicates that to form the hydrogen-bond interaction the ligand does not have to shift its position inside the pocket (thus breaking other important interactions). The reason for the missing hydrogen-bond interactions between ligand's carboxylate and Lys38 are therefore unrelated to geometric constraints (wrong stereochemistry or insufficient length of the substituent in R2).

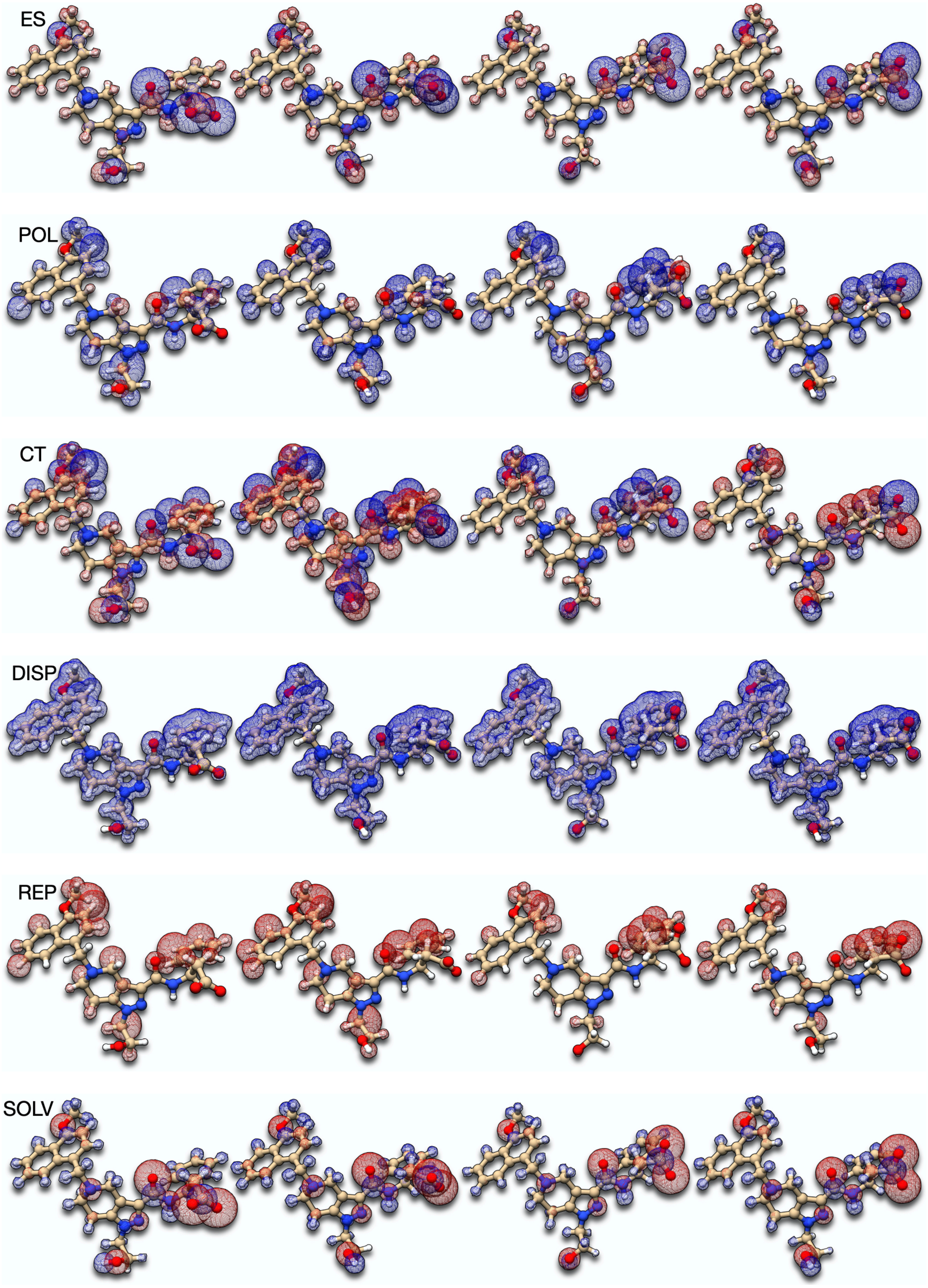
ES
POL
CT
DISP
REP
SOLV

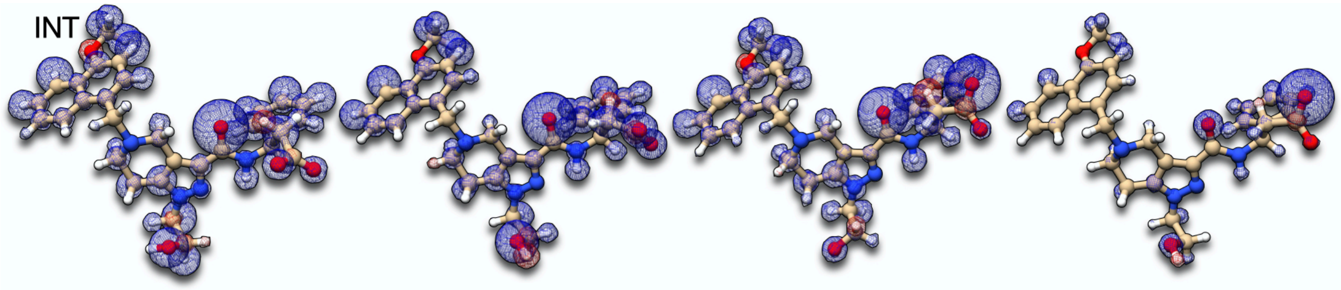


**Figure S15.2. Comparison of interaction maps for different poses of 5OML. From left to right, the complexes 6.320, 5.814, 7.808 and 7.709 Å.**

**Table S15.2. Summary of different EDA contributions to the 4 poses of 5OML. Energies in kcal/mol.**

| | poses | | | |
|---|---|---|---|---|
| | main | A | B | C |
| $R_{NC}$ (Å) | 6.320 | 5.814 | 7.808 | 7.709 |
| $E_{ES}$ | -147.507 | -126.974 | -173.530 | -156.709 |
| $E_{POL}$ | -20.721 | -14.583 | -29.402 | -26.646 |
| $E_{CT}$ | 0.040 | 0.035 | 0.062 | -0.760 |
| $E_{REP}$ | 30.420 | 30.080 | 34.934 | 35.352 |
| $E_{DISP}$ | -36.414 | -36.139 | -36.703 | -36.131 |
| $E_{SOLV}$ | 148.633 | 121.737 | 181.462 | 155.287 |
| $E_{INT}$ | -25.548 | -25.845 | -23.177 | -29.607 |

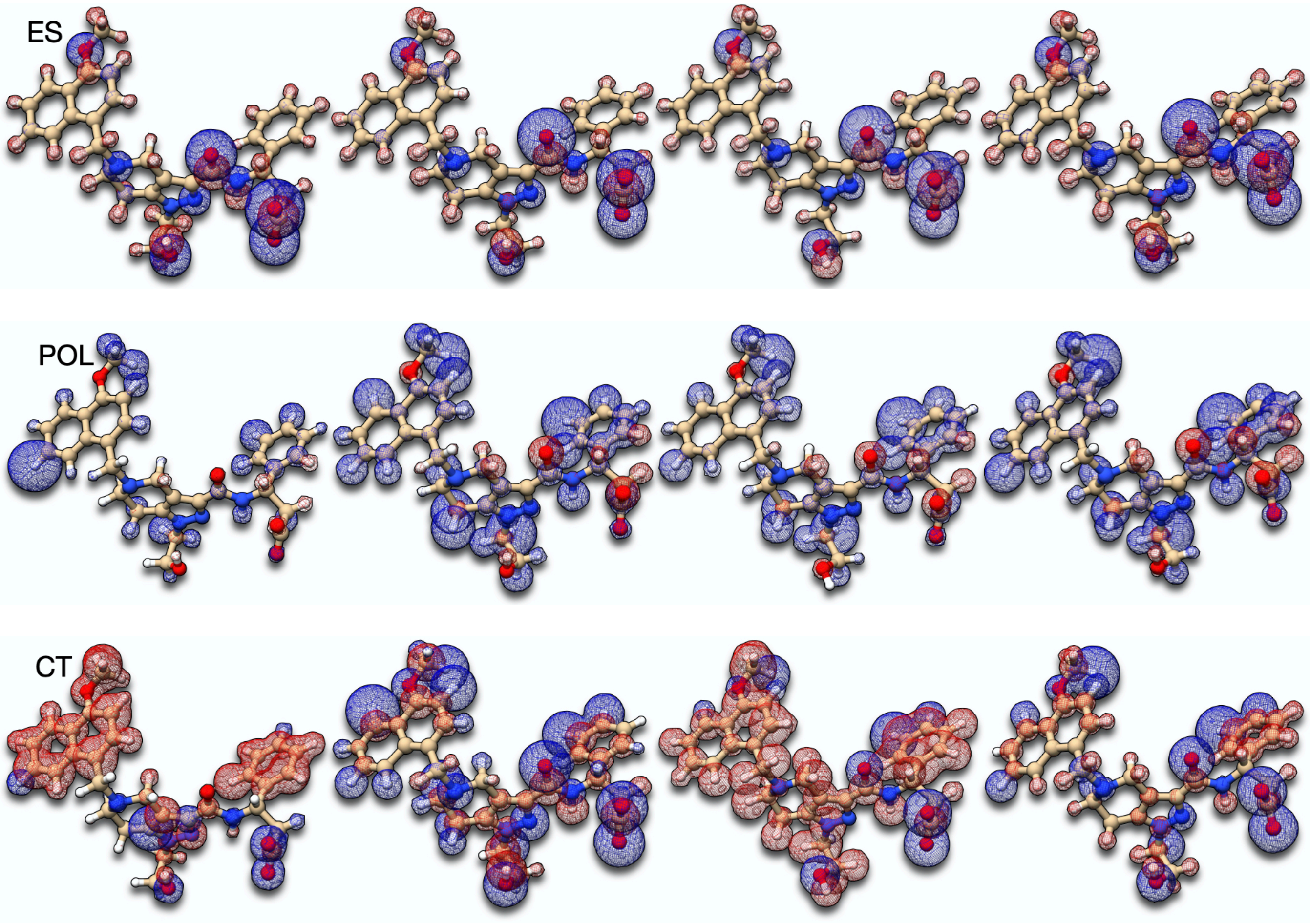

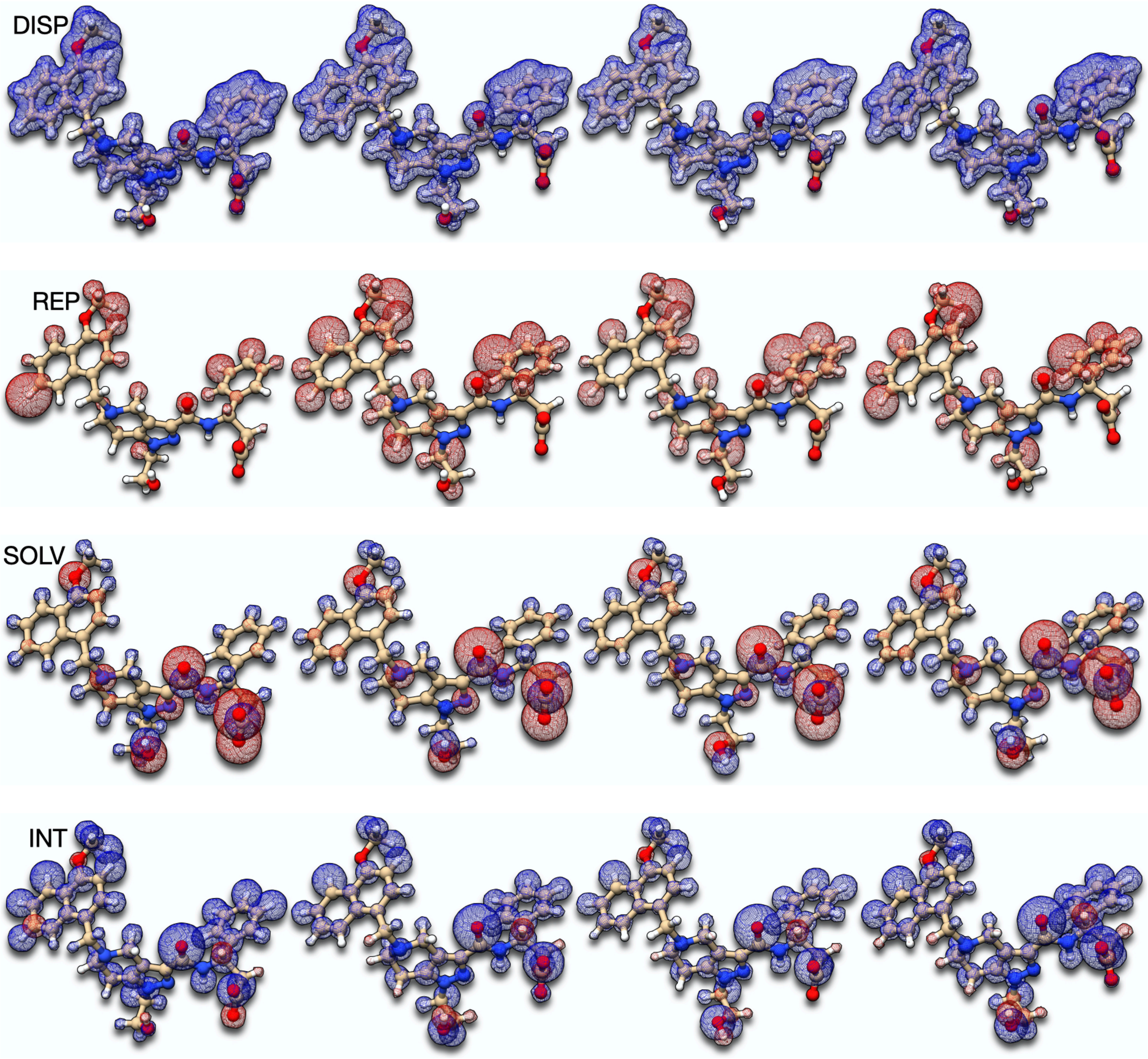


**Figure S15.3. Comparison of interaction maps for different poses of 6RT2. From left to right, the complexes 5.536, 6.339, 6.335 and 7.539 Å.**

**Table S15.3. Summary of different EDA contributions to the 4 poses of 6RT2. Energies in kcal/mol.**

| | poses | | | |
|---|---|---|---|---|
| | main | A | B | C |
| $R_{NC}$ (Å) | 5.536 | 6.339 | 6.335 | 7.539 |
| $E_{ES}$ | -168.030 | -129.953 | -137.042 | -123.858 |
| $E_{POL}$ | -29.447 | -19.124 | -19.466 | -14.150 |
| $E_{CT}$ | 0.004 | 0.037 | 0.008 | 0.012 |
| $E_{REP}$ | 37.020 | 34.228 | 32.512 | 28.671 |
| $E_{DISP}$ | -36.273 | -38.454 | -37.380 | -36.211 |
| $E_{SOLV}$ | 174.265 | 126.351 | 133.761 | 119.986 |
| $E_{INT}$ | -22.462 | -26.916 | -27.608 | -25.550 |

### S16 – Interactions Maps for Protein-Ligand Complexes with explicit waters and counter ions.

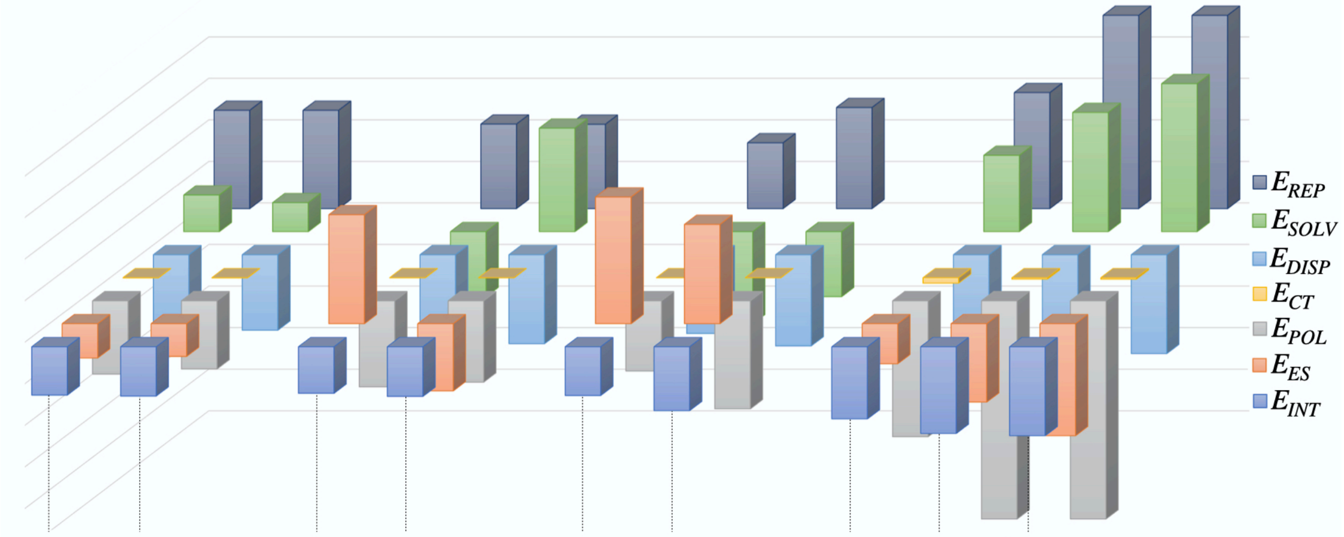


| 5L8A | | 6SPT | | 5N8V (6.857 Å) | | 5N8V (3.513 Å) | | | |
|---|---|---|---|---|---|---|---|---|---|
| (0,0) | (0,$Na^+$) | (0,0) | (0,$Cl^-$) | (0,0) | ($H_2O$,0) | (0,0) | ($H_2O$,0) | ($H_2O$,$Cl^-$) | |
| 47.38 | 47.38 | 40.73 | 40.64 | 31.75 | 48.70 | 55.87 | 93.14 | 93.13 | $E_{REP}$ |
| 17.66 | 13.97 | -31.27 | 49.86 | -44.36 | -31.27 | 36.81 | 57.42 | 71.26 | $E_{SOLV}$ |
| -36.42 | -36.36 | -42.80 | -42.84 | -37.93 | -43.91 | -40.10 | -47.59 | -47.61 | $E_{DISP}$ |
| -0.08 | -0.09 | -0.06 | -0.05 | -0.01 | -0.03 | -2.64 | -0.67 | -0.69 | $E_{CT}$ |
| -35.32 | -32.85 | -41.43 | -39.18 | -33.82 | -52.00 | -65.39 | -106.29 | -105.01 | $E_{POL}$ |
| -16.48 | -15.74 | 52.45 | -32.30 | 60.94 | 47.79 | -19.25 | -37.70 | -53.84 | $E_{ES}$ |
| -23.25 | -23.70 | -22.38 | -23.87 | -23.44 | -30.72 | -34.70 | -41.69 | -42.76 | $E_{INT}$ |

**Figure S16.1. Energy decomposition analyses for 3 protein-ligand complexes in different salt and explicit solvation conditions. Below the PDB code of each species, the code in parenthesis determines whether solvation waters were included ($H_2O$ on the left), whether ions were used (position right) or whether both elements were absent (0). For 5N8V, two poses with different $R_{NC}$ values were included. The latter are specified in parenthesis right to the PDB code. For the definition of $R_{NC}$, please refer to Figure 7 of the main manuscript. The embedded table gives the respective contributions in units of kcal/mol.**

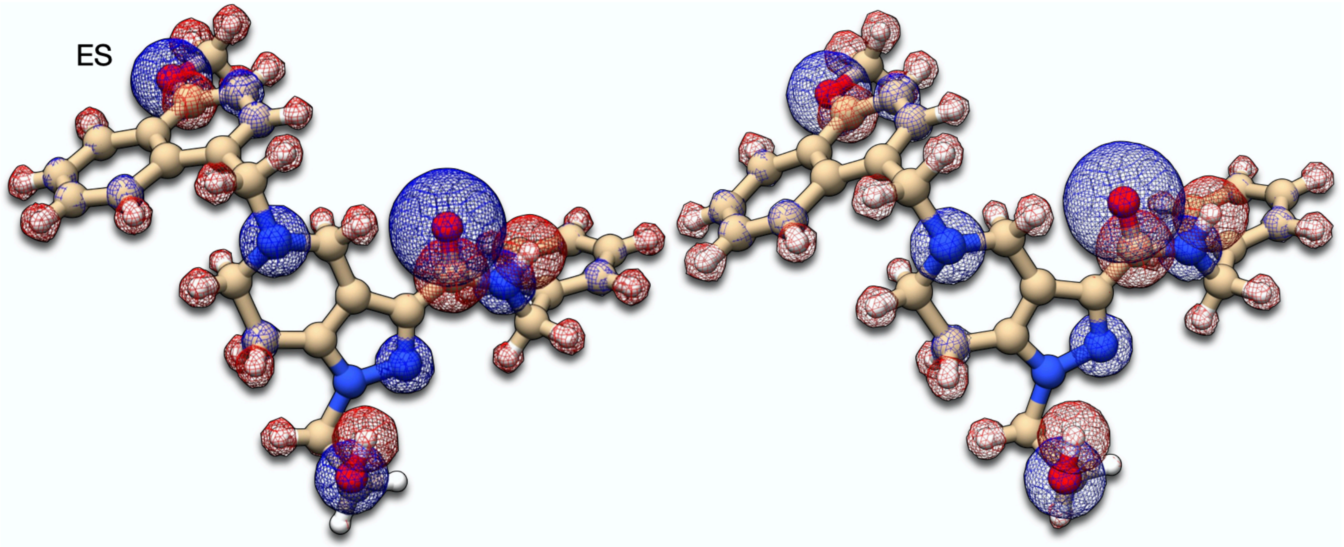

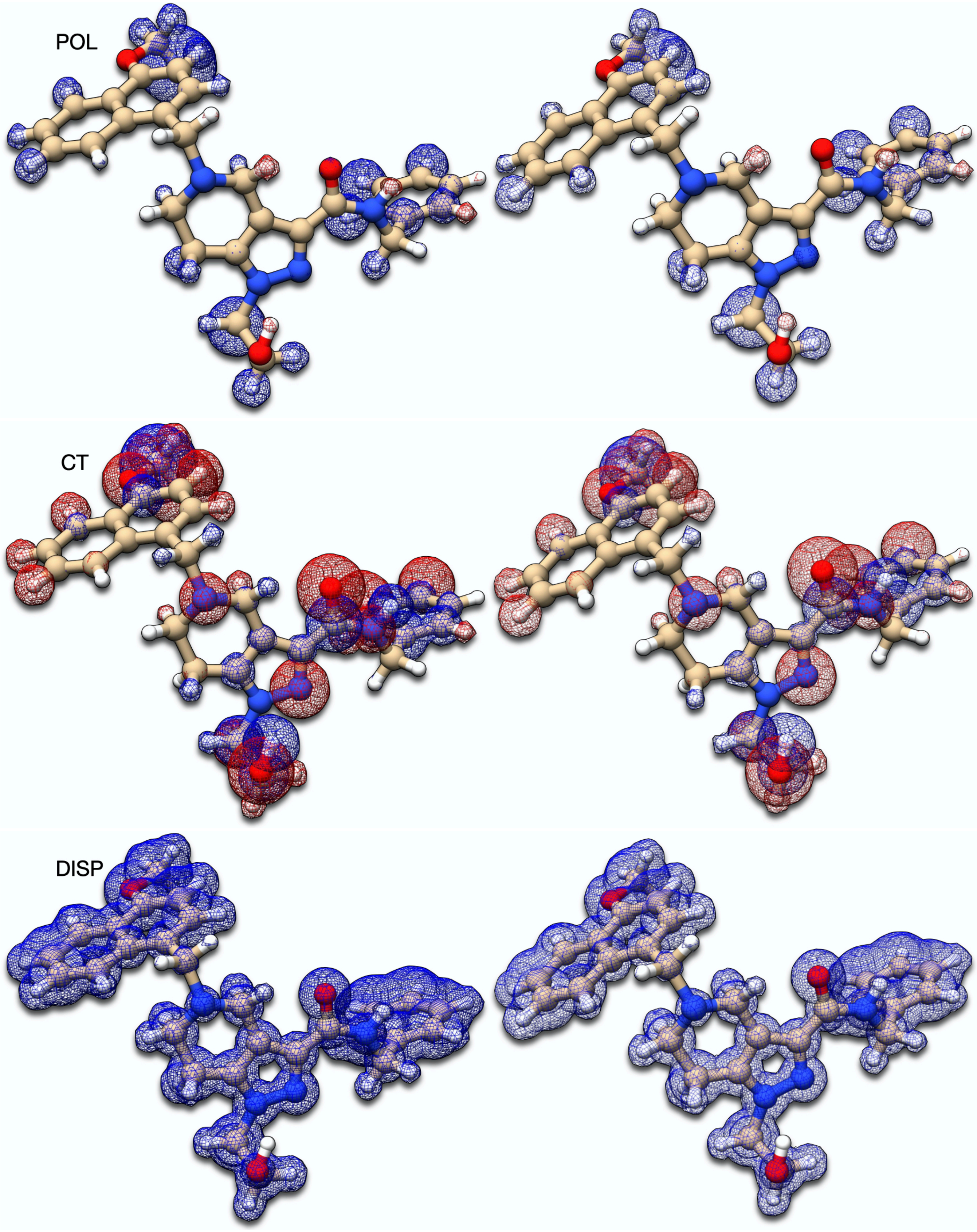
POL
CT
DISP

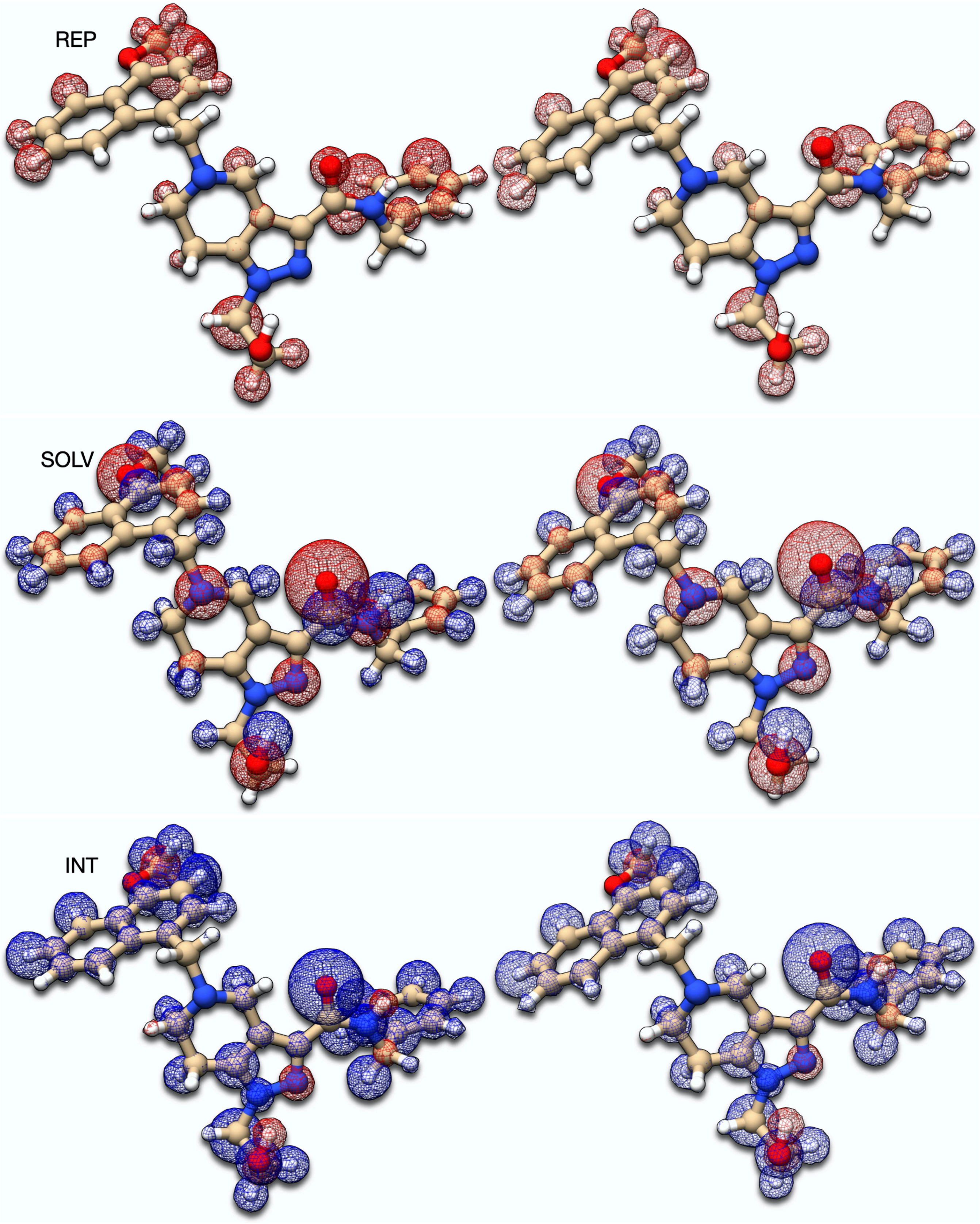


**Figure S16.2. Interaction maps for 5L8A without (left) and with (right) the sodium ion.**

The inhibitor in 5L8A has no group with formal charges. Most interactions remain therefore unaffected by the addition of a sodium cation that is a counter ion in the asymmetric unit. Exceptions are the solvation and polarization terms, with minor shifts that cancel each other out. The resulting binding energy is consequently of similar magnitude. The maps further reflect these observations, which is easily explained with the large distance between the sodium and the ligand: 8.95 Å to methoxy's oxygen, the closest polar group.

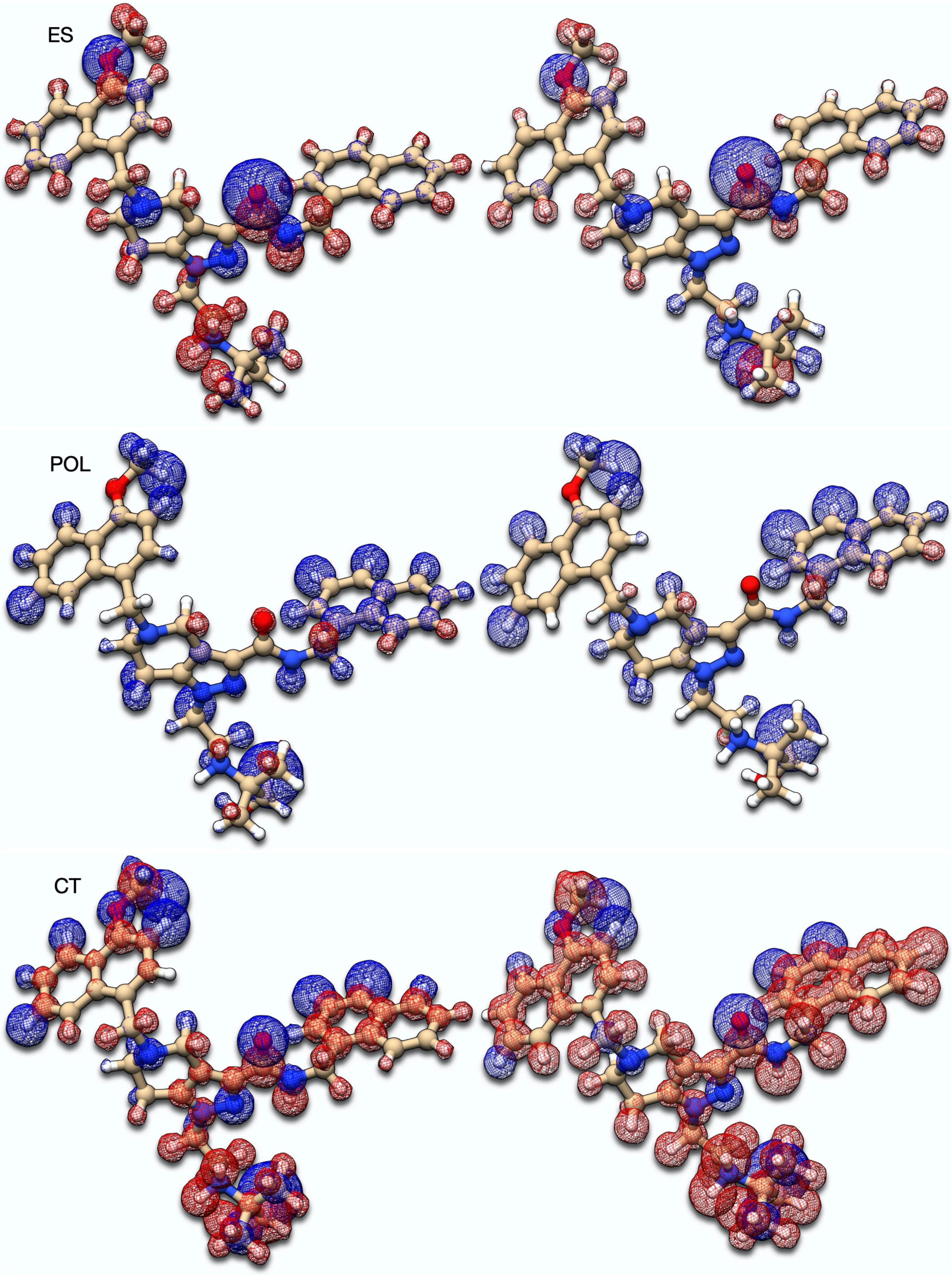
ES
POL
CT

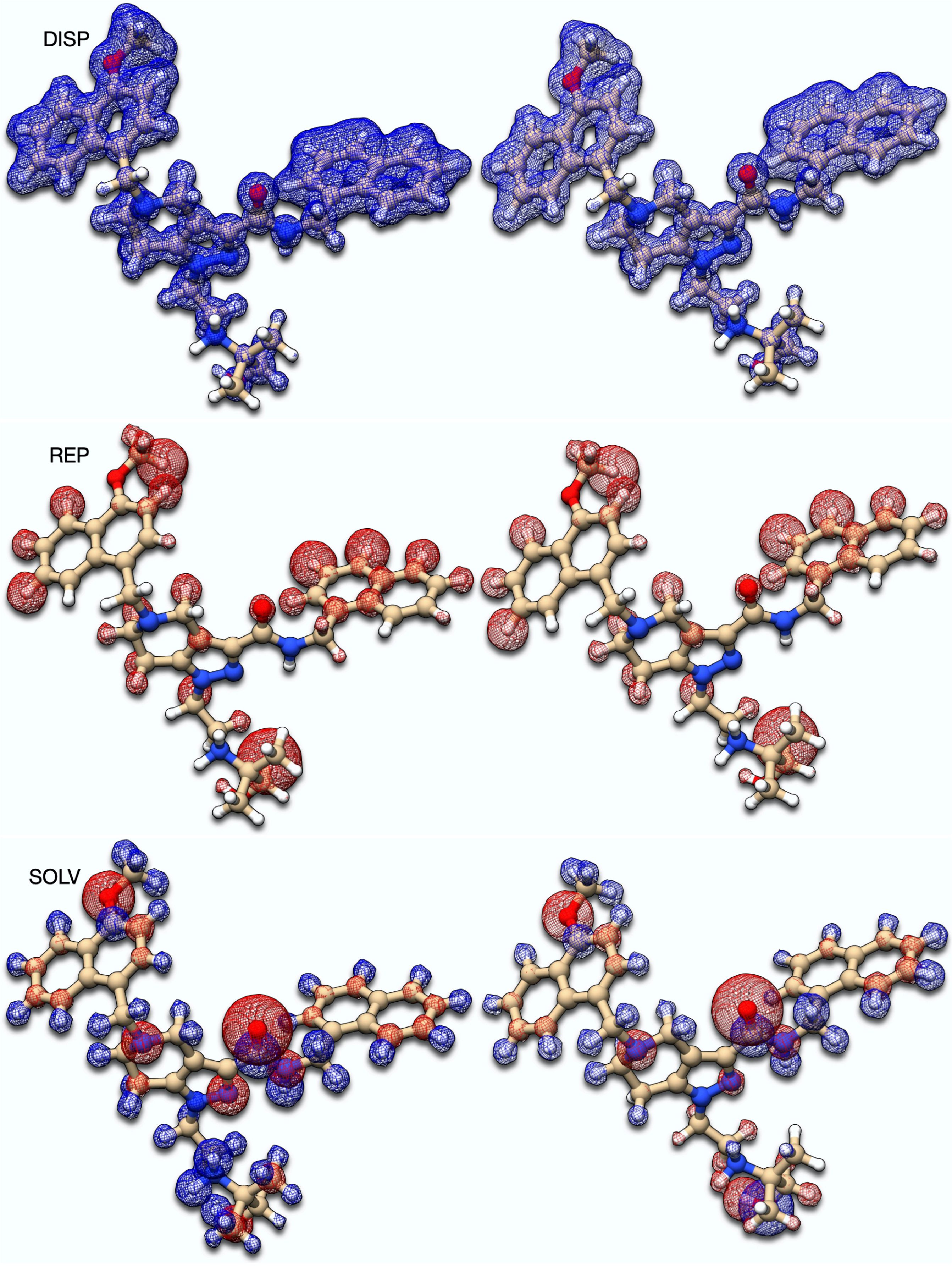
DISP
REP
SOLV

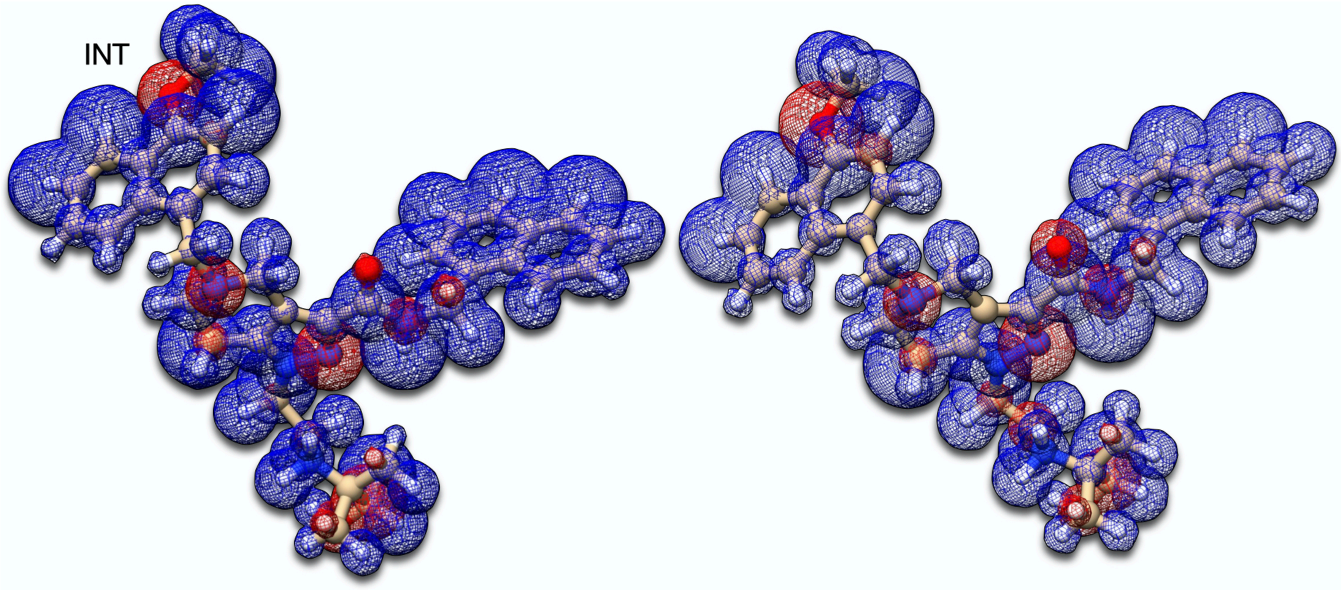


**Figure S16.3. Interaction maps for 6SPT without (left) and with (right) the chlorine ions.**

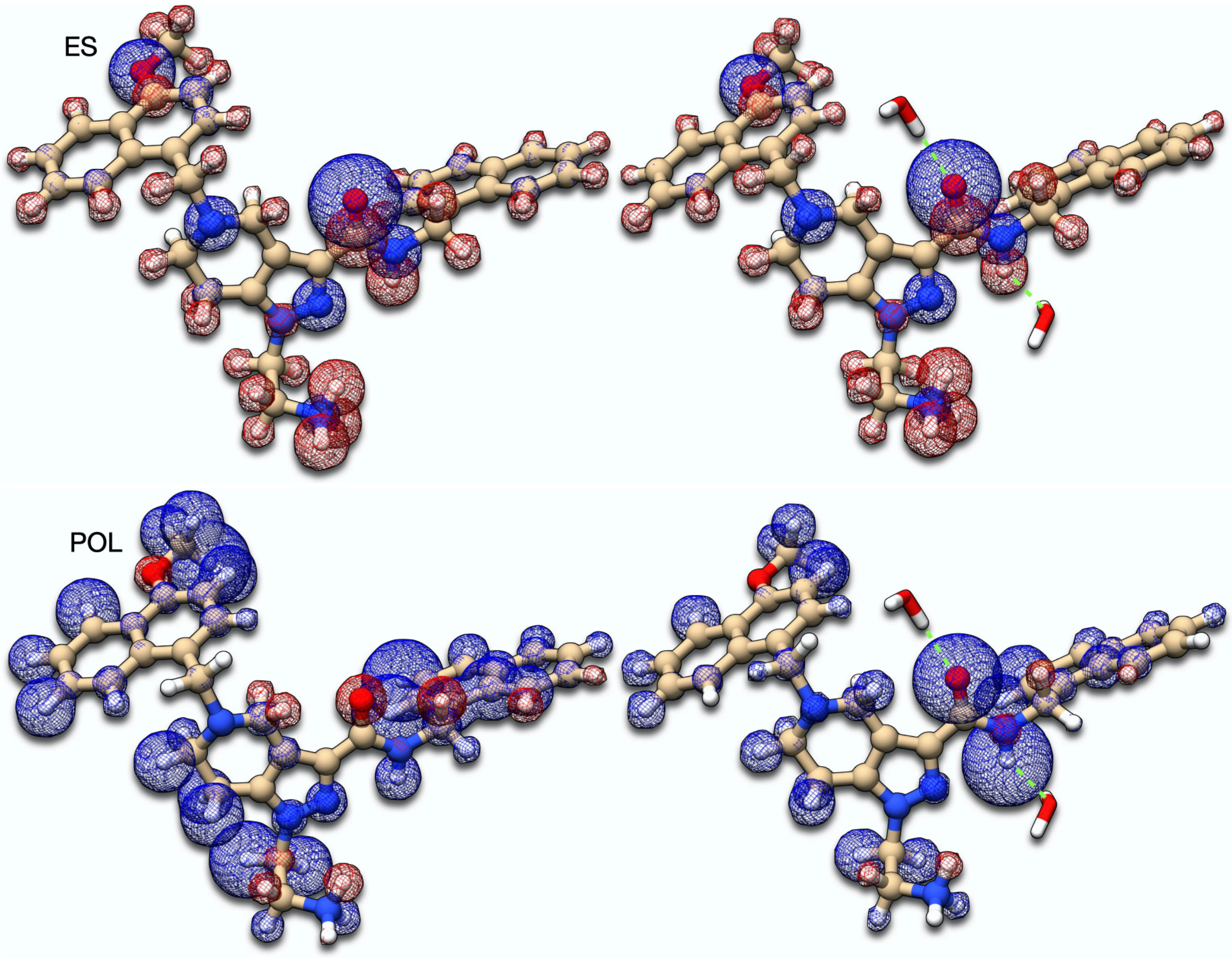

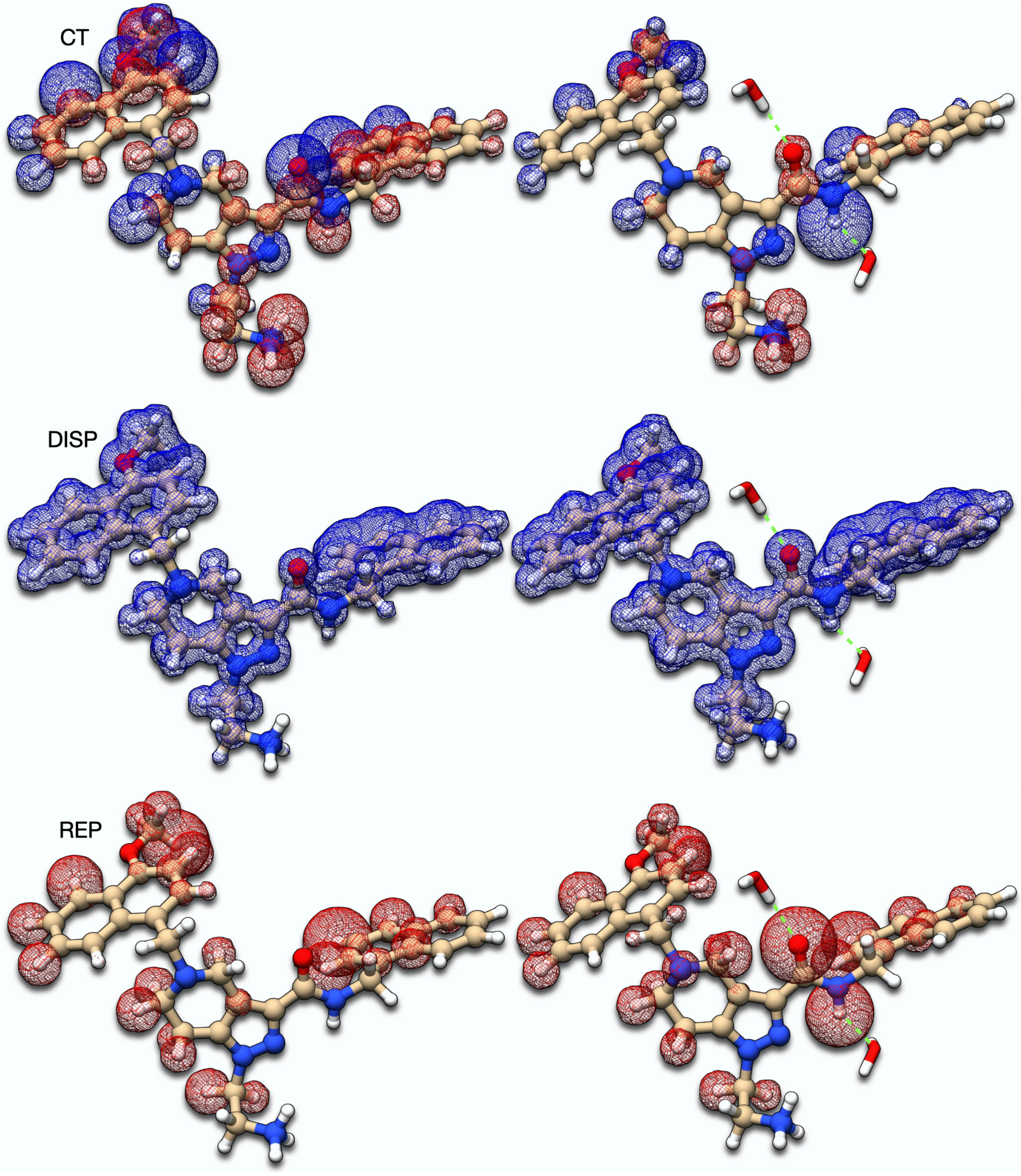
CT
DISP
REP

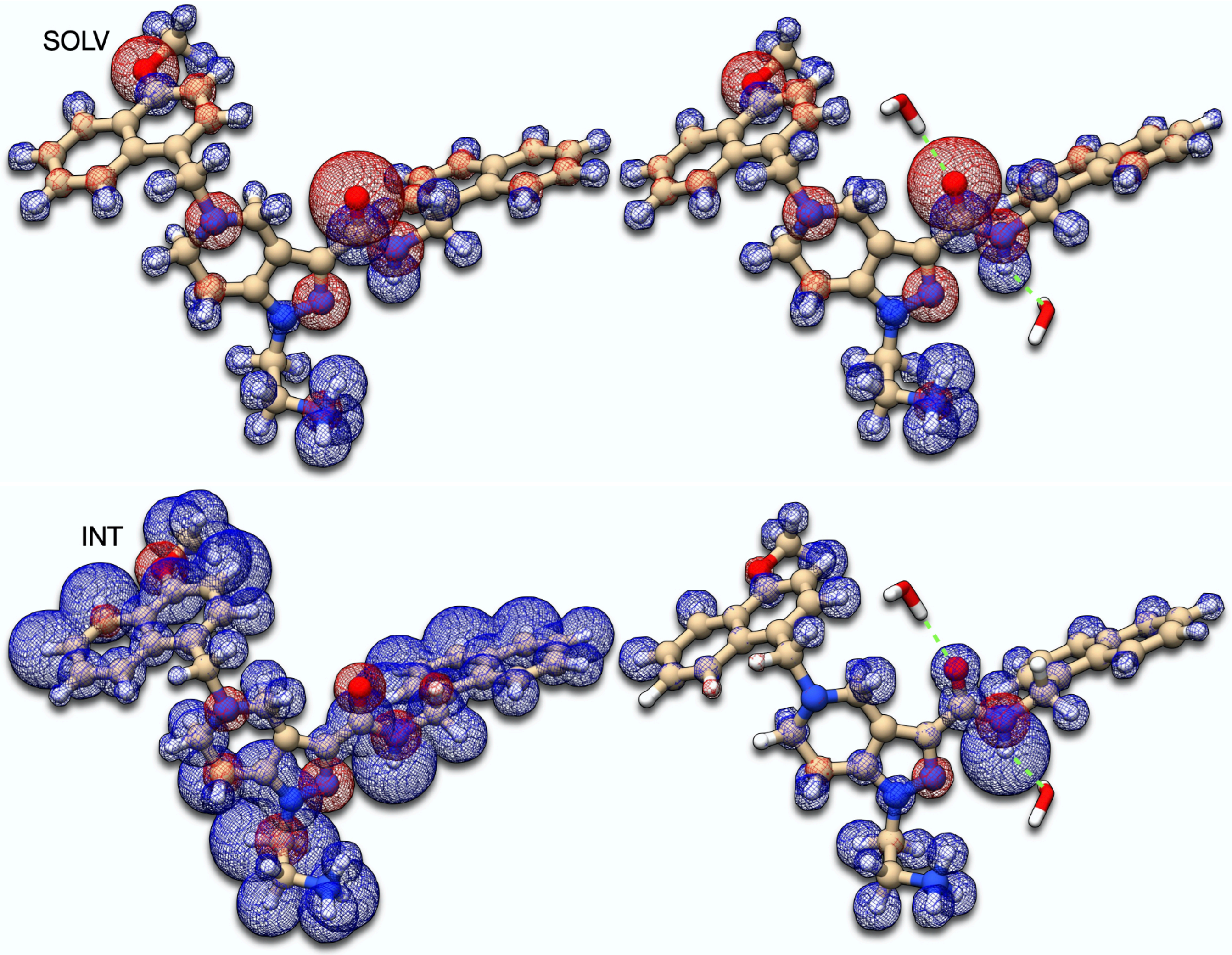


**Figure S16.4. Interaction maps for 5N8V (6.857) without (left) and with (right) explicit solvation waters. For clarity, only 2 out of 65 explicit waters are represented in the picture**

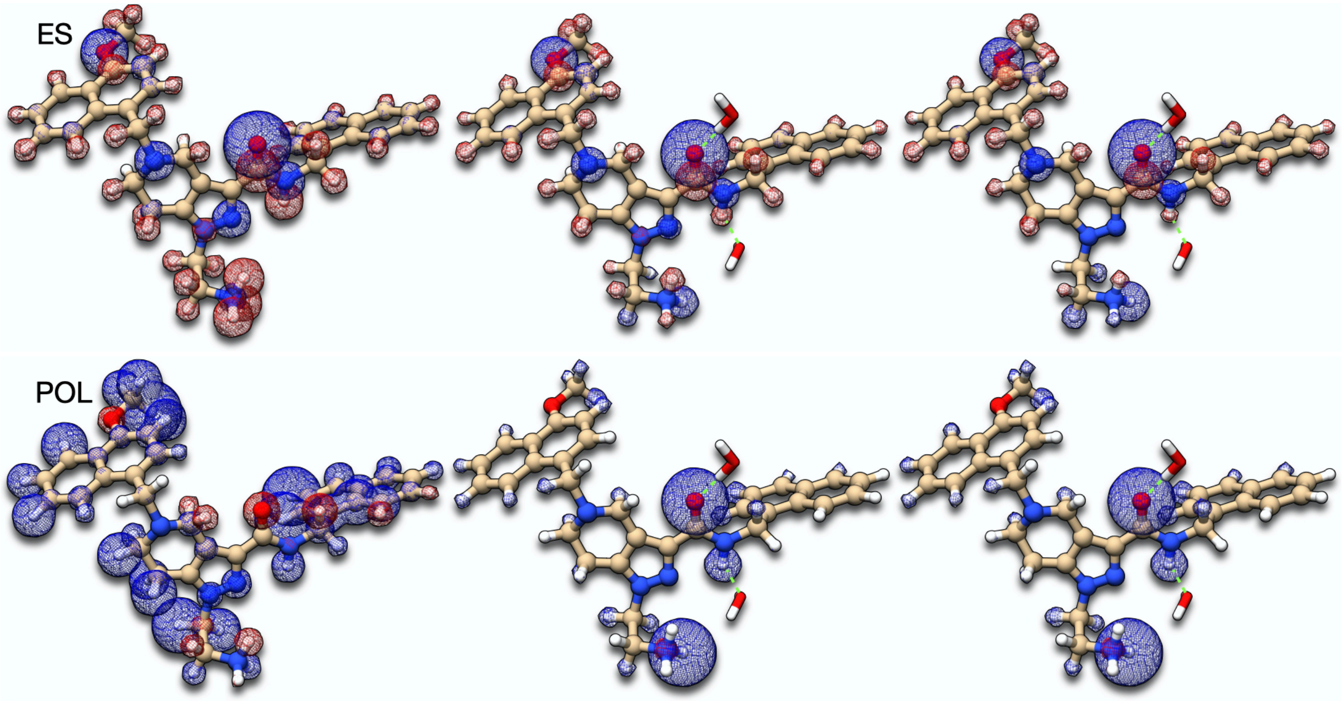

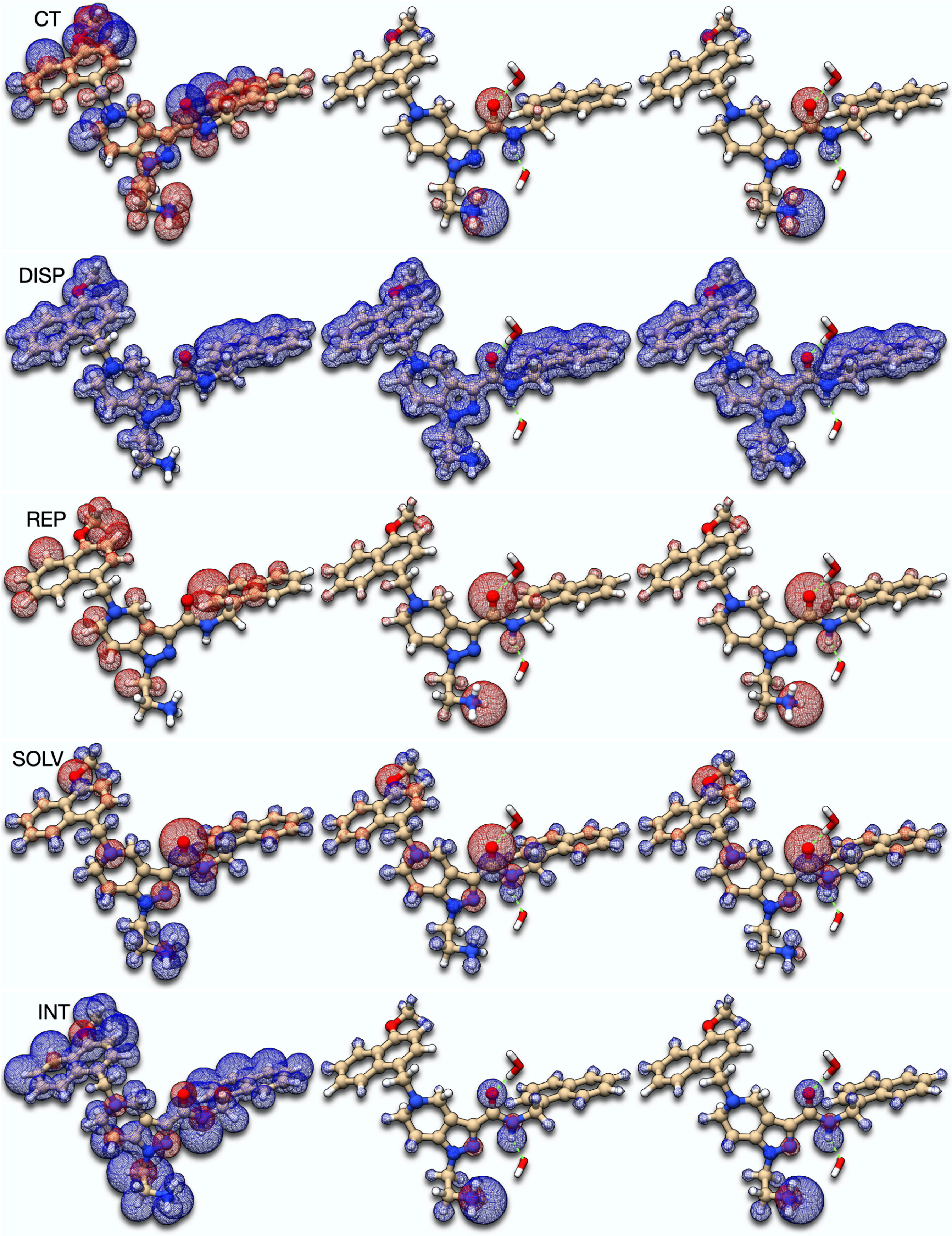


**Figure S16.5. Interaction maps for 5N8V (3.513) without water nor counter ions (left), with water but without counter ions (middle), and with water and with counter ions (right). For clarity, only 2 of the 59 explicit waters are represented in the picture.**

**Table S16.1. EDA data for 3 binding poses of 5N8V in two different protonation states of the ligand: left, positively charged with an ammonium group; right, the neutral amine.**

| | 5N8V (+) | | | 5N8V (0) | | |
|---|---|---|---|---|---|---|
| $R_{NC}$ | 3.51 | 3.88 | 6.86 | 3.51 | 3.88 | 6.86 |
| $E_{ES}$ | -37.70 | -26.82 | 47.79 | -43.82 | -45.36 | -27.19 |
| $E_{POL}$ | -106.29 | -89.11 | -52.00 | -89.09 | -75.91 | -45.72 |
| $E_{CT}$ | -0.67 | 0.05 | -0.03 | 0.03 | -0.03 | 0.02 |
| $E_{REP}$ | 93.14 | 83.45 | 48.70 | 95.18 | 84.72 | 48.87 |
| $E_{DISP}$ | -47.59 | -46.89 | -43.91 | -48.44 | -47.74 | -44.52 |
| $E_{SOLV}$ | 57.42 | 48.31 | -31.27 | 51.95 | 57.00 | 37.94 |
| $E_{INT}$ | -41.69 | -31.00 | -30.72 | -34.19 | -27.31 | -30.61 |

**Table S16.2. EDA data for 6SPT containing up to one chloride anion.**

| nCl | 0 | 1 | 1 | 1 | 1 |
|---|---|---|---|---|---|
| $E_{ES}$ | 52.45 | 28.33 | 22.37 | 39.13 | 35.12 |
| $E_{POL}$ | -41.43 | -41.29 | -42.14 | -40.54 | -40.31 |
| $E_{CT}$ | -0.06 | -0.06 | -0.06 | -0.06 | -0.06 |
| $E_{REP}$ | 40.73 | 40.68 | 40.73 | 40.71 | 40.71 |
| $E_{DISP}$ | -42.80 | -42.81 | -42.81 | -42.81 | -42.81 |
| $E_{SOLV}$ | -31.27 | -6.99 | 0.05 | -19.74 | -16.08 |
| $E_{INT}$ | -22.38 | -22.14 | -21.86 | -23.31 | -23.44 |

**Table S16.3. EDA data for 6SPT with 2 chloride anions.**

| nCl | 2 | 2 | 2 | 2 | 2 | 2 |
|---|---|---|---|---|---|---|
| $E_{ES}$ | -1.70 | 15.02 | 11.02 | 9.06 | 5.06 | 21.81 |
| $E_{POL}$ | -41.70 | -40.30 | -40.10 | -41.20 | -40.91 | -39.24 |
| $E_{CT}$ | -0.06 | -0.06 | -0.06 | -0.06 | -0.06 | -0.06 |
| $E_{REP}$ | 40.68 | 40.66 | 40.66 | 40.71 | 40.70 | 40.69 |
| $E_{DISP}$ | -42.82 | -42.82 | -42.82 | -42.82 | -42.82 | -42.82 |
| $E_{SOLV}$ | 23.91 | 4.39 | 8.07 | 11.51 | 15.05 | -4.79 |
| $E_{INT}$ | -21.69 | -23.11 | -23.22 | -22.81 | -22.97 | -24.42 |

**Table S16.4. EDA data for 6SPT with 3 or 4 chloride anions.**

| nCl | 3 | 3 | 3 | 3 | 4 |
|---|---|---|---|---|---|
| $E_{ES}$ | -15.01 | -18.99 | -2.29 | -8.25 | -32.30 |
| $E_{POL}$ | -40.67 | -40.39 | -38.92 | -39.79 | -39.18 |
| $E_{CT}$ | -0.06 | -0.05 | -0.05 | -0.06 | -0.05 |
| $E_{REP}$ | 40.66 | 40.66 | 40.64 | 40.68 | 40.64 |
| $E_{DISP}$ | -42.83 | -42.83 | -42.83 | -42.83 | -42.84 |
| $E_{SOLV}$ | 35.22 | 38.78 | 19.22 | 26.27 | 49.86 |
| $E_{INT}$ | -22.67 | -22.82 | -24.24 | -23.98 | -23.87 |

The ligand in 6SPT contains a positively charged group, and from the crystal structure, up to 4 chloride anions may be included in the calculations. The minimum distances between chlorides and the ligand are 11.65 Å, 14.05Å, 18.76 Å and 16.98 Å. In Figure S16.6 we show the effect of the different chlorides and their statistical combinations onto the binding of the inhibitor. Most energy contributions are minimally impacted when introducing anions, except for electrostatics and solvation: as the number of these counter ions increases, electrostatics become more favorable, and the solvation terms even become repulsive. The crosso-ver point (where the sign in ES and SOLV term reverses) seems to be when two chloride anions are included in the calculations. The maps are in this situation quite clear, as they show exclusively changes on the substituent R2 (Figure 8 A, B, C, and D). None-theless, the impact on binding energies is quite shallow, with a maximum difference of 1.5 kcal/mol. This difference may represent however a 10-fold variation in dissociation constants. Though inarguably important for binding, the chlorides may be classified as a long-range electrostatic effect.

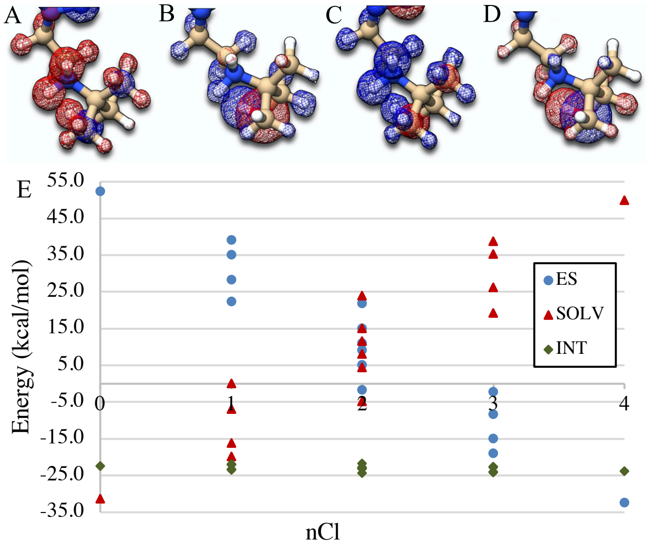


**Figure S16.6. The effect of the number of chloride anions on the electrostatics, solvation, and binding of 6SPT. A) the ES maps with 0 and B) with 4 $Cl^-$. C) The SOLV maps with 0 and D) with 4 $Cl^-$. E) The ES (blue circles), SOLV (red triangles), and INT (green diamonds) terms as a function of the number of $Cl^-$ included in the calculation.**

**Table S16.5. EDA data for 5N8V in its ammonium form as a function of time (in ps).**

| | 0 | 10 | 20 | 30 | 40 | 50 |
|---|---|---|---|---|---|---|
| $E_{ES}$ | -67.20 | -49.56 | -70.09 | -79.92 | -83.00 | -88.26 |
| $E_{POL}$ | -158.65 | -153.32 | -214.30 | -200.10 | -181.91 | -219.35 |
| $E_{CT}$ | -0.06 | -4.21 | -0.33 | -1.31 | -0.65 | -1.14 |
| $E_{REP}$ | 150.85 | 147.55 | 210.47 | 203.01 | 177.92 | 213.39 |
| $E_{DISP}$ | -57.54 | -56.82 | -59.22 | -59.94 | -57.24 | -57.25 |
| $E_{SOLV}$ | 57.28 | 54.52 | 68.35 | 76.86 | 77.39 | 97.70 |
| $E_{INT}$ | -75.32 | -61.83 | -65.12 | -61.39 | -67.48 | -54.90 |

**Table S16.6. EDA data for 5N8V in its amine form as a function of time (in ps).**

| | 0 | 10 | 20 | 30 | 40 | 50 |
|---|---|---|---|---|---|---|
| $E_{ES}$ | -46.15 | -51.16 | -48.18 | -43.98 | -45.69 | -41.92 |
| $E_{POL}$ | -154.56 | -204.49 | -127.17 | -135.00 | -140.46 | -171.95 |
| $E_{CT}$ | -3.79 | -0.36 | -4.12 | -5.85 | -6.87 | -1.62 |
| $E_{REP}$ | 165.46 | 216.60 | 136.02 | 144.81 | 154.81 | 184.33 |
| $E_{DISP}$ | -61.88 | -62.28 | -61.39 | -61.67 | -58.70 | -60.94 |
| $E_{SOLV}$ | 59.63 | 65.25 | 61.92 | 60.90 | 63.82 | 56.02 |
| $E_{INT}$ | -41.28 | -36.45 | -42.92 | -40.78 | -33.10 | -36.09 |

Below the data related to EDA on the dynamical simulations of the ammonium and amine variants of 5N8V. Information on the dynamical simulations given in S19.

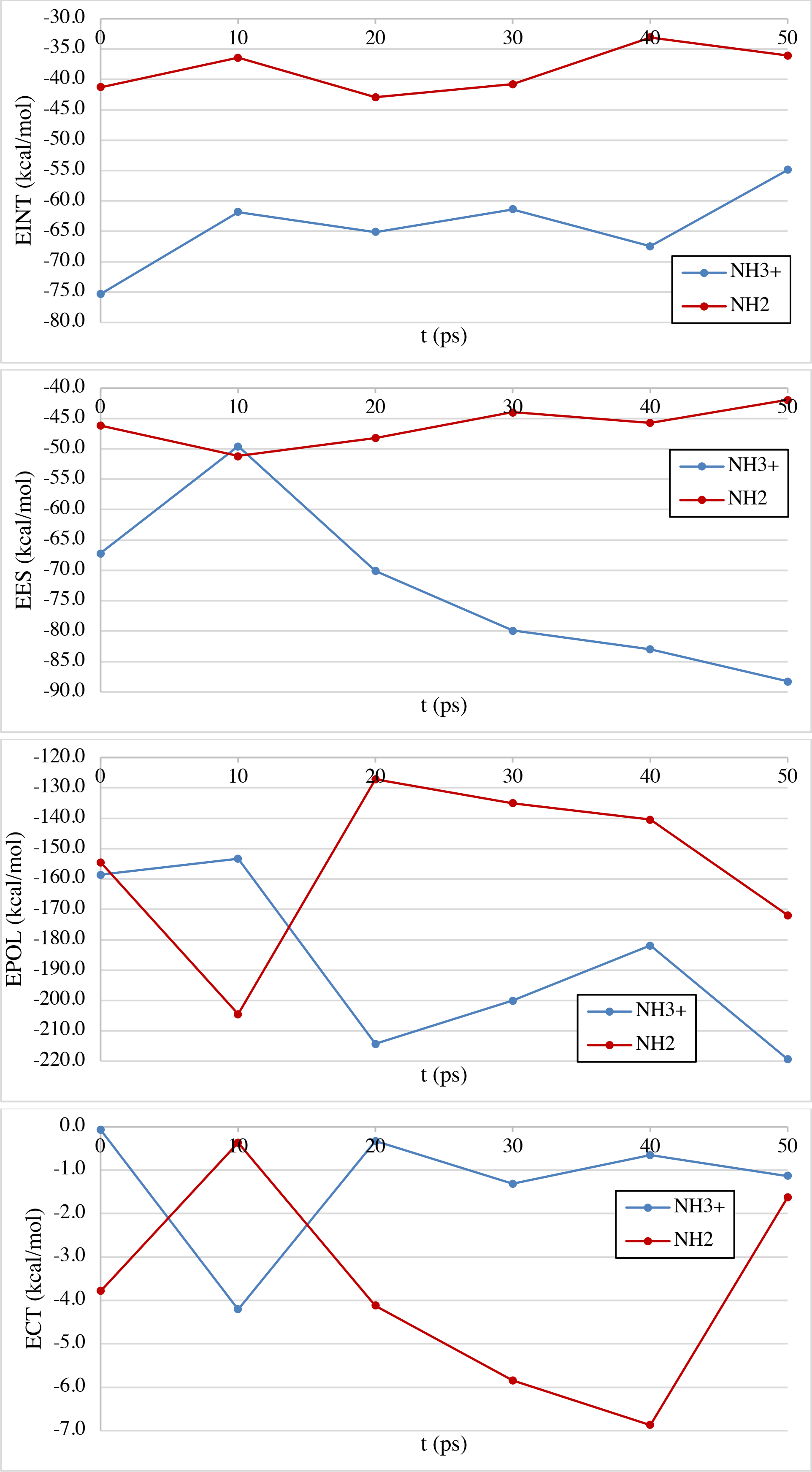

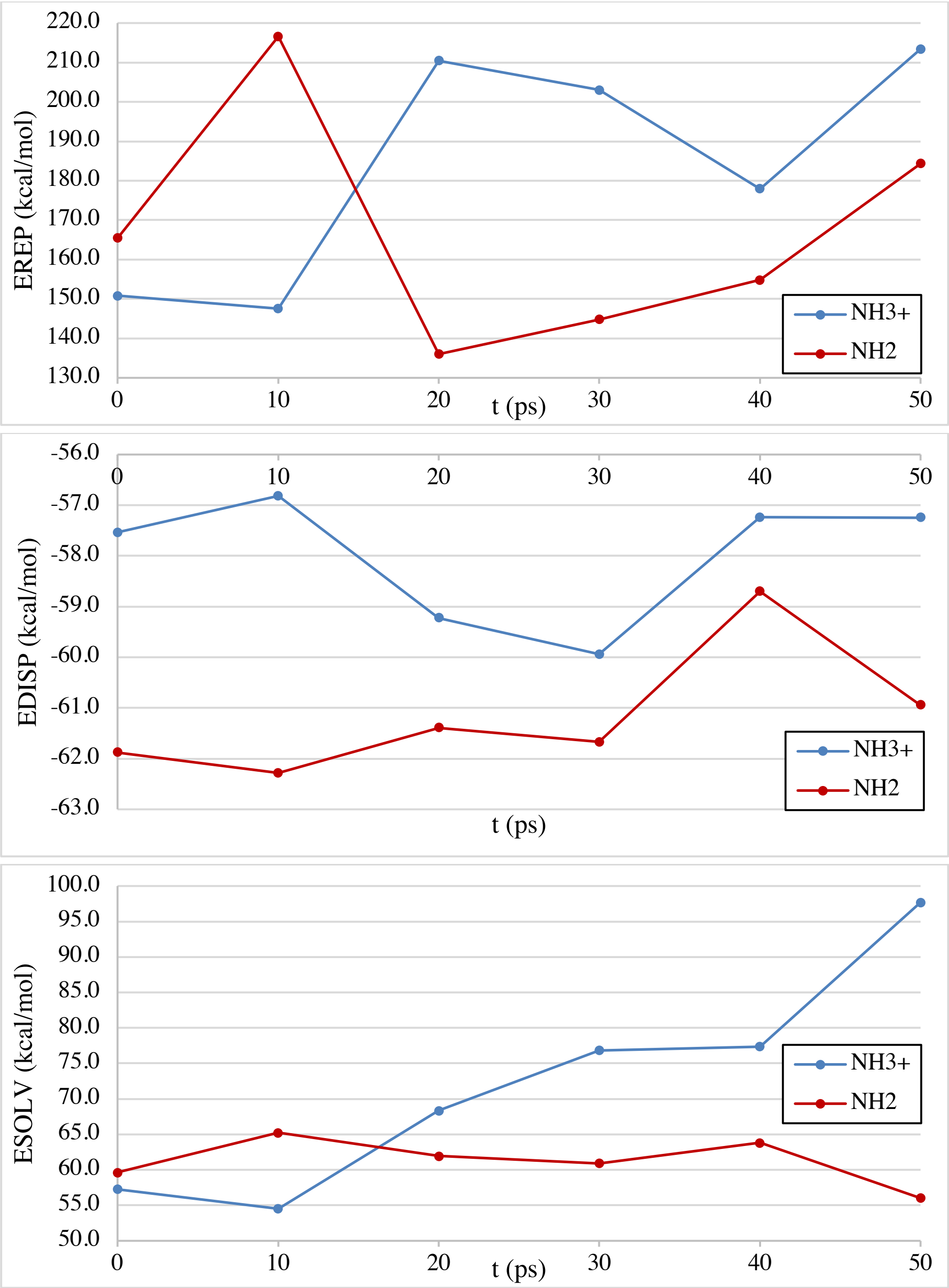


**Figure S16.7. Interaction maps for 5N8V (3.513) without water nor counter ions (left), with water but without counter ions (middle), and with water and with counter ions (right). For clarity, only 2 of the 59 explicit waters are represented in the picture.**

### S17 – Thermodynamic Cycle.

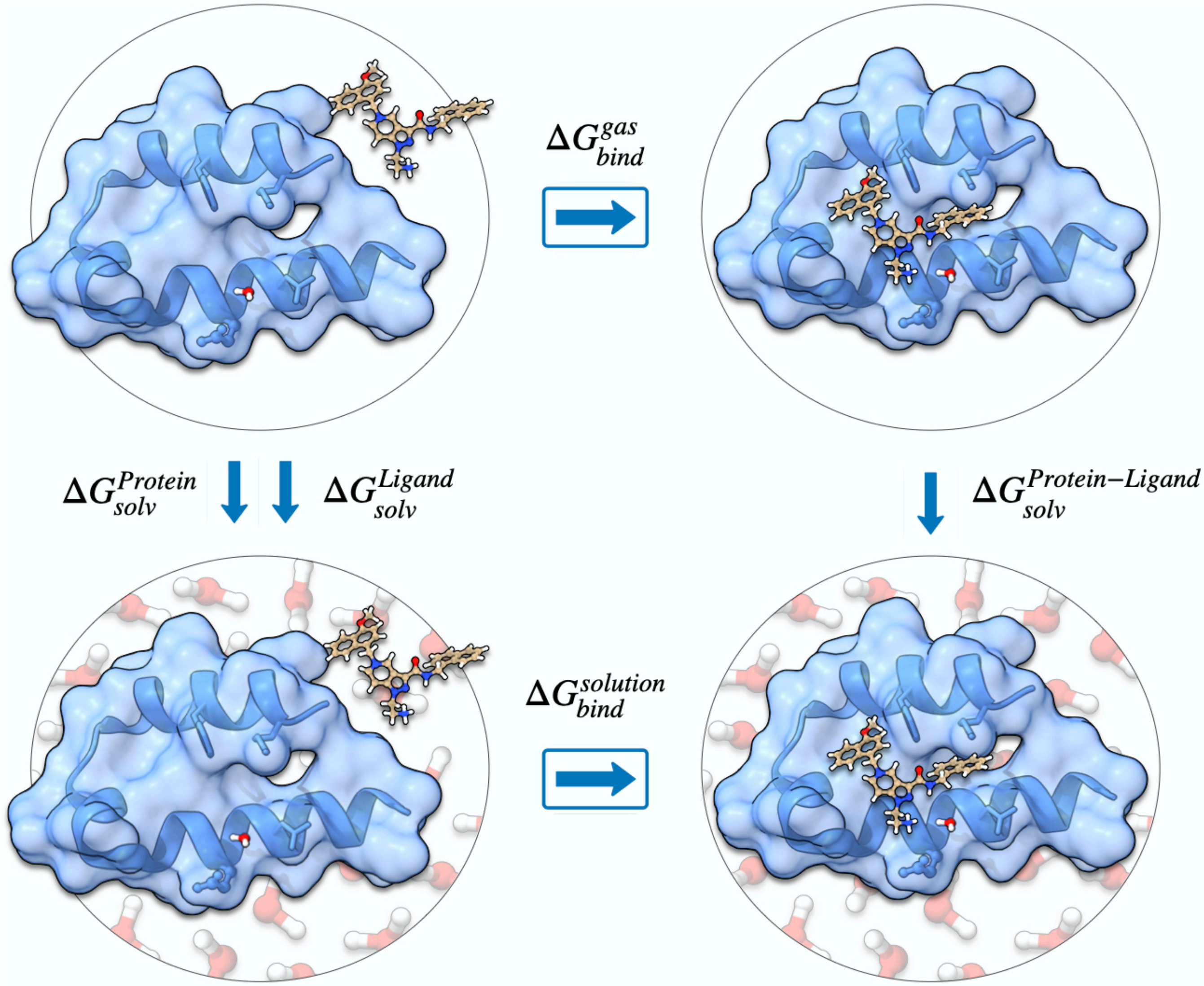


**Figure S17.1. Thermodynamic cycle on which the EDA calculation is based on.**

### S18 – Protein-Protein Interactions in the Crystal.

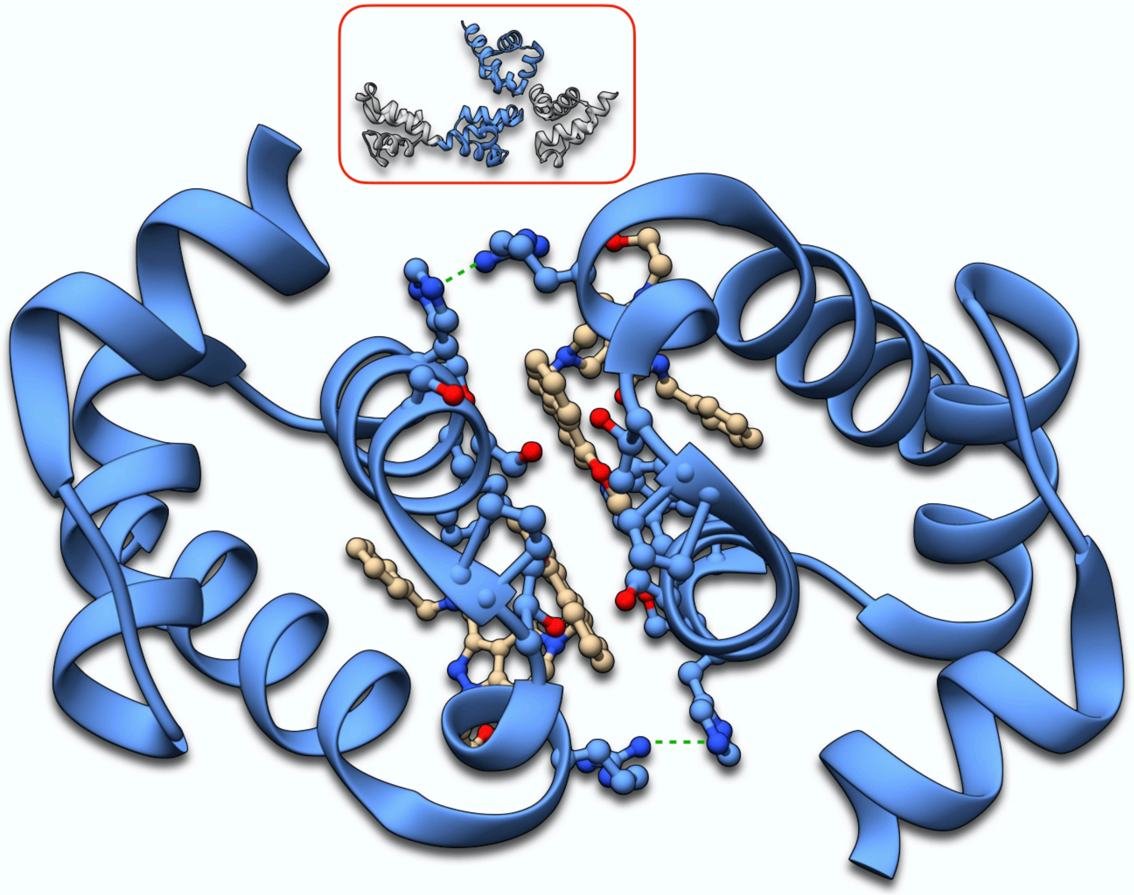

**Figure S18.1. Dimer 1 for 5L8A. Inhibitor (brown) included in the calculations. On the scheme on top the protein molecules composing the dimer are marked in blue. Spectator protein molecules (excluded from the calculation) are in grey.**

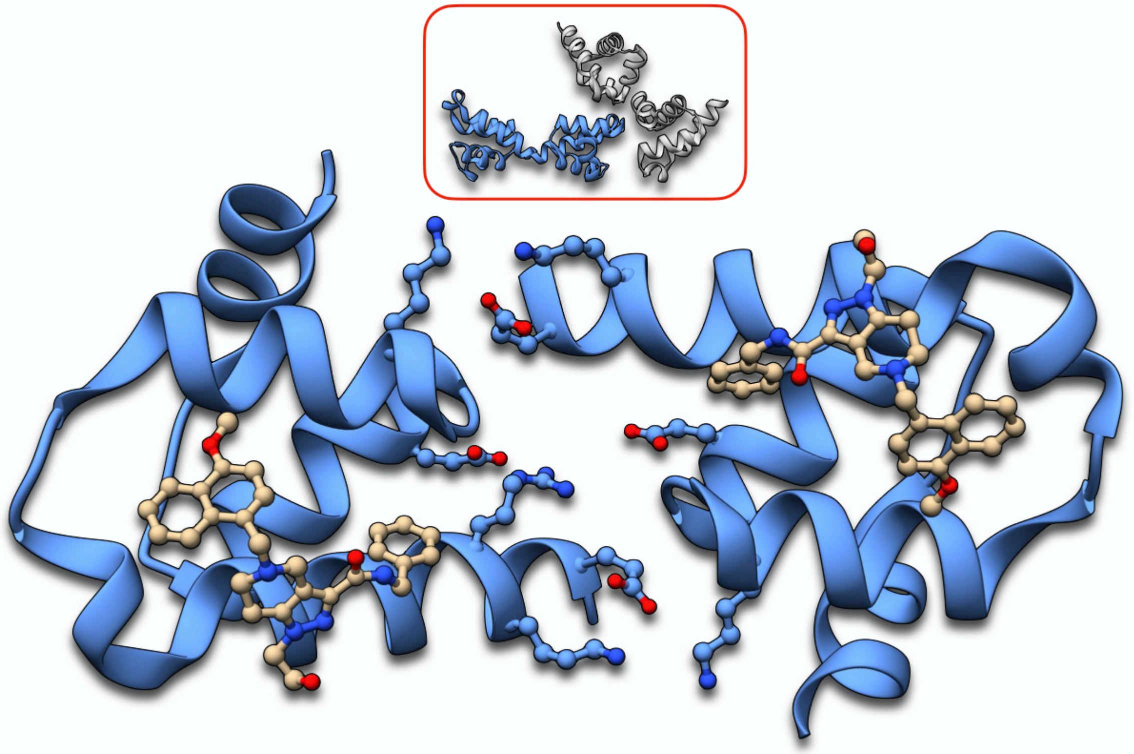

**Figure S18.2. Dimer 2 for 5L8A. Inhibitor (brown) included in the calculations. On the scheme on top the protein molecules composing the dimer are marked in blue. Spectator protein molecules (excluded from the calculation) are in grey.**

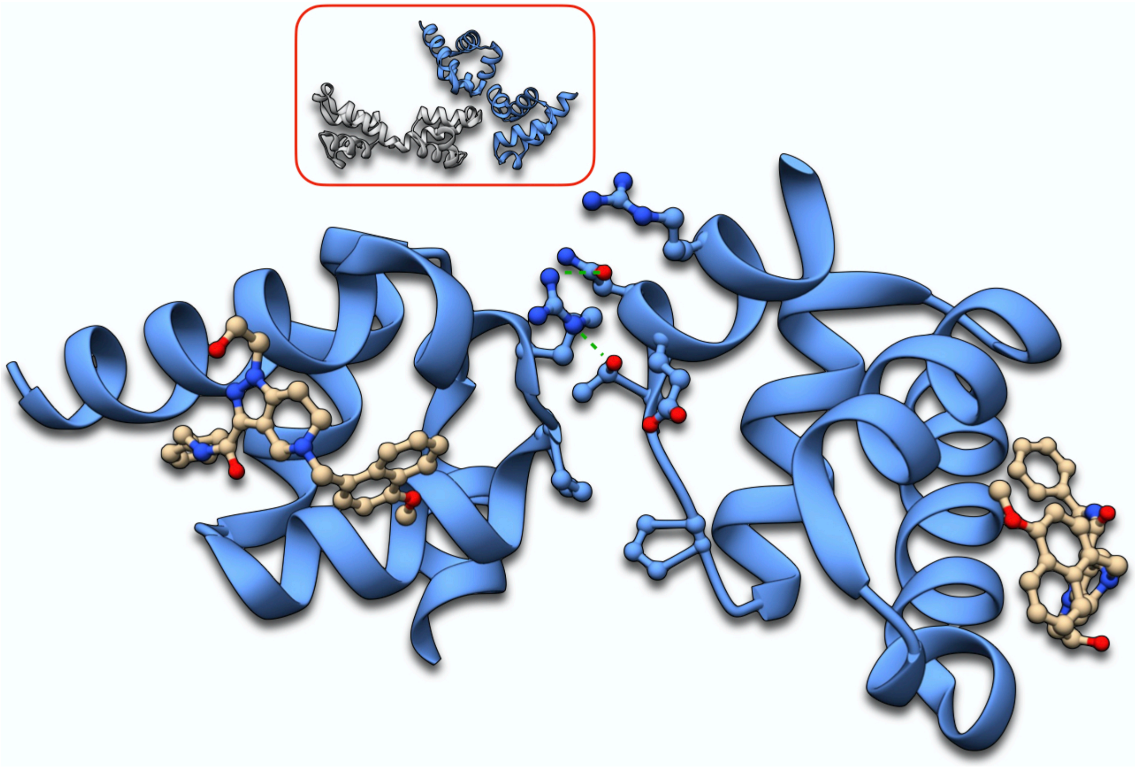

**Figure S18.3. Dimer 3 for 5L8A. Inhibitor (brown) included in the calculations. On the scheme on top the protein molecules composing the dimer are marked in blue. Spectator protein molecules (excluded from the calculation) are in grey.**

**Table S18.1. EDA data for Dimers of 5L8A.**

| | Dimer | | |
|---|---|---|---|
| | 1 | 2 | 3 |
| $E_{ES}$ | 291.84 | 328.32 | 338.52 |
| $E_{POL}$ | -13.16 | 10.52 | 30.66 |
| $E_{CT}$ | 0.00 | -0.29 | -0.01 |
| $E_{REP}$ | 50.57 | 14.90 | 12.88 |
| $E_{DISP}$ | -33.21 | -17.19 | -15.84 |
| $E_{SOLV}$ | -358.11 | -361.97 | -394.49 |
| $E_{INT}$ | -62.08 | -25.71 | -28.28 |

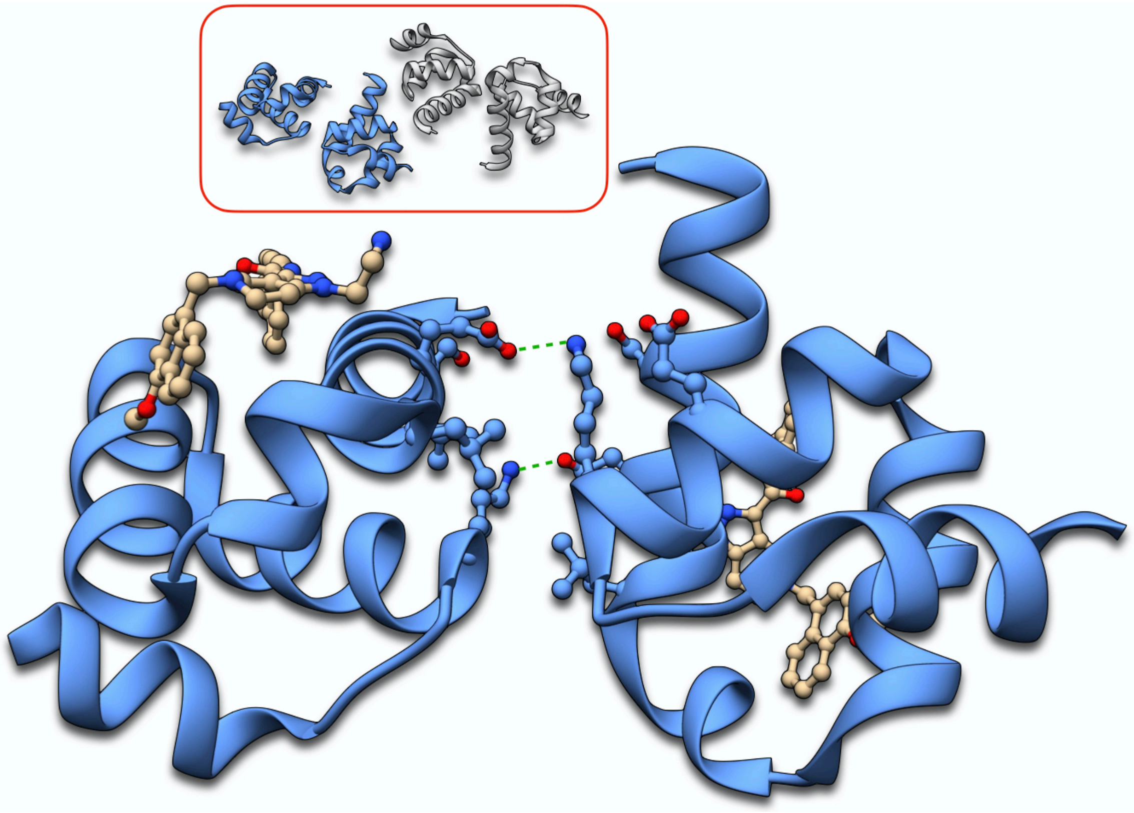

**Figure S18.4. Dimer 1 for 5N8V. Inhibitor (brown) included in the calculations. On the scheme on top the protein molecules composing the dimer are marked in blue. Spectator protein molecules (excluded from the calculation) are in grey.**

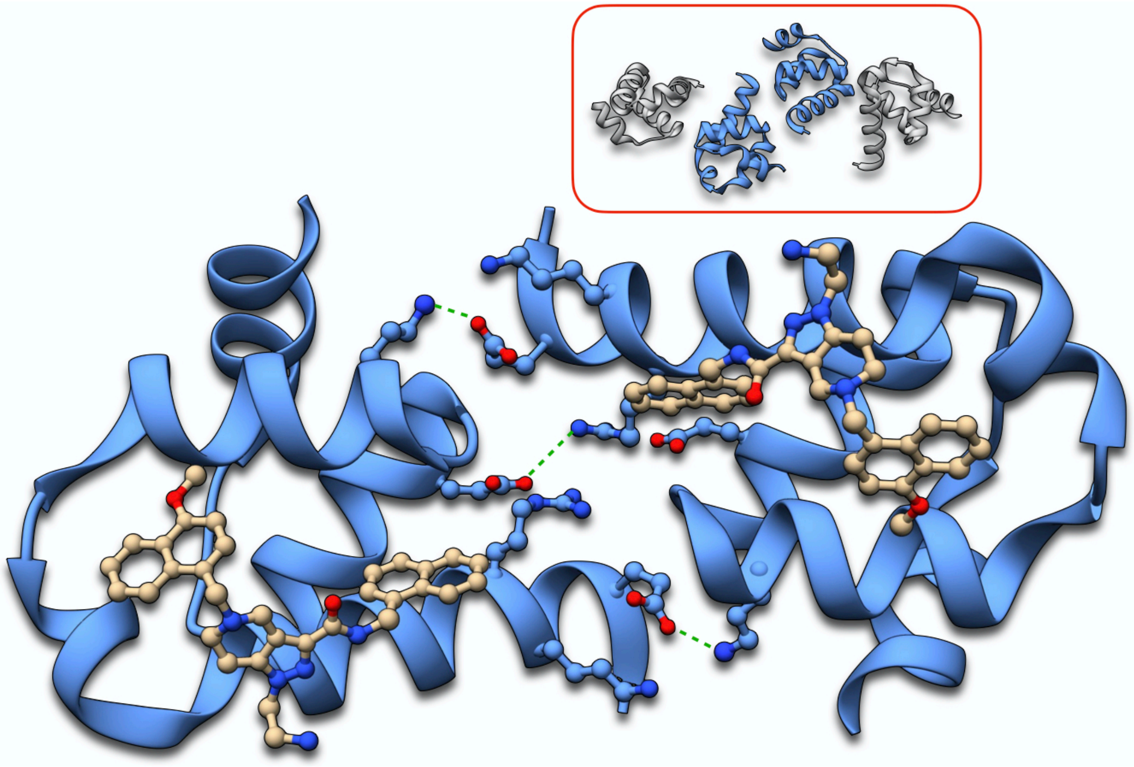

**Figure S18.5. Dimer 2 for 5N8V. Inhibitor (brown) included in the calculations. On the scheme on top the protein molecules composing the dimer are marked in blue. Spectator protein molecules (excluded from the calculation) are in grey.**

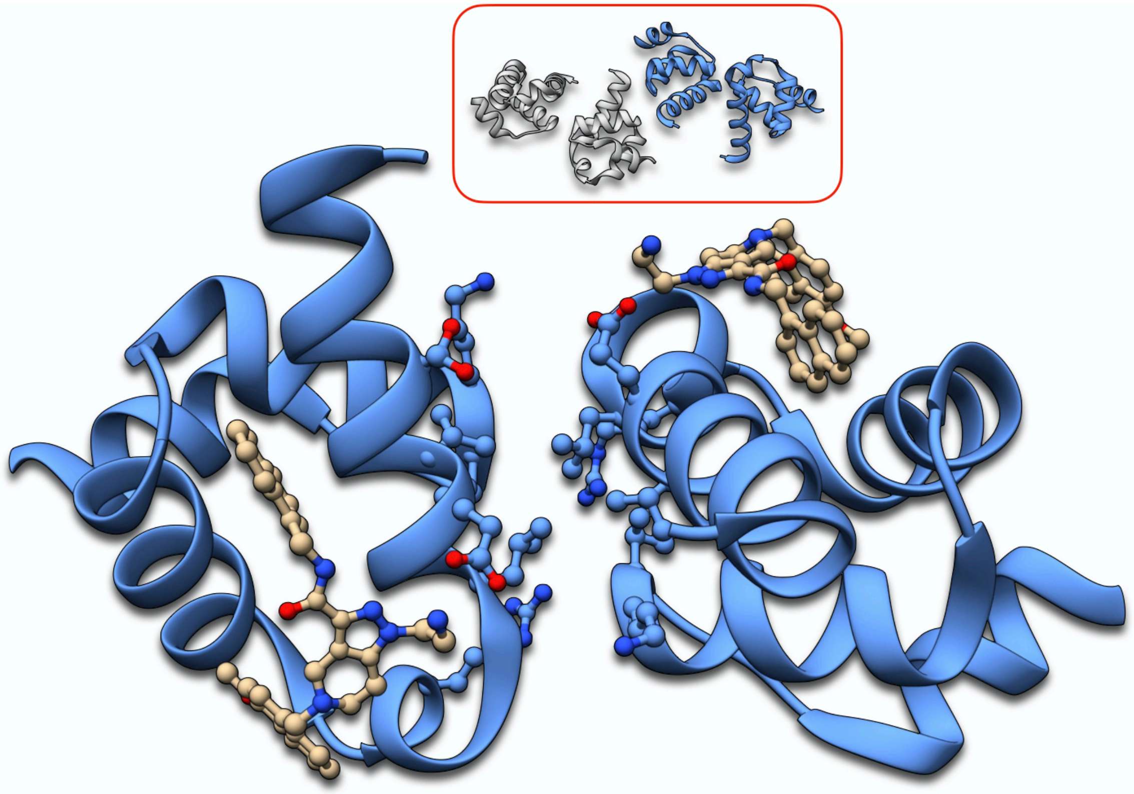

**Figure S18.6. Dimer 3 for 5N8V. Inhibitor (brown) included in the calculations. On the scheme on top the protein molecules composing the dimer are marked in blue. Spectator protein molecules (excluded from the calculation) are in grey.**

**Table S18.2. EDA data for Dimers of 5N8V.**

| | Dimer | | |
|---|---|---|---|
| | 1 | 2 | 3 |
| $E_{ES}$ | 178.50 | 291.17 | 246.95 |
| $E_{POL}$ | -28.63 | -10.22 | -14.63 |
| $E_{CT}$ | -0.44 | -1.15 | -0.02 |
| $E_{REP}$ | 22.10 | 25.10 | 14.90 |
| $E_{DISP}$ | -17.27 | -20.73 | -14.14 |
| $E_{SOLV}$ | -189.87 | -308.87 | -257.89 |
| $E_{INT}$ | -35.61 | -24.69 | -24.83 |

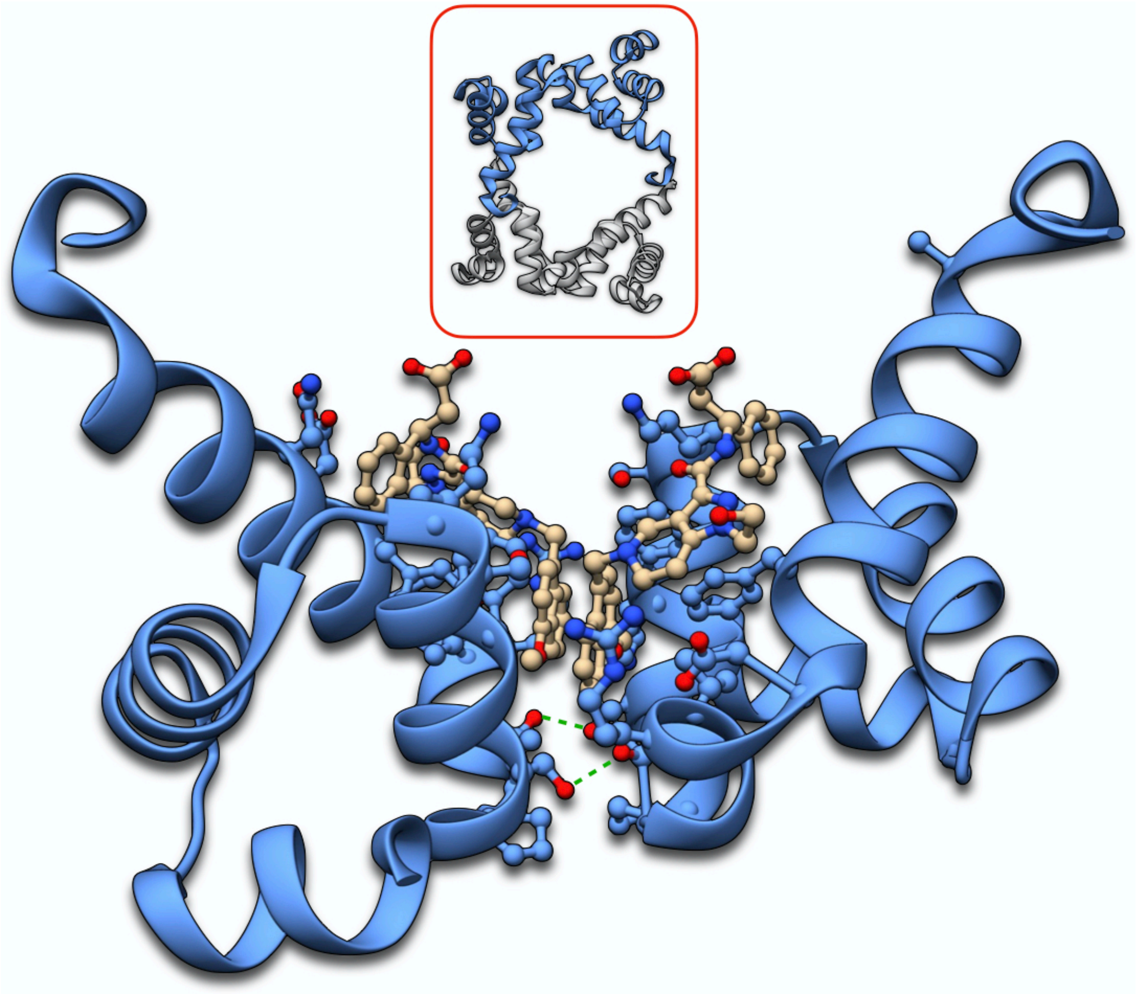

**Figure S18.7. Dimer 1 for 5OML. Inhibitor (brown) included in the calculations. On the scheme on top the protein molecules composing the dimer are marked in blue. Spectator protein molecules (excluded from the calculation) are in grey.**

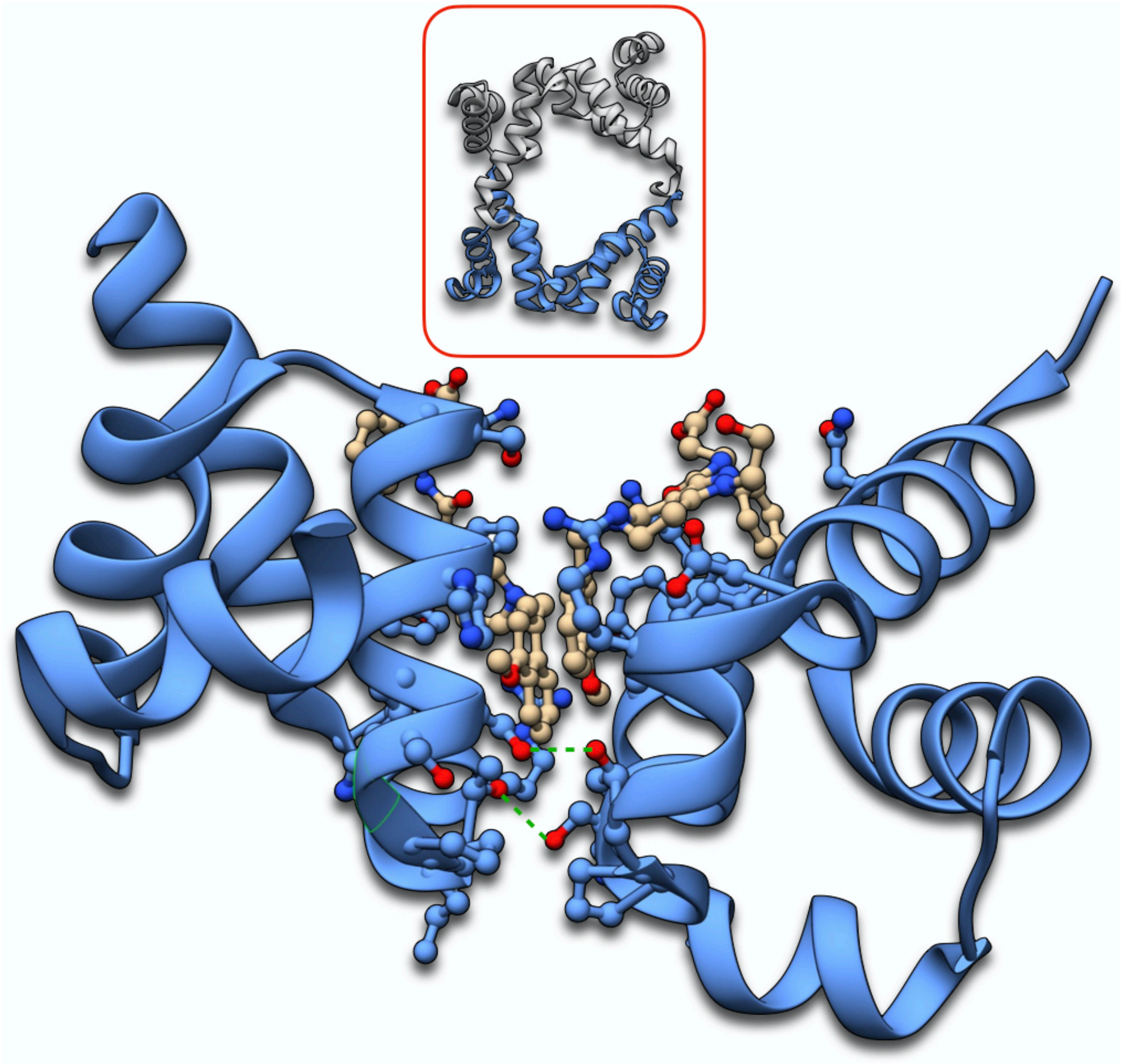

**Figure S18.8. Dimer 2 for 5OML. Inhibitor (brown) included in the calculations. On the scheme on top the protein molecules composing the dimer are marked in blue. Spectator protein molecules (excluded from the calculation) are in grey.**

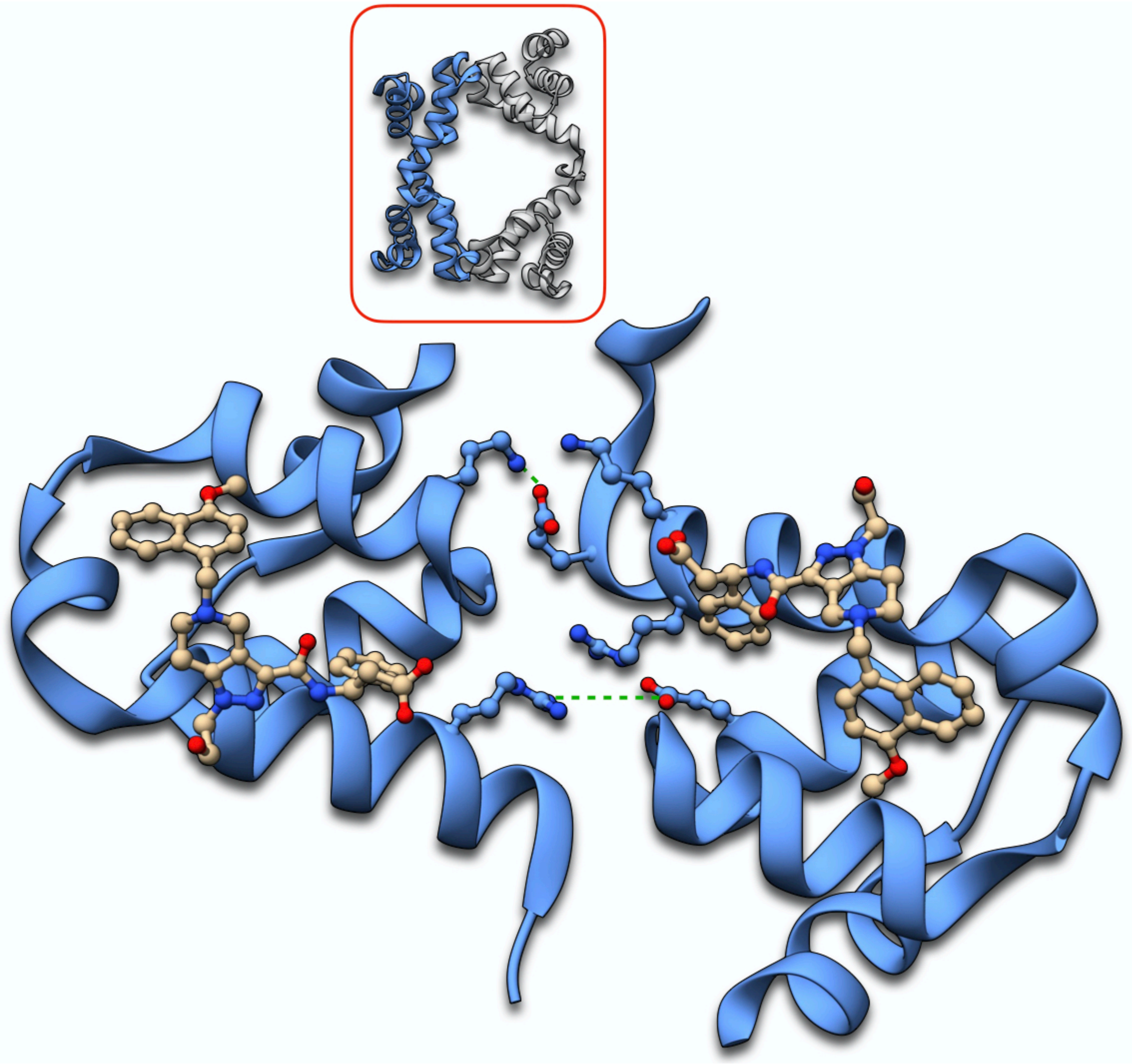

**Figure S18.9. Dimer 3 for 5OML. Inhibitor (brown) included in the calculations. On the scheme on top the protein molecules composing the dimer are marked in blue. Spectator protein molecules (excluded from the calculation) are in grey.**

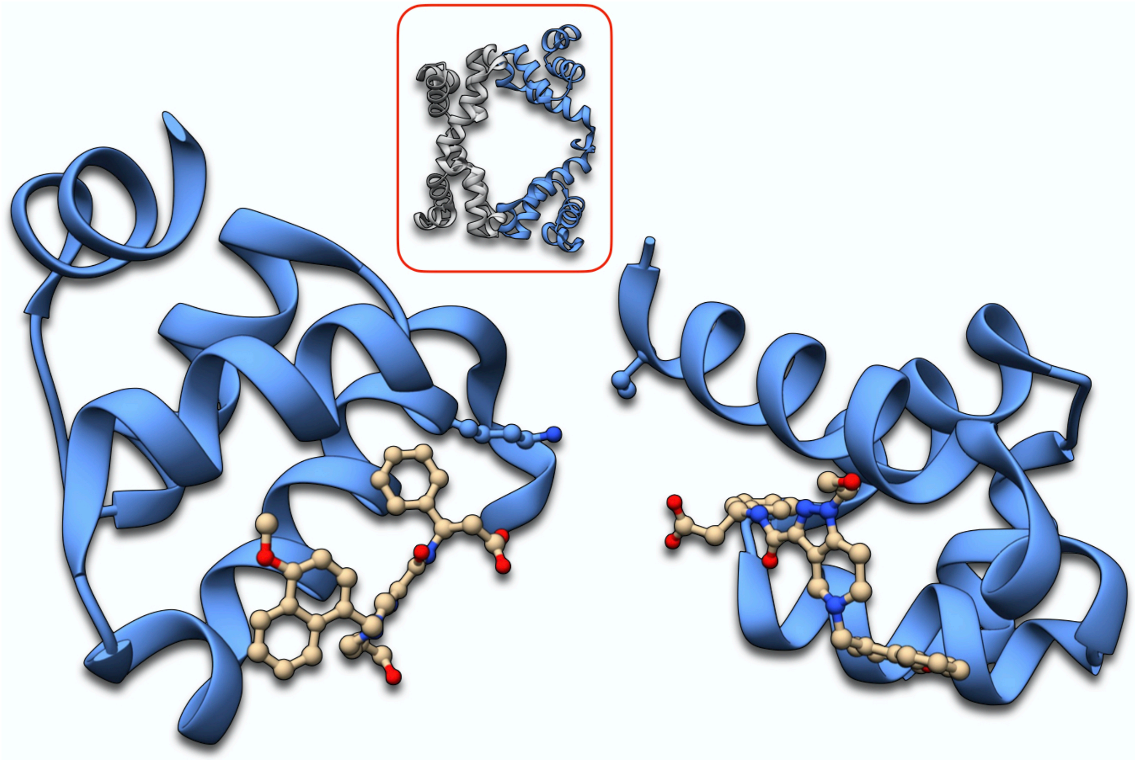

**Figure S18.10. Dimer 4 for 5OML. Inhibitor (brown) included in the calculations. On the scheme on top the protein molecules composing the dimer are marked in blue. Spectator protein molecules (excluded from the calculation) are in grey.**

**Table S18.3. EDA data for Dimers of 5OML.**

| | Dimer | | | |
|---|---|---|---|---|
| | 1 | 2 | 3 | 4 |
| $E_{ES}$ | 151.47 | 159.28 | 209.45 | 222.80 |
| $E_{POL}$ | 3.96 | -1.54 | -25.85 | 6.57 |
| $E_{CT}$ | 0.00 | -0.01 | -0.01 | -0.62 |
| $E_{REP}$ | 24.20 | 20.47 | 29.79 | 3.46 |
| $E_{DISP}$ | -31.62 | -30.69 | -20.39 | -4.14 |
| $E_{SOLV}$ | -202.17 | -198.88 | -229.52 | -240.26 |
| $E_{INT}$ | -54.16 | -51.36 | -36.52 | -12.19 |

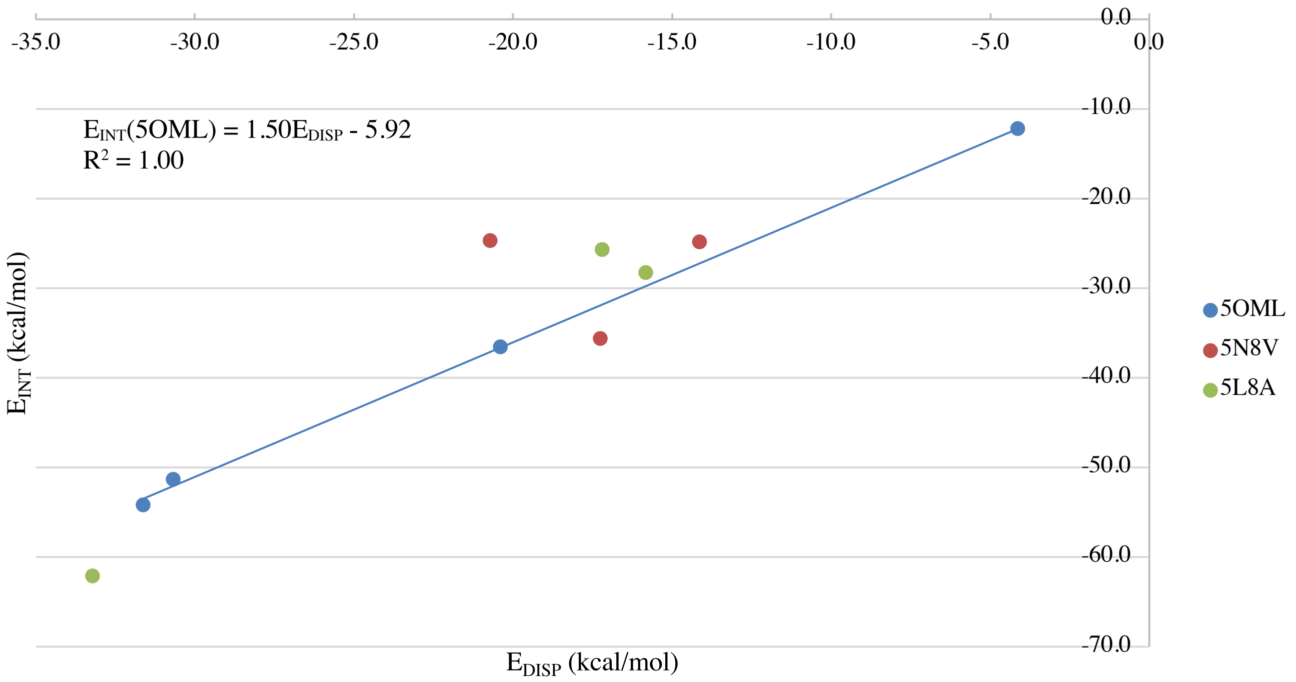


**Figure S18.11. Interaction energy Vs. the dispersion energy of the protein dimers in the crystal.**

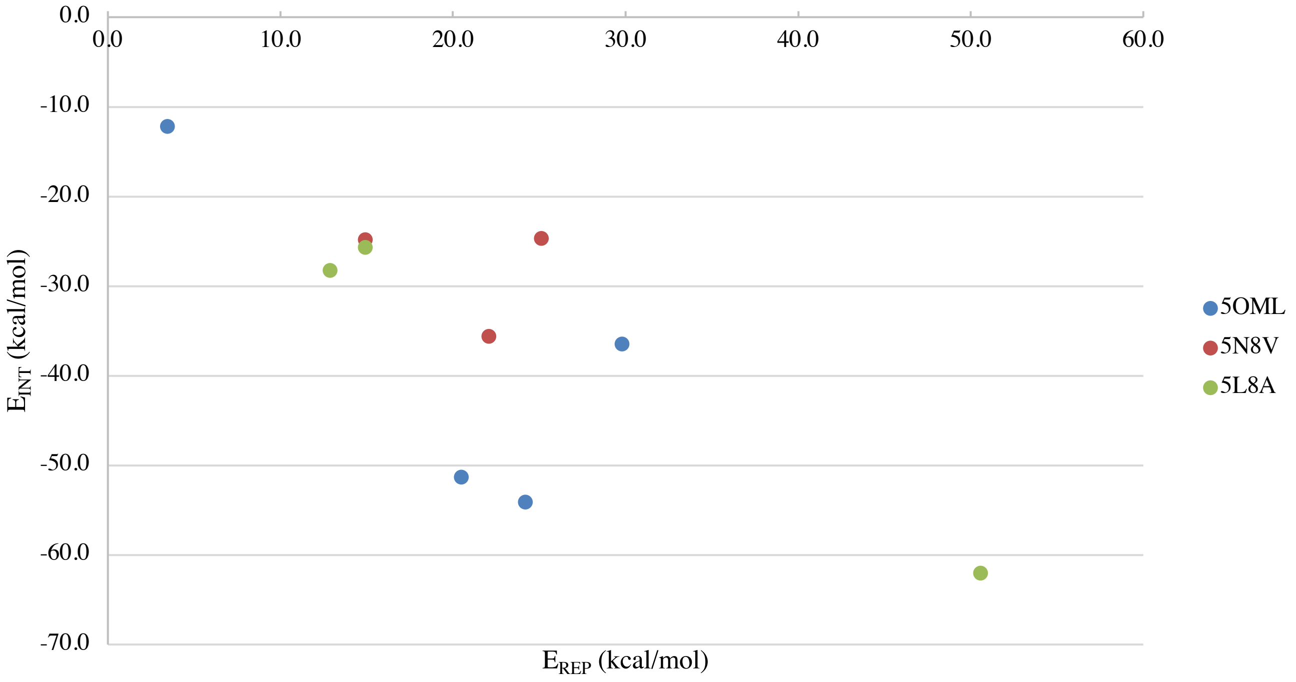


**Figure S18.12. Interaction energy Vs. the electronic density repulsion energy of the protein dimers in the crystal.**

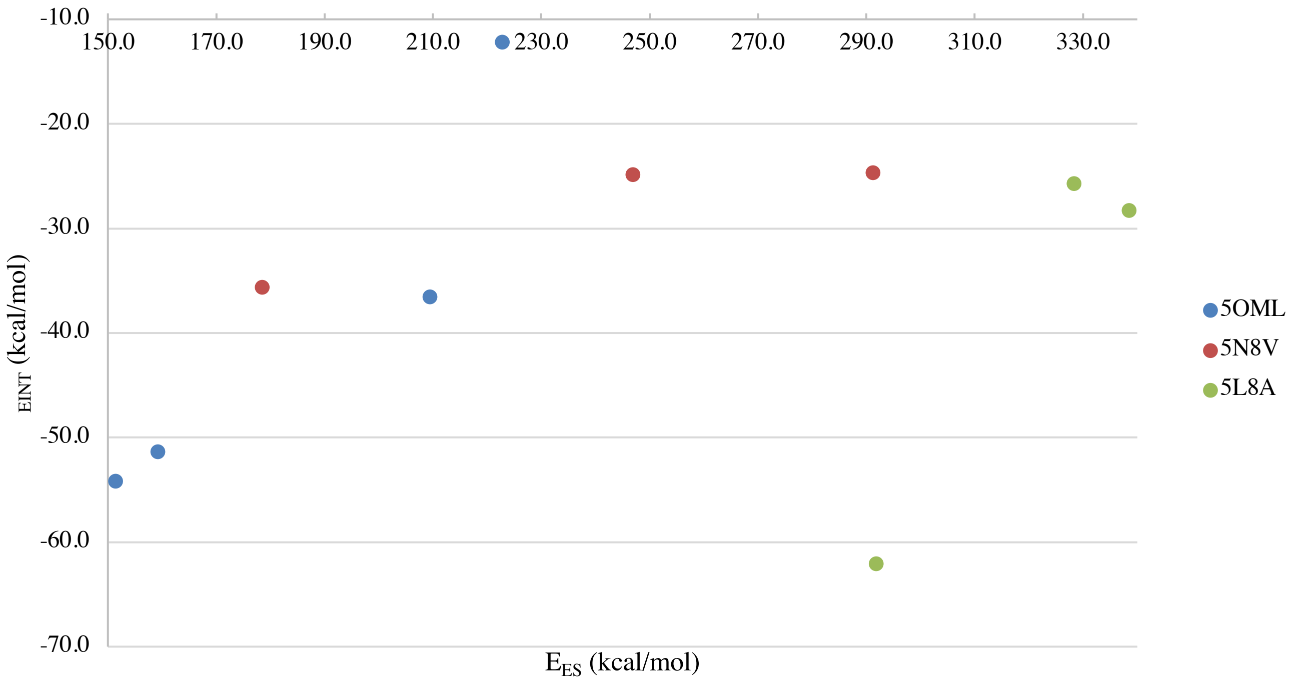


**Figure S18.13. Interaction energy Vs. the electrostatic energy of the protein dimers in the crystal.**

In all cases here analyzed, water molecules and counter-ions were excluded to avoid ambiguities in the assignment to a certain protein-ligand complex. Explicit water and counter-ions should however be needed for more accurate absolute electrostatics, solvation, and total binding.

We note that in all cases the interaction energy for all these dimers is negative (favorable). This might seem somewhat surprising from the electrostatics viewpoint, as each protein-ligand complex used in the calculations carries charge between +4 and +6. On the other hand, it also indicates that the driving force for crystallization is already recovered from the protein only.

### S19 – Comparative Dynamics of 5OML and 6RT2.

Dynamical simulations were run using OPLS-AA[61,62] and GROMACS.[63-70] Systems were handled and processed according to standard recommendations in the literature.[71] We note that since some of the ligands do not have zero net total charges, we systematically used AM1 charges for all simulations. The systems were equilibrated in NVT and then NPT ensembles for 1 ns each, followed by a 50 ns long simulation. EDA calculations were performed at every 10 ns of simulation time. The results are presented below.

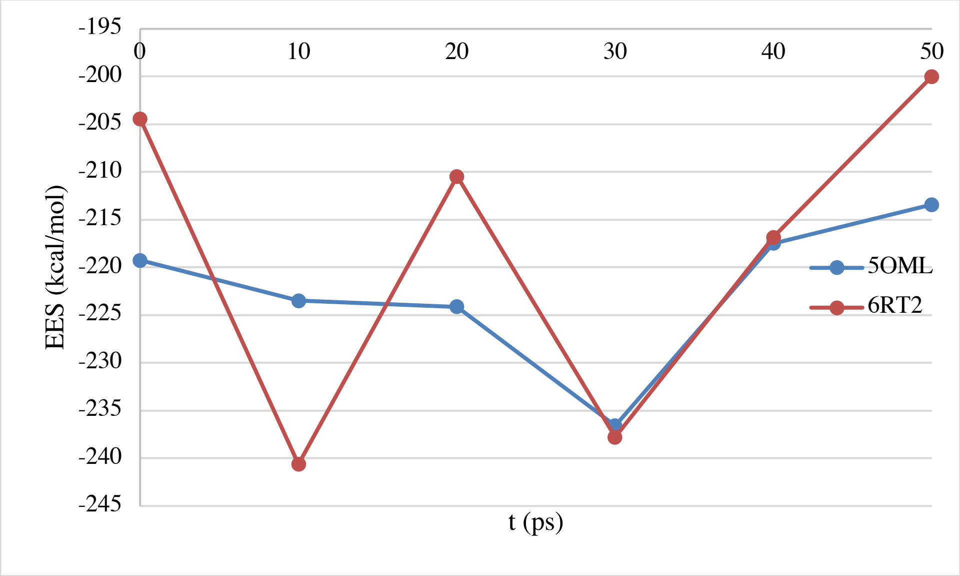
-195
-200
-205
-210
-215
-220
-225
-230
-235
-240
-245
0
10
20
30
40
50
EES (kcal/mol)
t (ps)
5OML
6RT2

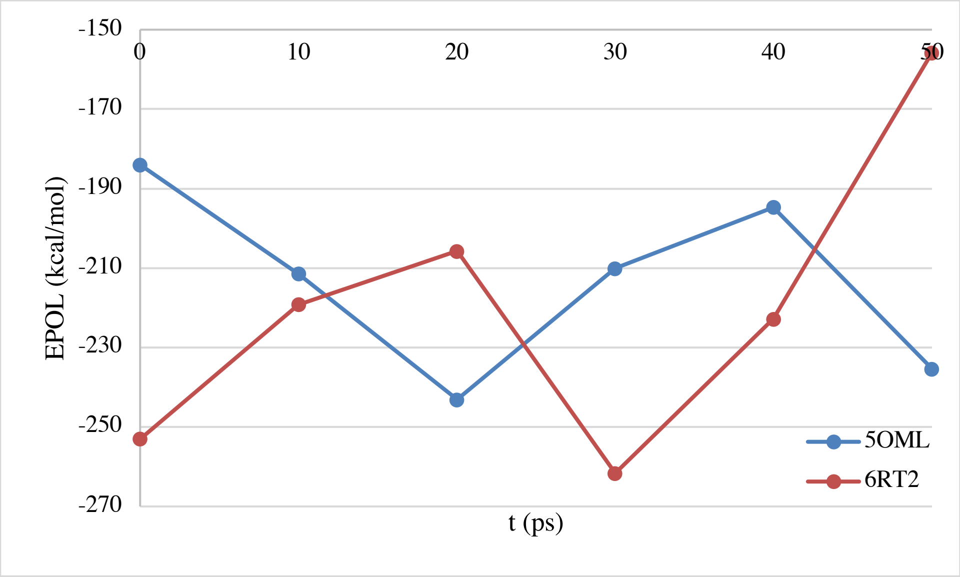
-150
-170
-190
-210
-230
-250
-270
0
10
20
30
40
50
EPOL (kcal/mol)
t (ps)
5OML
6RT2

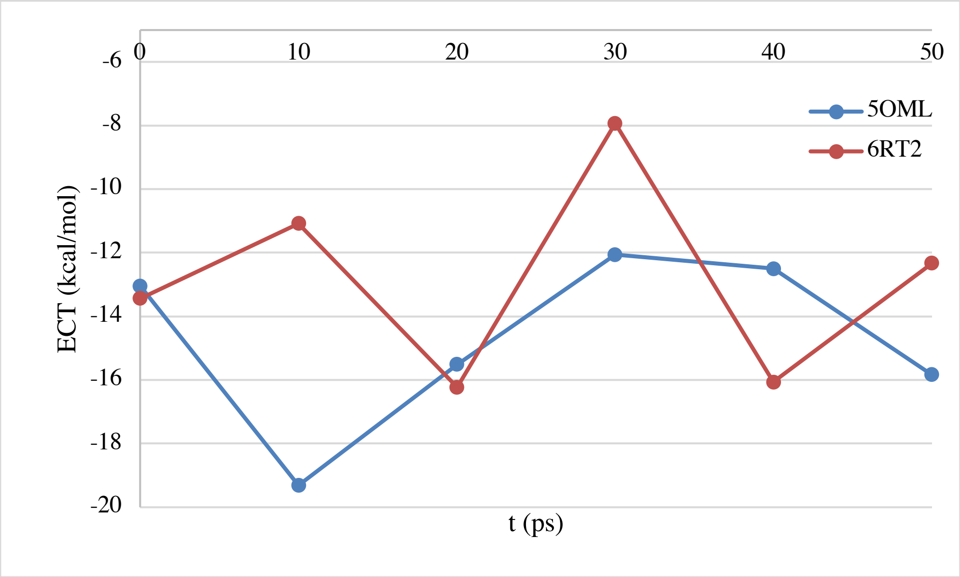
-6
-8
-10
-12
-14
-16
-18
-20
0
10
20
30
40
50
ECT (kcal/mol)
t (ps)
5OML
6RT2

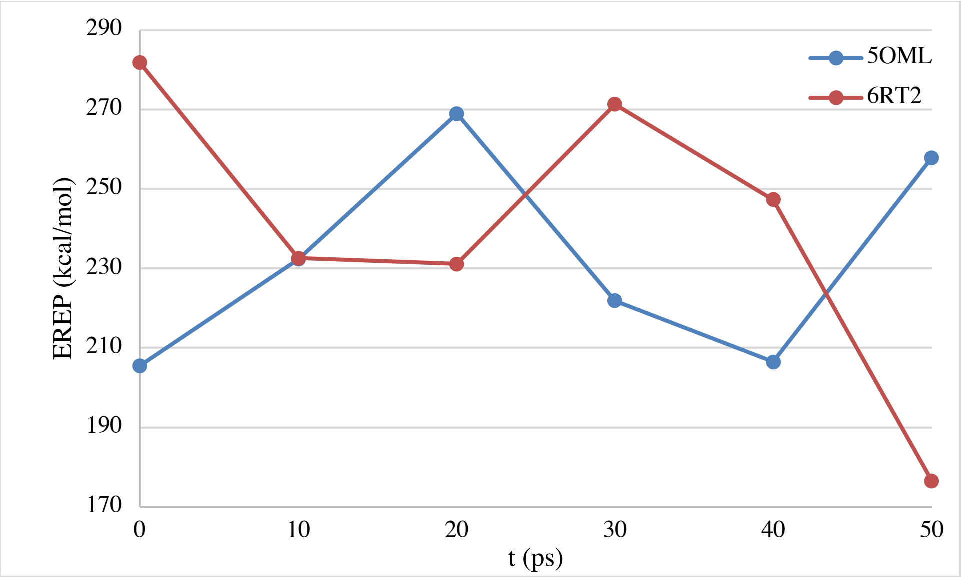
EREP (kcal/mol)
t (ps)
5OML
6RT2

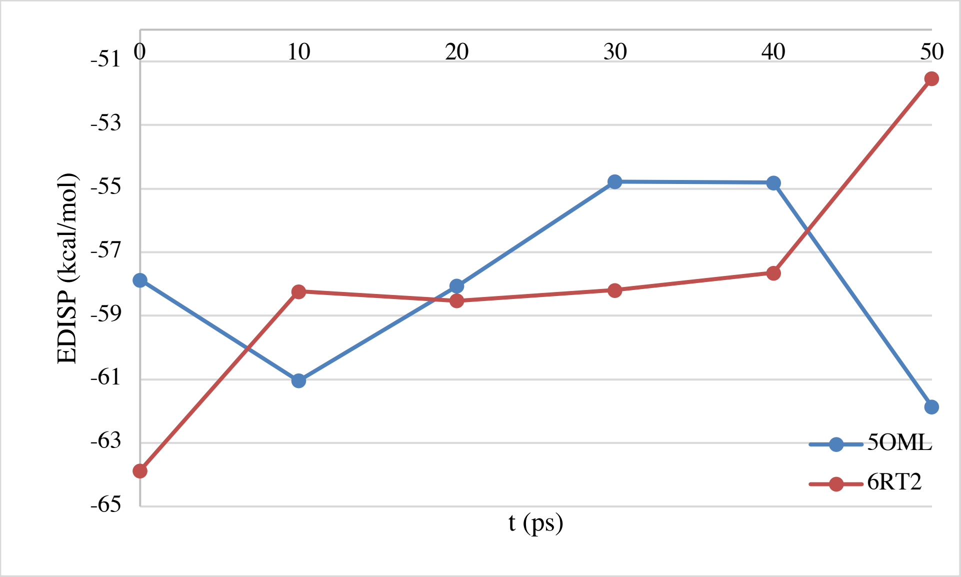
EDISP (kcal/mol)
t (ps)
5OML
6RT2

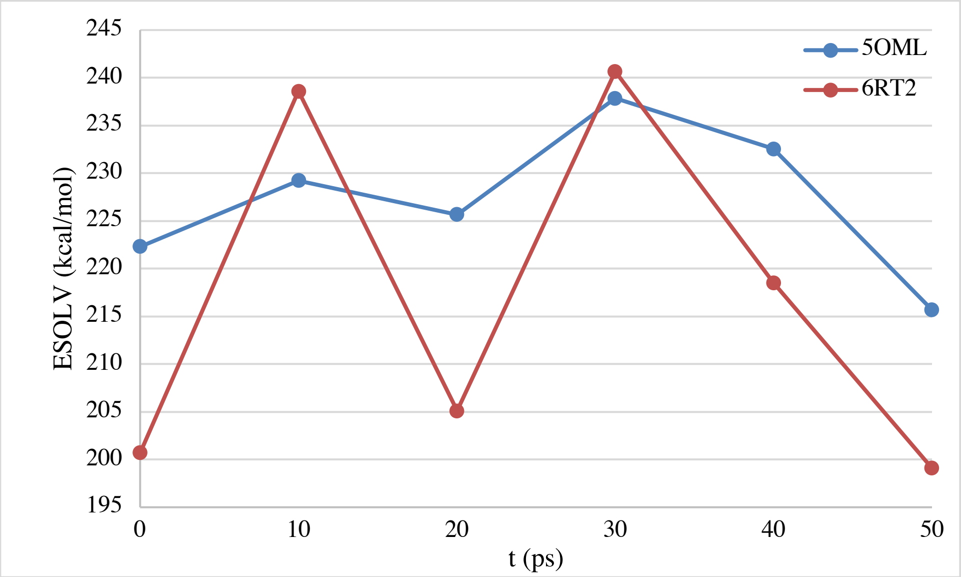
ESOLV (kcal/mol)
t (ps)
5OML
6RT2

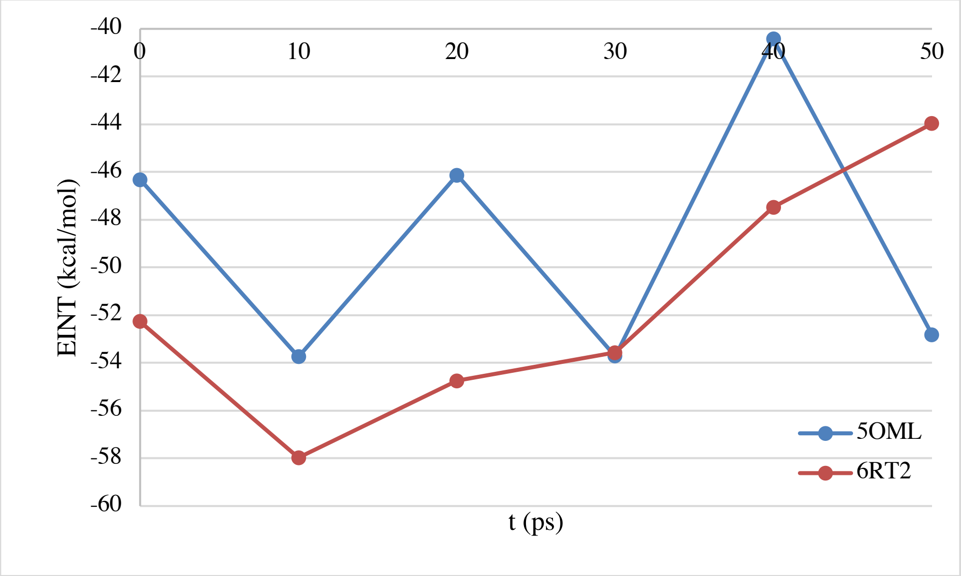
